# Site-specific O-glycosylation analysis of SARS-CoV-2 spike protein produced in insect and human cells

**DOI:** 10.1101/2021.02.03.429627

**Authors:** Ieva Bagdonaite, Andrew J. Thompson, Xiaoning Wang, Max Søgaard, Cyrielle Fougeroux, Martin Frank, Jolene K. Diedrich, John R. Yates, Ali Salanti, Sergey Y. Vakhrushev, James C. Paulson, Hans H. Wandall

## Abstract

Enveloped viruses hijack not only the host translation processes, but also its glycosylation machinery, and to a variable extent cover viral surface proteins with tolerogenic host-like structures. SARS-CoV-2 surface protein S presents as a trimer on the viral surface and is covered by a dense shield of N-linked glycans, and a few O-glycosites have been reported. The location of O-glycans is controlled by a large family of initiating enzymes with variable expression in cells and tissues and hence difficult to predict. Here, we used our well-established O-glycoproteomic workflows to map the precise positions of O-linked glycosylation sites on three different entities of protein S – insect cell or human cell-produced ectodomains, or insect cell derived receptor binding domain (RBD). In total 25 O-glycosites were identified, with similar patterns in the two ectodomains of different cell origin, and a distinct pattern of the monomeric RBD. Strikingly, 16 out of 25 O-glycosites were located within three amino acids from known N-glycosites. However, O-glycosylation was primarily found on peptides that were unoccupied by N-glycans, and otherwise had low overall occupancy. This suggests possible complementary functions of O-glycans in immune shielding and negligible effects of O-glycosylation on subunit vaccine design for SARS-CoV-2.

## Introduction

SARS-CoV-2 is a newly emerged zoonotic single-stranded RNA virus of the *Coronaviridae* family and the causative agent of the COVID-19 pandemic, the scale and impact of which is unprecedented in modern times. Genetic analysis of the spike protein (S) of SARS-CoV-2 shows that it has acquired a few features not found among previously emerged human coronaviruses [1]. This involves mutation of the receptor binding domain achieving high affinity binding to the human ACE2 receptor, as well as insertion of a functional polybasic furin cleavage site between the two subunits of the spike protein [2]. Proteolytic processing at the S1/S2 site is suggested to help adopt a favorable open conformation for ACE2 binding, and is speculated to be of value for transmissibility and pathogenicity [3–5].

Fast-acting measures are needed to relieve the global medical and economic burden imposed by the COVID-19 pandemic. Fortunately, several nucleic acid-based and inactivated virus vaccines have been rolled out at record speeds and vaccination programs have been initiated across the globe [6–9]. Many more vaccine clinical trials are underway, including subunit formulations, which rely on recombinant viral proteins for immunization [10]. An important element in the development of effective subunit vaccines is the characterization of the glycosylation of viral proteins. There are two major types of human glycosylation relevant for enveloped viruses – N-linked glycosylation, added co-translationally in the ER to N-X-S/T consensus sequons, and mucin type O-linked glycosylation, initiated in the Golgi by a family of polypeptide GalNAc transferases (GalNAc-Ts) modifying Ser, Thr, or Tyr. Different viruses have varying preferences for the two different types of glycans, with e.g. HIV-1, HCV and influenza virus having dense N-glycosylation on their surface proteins, whereas Ebola virus and herpesviruses additionally have many O-linked glycans [11]. Furthermore, both types of site-specific glycosylation have been shown to affect viral glycoprotein secretion or function [12]. Glycans represent structural features not encoded in the gene sequence, yet play a crucial role in immune recognition affecting vaccine designs. Such vaccine candidates are often expressed in cell lines that do not recapitulate the glycosylation pattern on native pathogens, and hence potentially do not elicit biologically relevant immune responses. Glycosylation of viral surface proteins is paramount for immune shielding, and altering positions of N-glycosylation sites by mutation of underlying acceptor amino acids is a well-established immune evasion strategy [11, 12]. A number of different protein formulations and expression systems have been used to deduce structure and post-translational modifications (PTMs) of SARS-CoV-2 spike protein (S), and some of those have been explored as candidates in vaccine research. So far, 22 highly occupied N-linked glycosylation sites have been identified on protein S, as well as a variable number of O-linked glycosylation sites at low stoichiometries [13–18].

Mucin type O-linked glycosylation is initiated by a competitive action of a family of 20 polypeptide GalNAc transferases (GalNAc-Ts) expressed in a tissue specific manner via the transfer of an N-acetylgalactosamine to Ser, Thr, and Tyr residues. It is difficult to predict such glycosylation events, and challenging to map specific locations of O-glycans [19, 20]. Collision induced dissociation (CID) based MS fragmentation methods often result in loss of the glycan modification on product fragment ions, which makes it difficult to map precise O-glycan positions on peptides with multiple Ser and Thr unambiguously. Thus, confident mapping of O-glycosites often require high performance instrumentation equipped with technology supporting electron transfer dissociation (ETD) fragmentation, which allows generation of glycan retaining peptide fragment ions. In some instances, optimized higher energy CID (HCD) methods can be sufficient, especially for simple O-glycan structures [21]. We have previously established paired ETciD and HCD workflows to enable robust peptide sequencing and high precision glycosite mapping and applied them for generating human and viral O-glycoproteomes [22–26]. Here, we aimed to compare the O-glycosylation patterns on different recombinant SARS-COV-2 S proteins expressed in insect or human cells, and investigate, whether O-glycosylation may have any implications on the choice of immunogens for vaccination studies. Using the described MS techniques, we mapped a total of 25 O-glycosites on three different subunit vaccine candidates expressed either in human or insect cells and estimated their occupancy, as well as location in the context of 3D S protein structure.

## Results

### Recombinant SARS-CoV-2 spike proteins are modified with multiple O-glycosylation sites

We first expressed pre-fusion stabilized S ectodomains in both *Drosophila* S2 or human embryonic kidney (HEK) 293F (Freestyle) cells using two slightly different constructs, both of which included a mutated S1/S2 cleavage site, as well as stabilizing mutations (986P, 987P) in the S2 region [18]. The human cell product featured a C-terminal leucine zipper (GCN4) motif, and primarily existed as a trimer, whereas the insect cell product did not contain a forced trimerization motif, and existed as several distinct entities, based on size exclusion chromatography, but predominantly as a dimer. Furthermore, we expressed a soluble monomeric receptor binding domain (RBD) in insect cells. We then mapped mucin type O-glycosylation sites on the recombinantly expressed S proteins using our well established ETciD and HCD nLC-MS/MS workflow on a Thermo Fisher Orbitrap Fusion Lumos Tribrid mass spectrometer. We applied an in-gel digestion approach with chymotrypsin or Glu-C followed by trypsin digestion in parallel to achieve deeper sequence coverage (Fig. S1). We pre-treated the gel-embedded proteins with PNGase F to remove the abundantly found N-glycans, and then searched for both unmodified and de-N-glycosylated peptides, where deamidation of asparagine yields an aspartic acid residue and a mass shift of +1 Da (Fig. 1A). The combined digestion strategy resulted in high sequence coverage, with more than 96 % of individual sequences identified. Chymotrypsin digestion yielded higher numbers of peptide groups as well as glycoPSMs (Dataset S1), though, suggesting either better separation or sensitivity of the individual runs (Fig. S2). In relation to neuraminidase treatment of peptides, we searched for simple GalNAcα-S/T (Tn) and Galβ3GalNAcα-S/T (core 1) structures abundantly found in both HEK 293F and S2 cell lines, where the core 1 structures are usually capped with sialic acid in human cells and unmodified in insects (Fig 1A).

**Figure 1.**
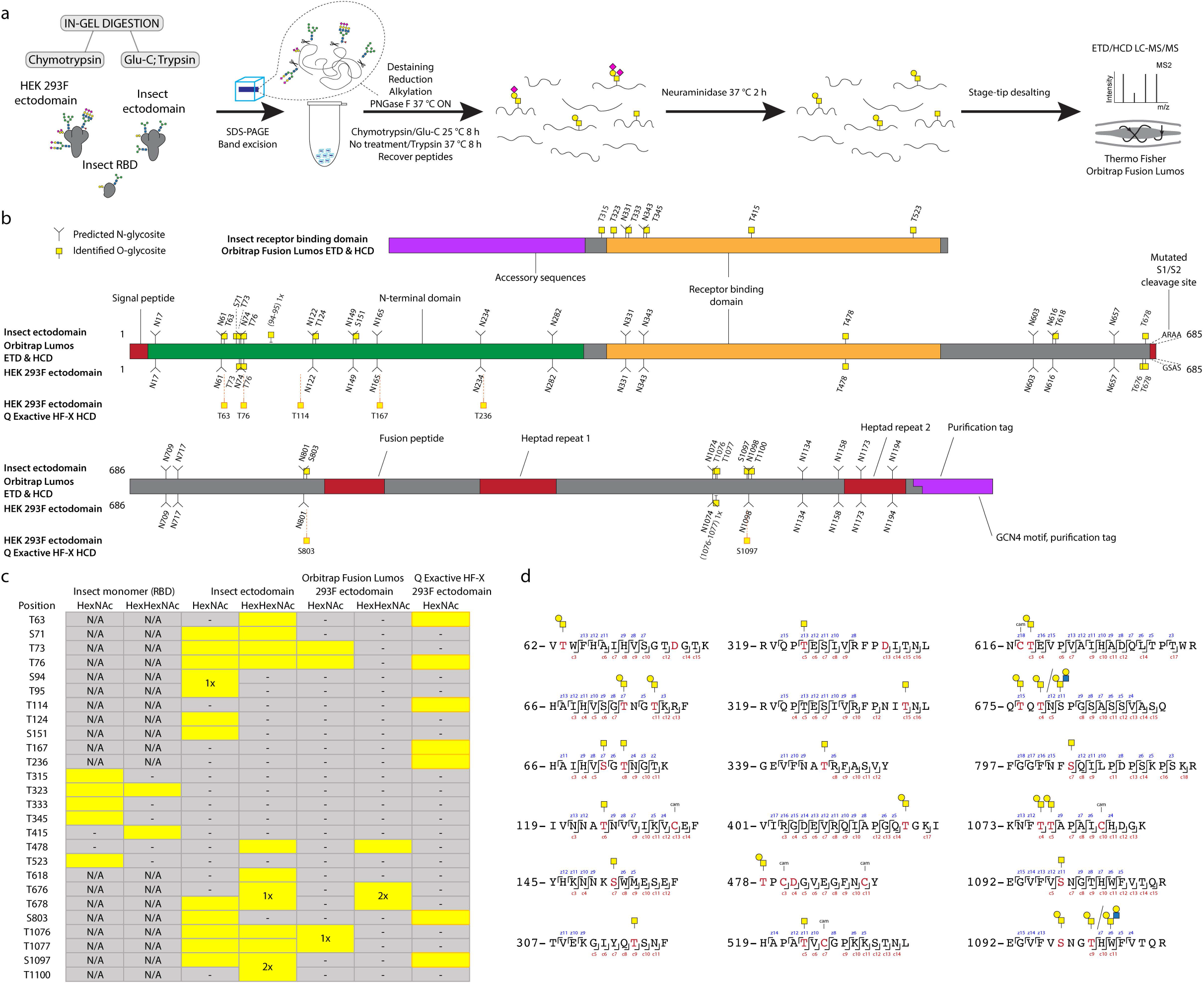
O-glycosylation of SARS-CoV-2 spike protein expressed in insect or human cells. (a) Experimental strategy. (b) A graphical layout of spike protein sequence annotated with identified O-glycosylation sites using Orbitrap Lumos mass spectrometer with ETD and HCD MS2 fragmentation. Independently identified O-glycosites on HEK 293F-derived ectodomain using Q Exactive HF-X mass spectrometer with HCD MS2 fragmentation are shown with orange outlines. (c) A table summarizing O-glycosites and respective structures found in different S formulations (yellow rectangles). Ambiguous sites are shown as merged rectangles across several positions. “N/A” - not applicable; “-” - not detected. Orbitrap Fusion Lumos derived sites are marked with grey outlines. Additional sites identified on HEK 293F ectodomain with Q Exactive HF-X are marked with orange outlines. (d) Examples of O-glycopeptides identified with Orbitrap Fusion Lumos using ETD fragmentation. MS2 c and z product ion fragments are annotated based on the respective ETD spectra (Fig S1).

A paired ETciD and HCD MS2 fragmentation strategy allowed sequencing of the peptide backbone with high confidence and identification of the precise positions of modified amino acids (Fig. 1 B-D). We identified 6, 15, and 6 O-glycosites on the insect RBD, insect ectodomain, and HEK 293F ectodomain, respectively. *Drosophila* S2 expression and purification system generated exceptionally clean products, likely resulting in the higher relative abundance and detection of O-glycopeptides (Fig. S1). In contrast, it is more difficult to achieve high purity in human cell expression systems, and despite excision of specific bands, we found a high number of human proteins in the HEK 293F-expressed S protein samples, resulting in increased sample complexity and reduced sensitivity of O-glycopeptide identification (Figs. S1, S2). We therefore applied additional analytical strategies for the HEK 293F product. We took advantage of a well-established N-glycoproteomic workflow [27] and modified it to include a search for O-GalNAc structures, leading to independent identification of 7 O-glycosites using a Thermo Fisher Q Exactive HF-X instrument equipped with HCD MS2 fragmentation (Datasets S2, S7). The combined results from the two different analyses yielded 10 total O-glycosites on the HEK 293F-expressed S protein, seven of which were analogous to those found on the insect cell expressed ectodomain (Fig. 1B,C).

### Differences in O-glycosylation patterns between multimeric and monomeric S

We obtained rather similar O-glycosylation patterns on the two multimeric ectodomains expressed in different cell lines. O-glycans were most abundantly found at the N-terminal domain, and close to the C-terminus (Fig. 1B). One O-glycosite (T478) was mapped to the RBD, and a few more were found in close proximity to other functionally relevant regions, such as the S1/S2 cleavage site (T676, T678), and the fusion peptide (S803). In contrast, there were no similarities between the RBD O-glycosylation patterns of insect monomer and dimer, as none of the six sites found on the monomer overlapped with the single RBD-localized site on the dimeric ectodomain. This suggests better accessibility to enzymes on the monomeric S protein RBD, which is taken out of the context of the presentation within full-length S protein scaffold, as well as the quaternary protein structure.

### Majority of O-glycosites are located in vicinity of N-glycosites

For Orbitrap Fusion Lumos derived data, we summed the integrated MS1 peak areas of relevant peptides covering individual O-glycosites or their clusters to estimate the abundances of O-glycosylated versus unmodified peptides. This revealed low occupancies at most of the sites (Fig. 2A, “All”, Datasets S3-S5), with less than 5 % of quantified peptides carrying O-glycans. The highest occupied O-glycosites were found next to N74, N616, and N801, as well as at T478. Strikingly, a large proportion of O-glycosites were found either within or just before N-X-S/T sequons of N-glycosylation. We wanted to understand, whether O-glycans could co-exist on the same peptide molecules as N-glycans. Therefore, we split up the N-X-S/T covering O-glycopeptides into two groups, those that had an asparagine in the sequence, and thus were not N-glycosylated, and those that instead contained an aspartic acid, suggestive of enzymatic deamidation after PNGase F treatment. Whilst spontaneous deamidation of Asn can happen, multiple reports of protein S N-glycosite analyses suggest very high occupancies of N-glycosylation and it is very likely that such peptides represent deglycosylated peptides. We then again estimated occupancy in the individual groups as proportion of sum MS1 peak area. What we observed was that most of the O-glycosites located next to N-X-S/T sequons were found on non-N-glycosylated peptides (Fig. 2A, Datasets S3-S5). Very few O-glycosites were identified on N-deamidated peptides, and only few non-N-glycosylated peptides did not contain O-glycans. For 7 out of 9 glycan clusters of interest, more than 85 % of quantification signals came from O-glycopeptides, if there were no N-glycans present (Fig. 2C). For Q Exactive HF-X derived data, generated using a combination of different protease-specific products per run, we grouped peptides of different length covering individual O-glycosites, calculated occupancy for each of them, and presented average occupancy (Fig. 2B, Dataset S6). In agreement with Orbitrap data, very low occupancies were estimated for commonly identified O-glycosites. The present strategy has also strengthened our conclusions regarding co-ocurrence of N- and O-linked glycans, since sequential de-glycosylation with Endo H followed by PNGase F in O^18^ water allows for discrimination of occupied N-glycosites prior to the treatment. None of the N+203 or N+3 peptides contained O-glycans, and O-glycans were only found on non-N-glycosylated peptides (Fig. 2C).

**Figure 2.**
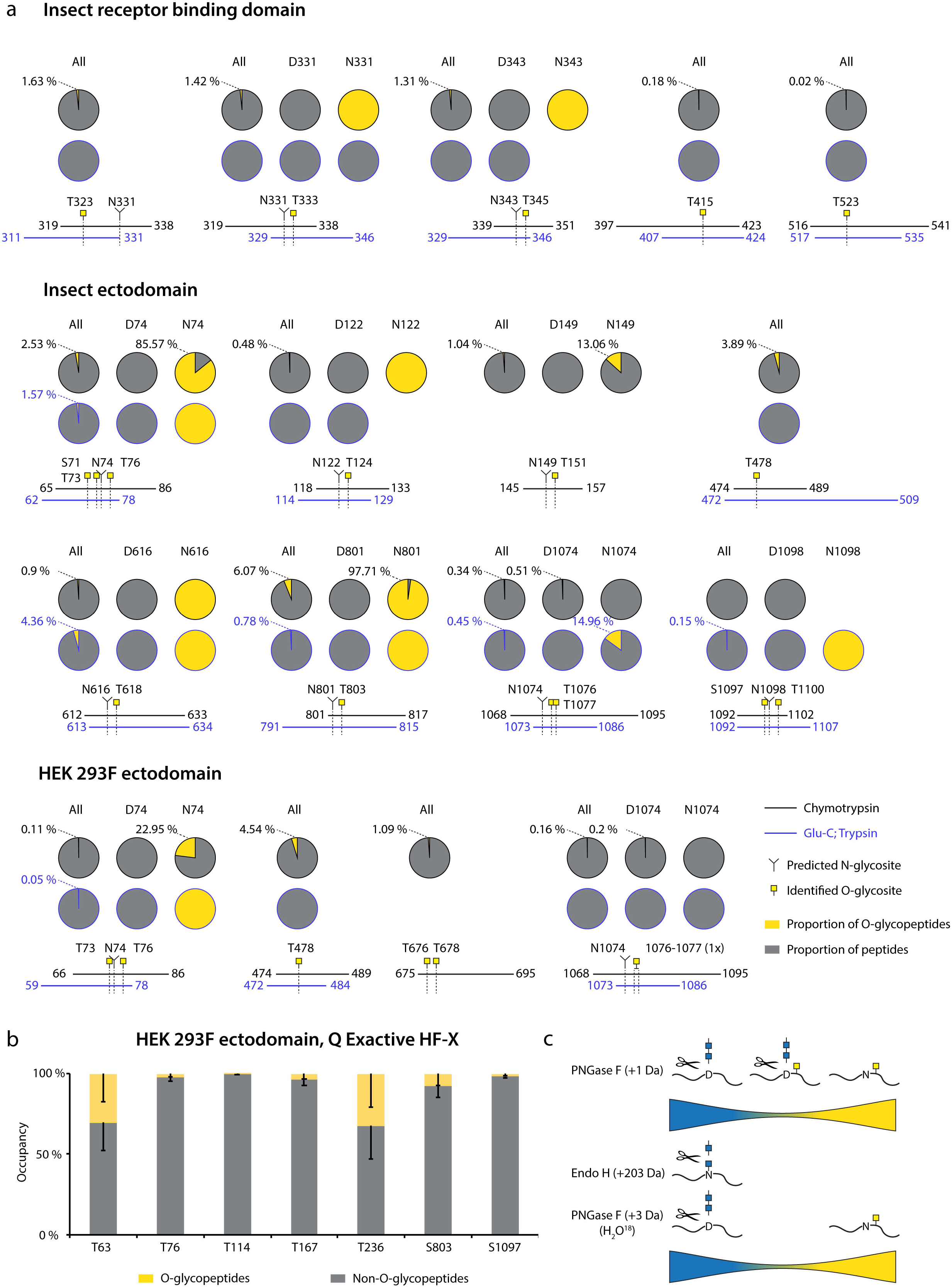
Estimation of O-glycosite occupancy. (a) Estimated occupancy of samples analyzed with Orbitrap Fusion Lumos mass spectrometer based on available quantification values of (glyco)peptides. Circle diagrams depict proportions of O-glycosylated (yellow) and non-O-glycosylated (grey) peptides based on integrated MS1 signals of relevant peptides. When in close vicinity to predicted N-glycosites, separate quantifications for non-N-glycosylated peptides (N(Asn)-peptides) and presumably N-deglycosylated deamidated peptides (D(Asp)-peptides) after PNGase F treatment are provided. Coverage by peptides is shown under each graph. (b) Peptides covering individual O-glycosites indentified with Q Exactive HF-X mass spectrometer were grouped based on length and occupancies based on integrated MS1 signals calculated for each group. Average occupancies of individual O-glycosites -SEM are shown. (c) The cartoon suggests a model, where the majority of identified O-glycopeptides are not N-glycosylated.

### O-glycosylation in the context of S protein structural features

We next aimed to map the identified O-glycosites onto the fully-glycosylated full-length S protein structure. Model 6vsb_1_1_1 was downloaded from http://www.charmm-gui.org/?doc=archive&lib=covid19 [28]. This model contains N-glycans in agreement with [16]; however, we modified the N-glycan structures to paucimannosidic structures for insect-derived products. We used Conformational Analysis Tools software (www.md-simulations.de/CAT/) interfaced with STRIDE [29] to estimate the solvent accessible surface (SAS) of the amino acids. To visualize the 3D context of O-glycosites Galβ3GalNAcα disaccharides were attached manually to the 3D model and optimized using YASARA [30] (Fig. 3A-C). The structures were trimmed to GalNAcα for visualization, if only such structures were identified by mass spectrometry.

**Figure 3.**
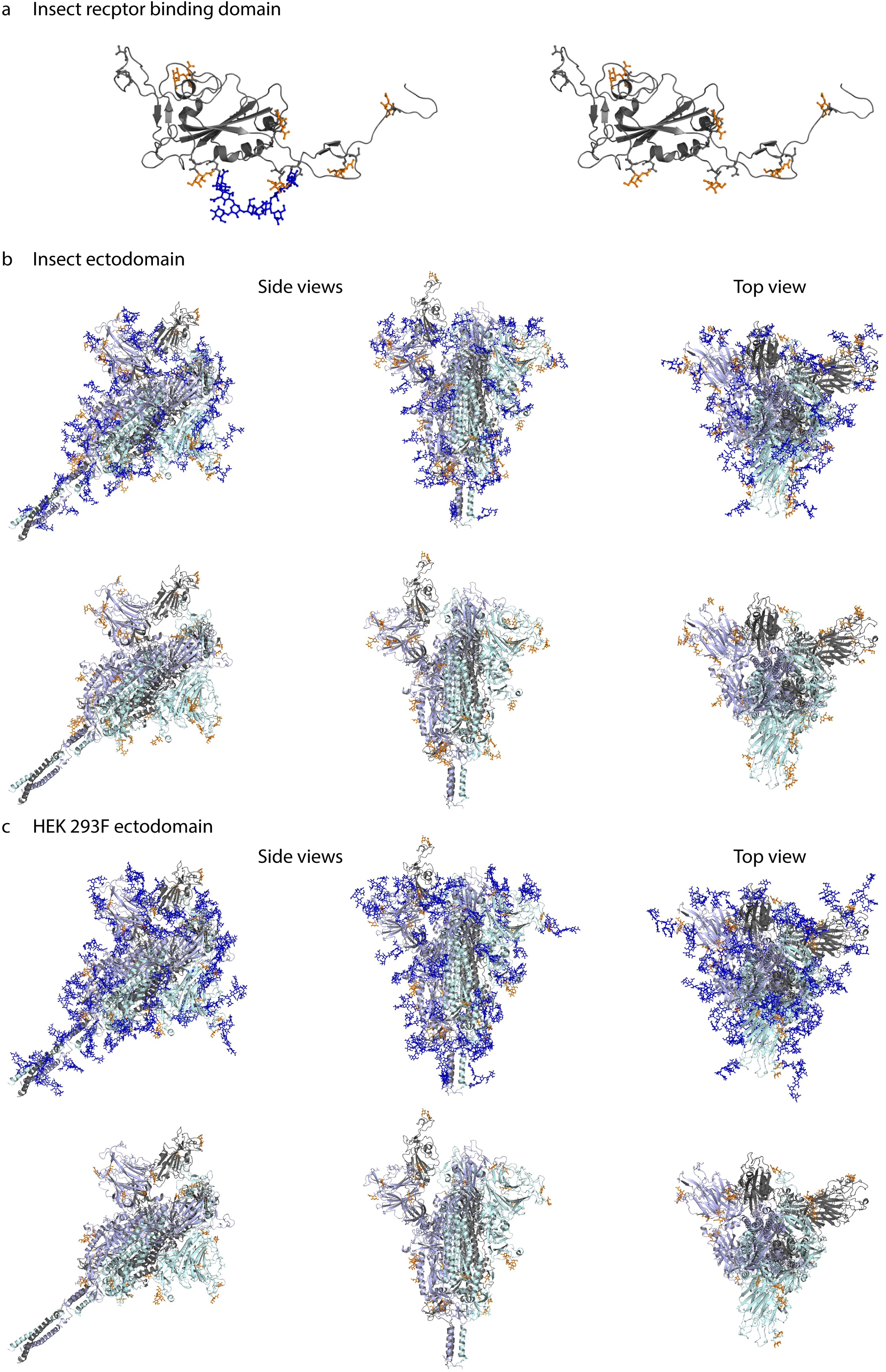
Molecular modelling of identified O-glycosites. Identified site-specific O-glycans (orange) and known N-glycans (blue) were modelled onto 3D reconstructions of SARS-CoV-2 S protein. Sites found on monomeric RBD expressed in insect cells (a), ectdomain expressed in insect cells (b), or ectodomain expressed in HEK 293F cells (c) are shown. Longest identified O-glycan structures are presented at specific sites of each protein. Paucimannose type N-glycan structures are shown on the insect cell-expressed products, whereas structures defined in the full-length glycosylated model by Woo *et al.* [28] are shown on the HEK 293F-expressed ectodomains. Models with and without N-glycans are included in each panel. Only the amino acids covered by different construct designs are shown.

Most of the identified O-glycosites were mapped to amino acids with good solvent accessibility (Fig. S4). For the monomeric RBD expressed in insect cells, O-glycans primarily located to flexible loops surrounding the RBD core (Fig 3A). For the two different cell-derived ectodomains, O-glycans distributed rather evenly over the exposed surface area, blending into the network of known N-glycans, and protruding away from the core (Fig. 3 B,C). Of interest, Thr478, found glycosylated on both ectodomains, was located in the vicinity of receptor binding motif important for interaction with ACE2 (Fig. 3B,C).

A few sites, Ser94, Thr95, Thr1077 and Ser1097 (Figure S4), did not per se localize to amino acids with good solvent accessibility. The 3D modelling revealed that Ser94, which is one of the ambiguously identified sites (94-95 (1x)), is not accessible for attachment of an O-glycan. However, it was possible to attach an O-glycan to Thr95 with only minor adjustment of the neighboring amino acid side chains, which makes Thr95 the likely occupied position. The lack of accessibility at Thr1077 suggests that the HexHexNAcs found on Thr1076 and Thr1077, respectively may be derived from a single core 2 structure at the Thr1076 site. After minor adjustment of the side chain conformation of Ser1097 it was also possible to attach an O-glycan at this position. In summary, although the SAS analysis already provided reasonable information on the surface exposure for the O-glycosites, there were still some ambiguous cases where only the detailed atomistic 3D modelling provided the necessary insight to explain the experimental results.

## Discussion

Enveloped viruses acquire host glycosylation while their proteins travel through the secretory pathway. Glycosylation of viral envelope proteins significantly alter the shape of the protein molecules, shielding the underlying amino acids and thus their recognition by the host immune system. Viruses can acquire both N- and O-linked glycans, but the former is much more widely explored [12]. Here, we aimed to discover O-linked glycan attachment sites on SARS-CoV-2 S protein, and explore the potential implications of such modifications using three different recombinant protein entities. We report 25 unique O-glycosites, most of which were identified unambiguously. This includes the previously identified T323 and T678 adjacent to S1/S2 cleavage site [14–17].

We found similar glycosylation patterns on insect and human cell derived ectodomains, which suggests that the peptides can be glycosylated by isoforms of GalNAc-Ts with broad acceptor substrate specificities. We discovered wide distribution of O-glycosites over the surface areas of the two investigated proteins, and found no bias towards specific regions. A larger proportion of sites harboring core 1 structures (Galβ3GalNAcα-S/T) were identified on the insect cell-derived ectodomain, compared to mostly short immature Tn structures (GalNAcα-S/T) on the HEK 293F-derived ectodomain. Since S2 cells abundantly express paucimannosidic N-glycans, it is possible that the surface is more accessible to the Core-1 enzyme elongating the short Tn structure with a galactose residue [31]. Furthermore, the predominantly dimeric conformation of the insect ectodomain may also provide better access for enzymes. However, since the size exclusion chromatography analysis is done on a purified protein, we do not know, whether it is an accurate representation of the quaternary structure in the cell culture supernatants. In a similar context, more O-glycosites were found on the monomeric RBD compared to ectodomain presented RBD, suggesting better accessibility to the initiating GalNAc-Ts. However, low stoichiometries were estimated, making it unlikely to have an effect on receptor binding. In the future it would be relevant though to map O-glycosites on native viruses derived from specified respiratory cell subtypes.

When looking at molecular dynamics simulations of N-linked glycan molecular movements on S, it is evident that some surfaces of the molecule are still accessible, and these would be the logical regions to find O-glycans [17]. In this context, we were surprised to find, that more than 60 % of our identified sites were located right next to N-glycosite positions. After inspecting the sequences of such O-glycopeptides, we came to realize that they were primarily located on Asn containing peptides unmodified by N-glycans. This is in contrast to deamidated and presumably de-N-glycosylated peptides, where we only in a few exceptions found O-glycans. Importantly, for 7 out of 9 investigated regions in the Orbitrap Fusion Lumos dataset, more than 85 % of non-N-glycosylated peptides were in fact modified with O-glycans, suggesting O-glycosylation machinery in most instances fill in the “empty space” unoccupied by N-glycans. This was confirmed by the Q Exactive HF-X data, where PNGase F N-deglycosylation was performed in O^18^ water, and all the identified peptides contained either N-glycans, or O-glycans, but not both.

All of our identified O-glycosites are of very low occupancy, based on integrated MS1 signals of relevant (glyco)peptides. It is important to consider, however, that glycopeptides exhibit poorer ionization efficiencies, and quantification purely based on signal magnitude is likely underestimating the true stoichiometry. Regardless, low occupancy of O-glycosites close to N-glycosites fits well with published data on S N-glycosylation. The 22 SARS-CoV-2 S N-glycosites are reported to be more than 95 % occupied in recent glycoproteomic analyses of SARS-CoV-2 S [13–17]. Based on our data, it is unlikely, that both N- and O-glycans exist in close vicinity to each other, as we found such O-glycosites almost exclusively on peptides with unoccupied N-X-S/T sequons. It is unlikely that the low abundance of O-glycosites would have biological consequences to protein function or immunogenicity, and should not be of concern for choice of immunogen formulation of a highly N-glycosylated protein. Our analyses suggest that O-glycans rather serve to ensure maximum shielding of the minor fraction of peptides that are unoccupied by N-glycans. Based on our molecular modelling, N- and O-glycans were often “pointed” at different angles away from each other; however, there is likely insufficient accessibility to initiating enzymes, when N-glycans are occupied. In this context, it would be interesting to find out, whether site directed mutagenesis of specific N-linked glycans would increase the occupancy of O-linked glycans in those regions, as well as investigate O-glycan occupancy in less densely N-glycosylated viruses.

In conclusion, sensitive instrumentation and appropriate techniques make it possible to identify low abundance post-translational modifications and accurately map their positions, which allowed us to contribute to a more comprehensive map of SARS-CoV-2 S protein glycosylation, with wide distribution of O-glycosites on the surface of S protein ectodomains. We suggest that O-glycans constitute a minor component of the S protein glycan shield, yet they may serve an important function of covering up the unoccupied N-glycosylation sequons.

## Materials and methods

### Protein expression and purification in S2 cells

The nucleotide sequence for the prefusion-stabilized spike protein ectodomain (aa 16–1208), modified with an N-terminal BiP signal peptide, two proline substitutions (aa 986, 987) an AARA substitution at the furin cleavage site, and a twin strep tag (IBA, GmbH), was synthesized and subcloned into a pExpreS2-1 (ExpreS^2^ion Biotechnologies) vector by Geneart. Transiently transfected *Drosophila melanogaster* S2 cells (ExpreS^2^ Cells, ExpreS^2^ion Biotechnologies) were grown shaking in suspension at 25 °C for three days after which the supernatant was harvested by centrifugation, concentrated and buffer exchanged approximately 10-fold. The prefusion-stabilized spike protein ectodomain was captured on a 5 mL Strep-Tactin XT column (IBA, GmbH) and eluted using 50 mM biotin. The protein was further purified by size exclusion chromatography using a Superdex200 column (Cytiva, Massachusetts, USA) equilibrated in 1 × PBS. By analytical SEC the protein behaves predominantly like a dimer protein.

The nucleotide sequence encoding the spike protein receptor binding domain (RBD, amino acids 305 to 543) fused N terminally to a BiP secretion signal, a 10x His tag and proprietary catcher domain (Adaptvac) was synthesized and subcloned into a pExpreS2-1 (ExpreS^2^ion Biotechnologies) vector by Geneart. The protein was expressed transiently in insect cells (ExpreS^2^ Cells, ExpreS^2^ion Biotechnologies) and harvested after three days of culture. The cell supernatant was harvested by centrifugation and concentrated (10-fold) and buffer exchanged to loading buffer (10-fold) by tangential flow filtration (TFF). The protein was loaded onto a 1 mL HisTrap excel (Cytiva, Massachusetts, USA) and washed to baseline in loading buffer. The protein was eluted with Imidazole. Elution fractions were concentrated, and the protein was loaded onto a gel filtration column (Superdex 200pg, Cytiva) equilibrated in 1 × PBS (Gibco). The RBD protein is monomer in solution by analytical SEC analysis (not shown).

### Protein expression and purification in HEK 293 F cells

A synthetic gene encoding SARS-CoV-2 S (sequence derived from Genbank accession: MN908947.3) was cloned into a customized DNA vector for expression in mammalian tissue culture. Final expression constructs featured a fragment encoding the native S ectodomain, including viral signal peptide (residues 1 – 1214), with prefusion stabilizing and 986P, 987P variants first reported by [18]. HEK 293F-expressed S protein featured a C-terminal leucine zipper (GCN4) motif, and His8-tag for Ni^2+^-affinity purification. Pure S-encoding plasmid DNA was transfected into HEK 293F cells (approximately 10^6^ mL^−1^) using PEI (polyethylimine) MAX at 5:1 w/w (final culture DNA concentration approximately 1 μg mL^−1^). After 96 hours condition media supernatant was harvested by low-speed centrifugation, and recombinant S trimers were purified directly from media by IMAC using a 5 mL HisTrap FF crude column (GE). Protein-containing fractions were pooled and concentrated before application to a Superose 6 10/300 GL column (GE) for final purification via size-exclusion chromatography.

### In-gel digestion

2 × 20 μg of each of the three different SARS-CoV-2 spike proteins were mixed with 4x NuPAGE™ LDS sample buffer (Thermo Fisher Scientific) and up to 10 mM dithiothreitol (DTT, Sigma-Aldrich). The samples were run on Novex 4-12 % gradient gel (Bis-Tris) in 1x NuPAGE™ MES buffer (Invitrogen) at 150 V for 60 min on ice followed by 30 min at 200 V. The gel was then stained with InstantBlue® protein stain (Abcam). Bands of interest were excised, cut into smaller pieces (1 × 1 mm), rinsed with water and 2x 30 min washed in 50 mM ammonium bicarbonate (AmBic, Sigma-Aldrich) at 37 °C. The gel pieces were then shrunk in 100 % acetonitrile (ACN) and shaken for 5 min. Solvent was removed, and gel pieces rehydrated in 10 mM DTT in 50 mM AmBic, followed by 40 min incubation at 60 °C. The gel pieces were again shrunk in 100 % ACN and shaken for 30 min. Solvent was removed, and gel pieces rehydrated in 55 mM iodoacetamide (IAA, Sigma-Aldrich) in 50 mM AmBic, followed by 40 min incubation at room temperature. The gel pieces were again shrunk in 100 % ACN and shaken for 15 min. The solvent was removed, gel pieces briefly rinsed with 50 mM AmBic and rehydrated in a small volume (10 μL) of 50 mM AmBic supplemented with 1 U PNGase F (Roche) at 37 °C for 30 min. The gel pieces were then topped up with 50 mM AmBic to cover the surface and incubated at 37 °C over night. After N-glycan removal, the gel pieces were washed in 50 mM AmBic at 37 °C for 30 min, followed by shrinking in 100 % ACN for 5 min. The solvent was removed, briefly rinsed with 50 mM AmBic and rehydrated in a small volume (15 μL) of 50 mM AmBic supplemented with 0.5 μg of chymotrypsin (Roche) or Glu-C (Roche) at 25 °C for 30 min. The gel pieces were then topped up with 50 mM AmBic to cover the surface and incubated at 25 °C for 8 hours. The Glu-C treated gel pieces were then dried by SpeedVac and rehydrated in a small volume (15 μL) of 50 mM AmBic supplemented with 0.5 μg of trypsin (Roche) at 37 °C for 30 min. The gel pieces were then topped up with 50 mM AmBic to cover the surface and incubated at 37 °C for 8 hours. Peptides were collected followed by 2x 15 min water bath sonication in 300 μL 50 mM AmBic followed by 15 min sonication in 300 μL 50 % ACN. For individual samples, all supernatants were pooled and dried using SpeedVac to less than 500 μL. The peptides were desialylated in 50 mM sodium acetate (Merck) buffer pH 5 using *Clostridium perfringens* neuraminidase (Sigma-Aldrich) at 0.1 U/mL for 2 hours at 37 °C. Desialylated peptides were stage-tip purified as previously described, dried, and reconstituted in 12 μL of 0.1 % formic acid prior to analysis.

### Mass spectrometry analysis with Orbitrap Fusion Lumos mass spectrometer

LC MS/MS site-specific O-glycopeptide analysis of was performed on EASY-nLC1200 UHPLC (Thermo Fisher Scientific) interfaced via nanoSpray Flex ion source to an Orbitrap Fusion Lumos Tribrid MS (Thermo Fisher Scientific). The nLC was operated in a single analytical column set up using PicoFrit Emitters (New Objectives, 75 mm inner diameter) packed in-house with Reprosil-Pure-AQ C18 phase (Dr. Maisch, 1.9-mm particle size, 19-21 cm column length). Each sample was injected onto the column and eluted in gradients from 3 to 32% B in 95 min, from 32 to 100% B in 10 min and 100 % B in 15 min at 200 nL/min (Solvent A, 100 % H2O; Solvent B, 80 % acetonitrile; both containing 0.1% (v/v) formic acid). A precursor MS1 scan (m/z 350–1,700) was acquired in the Orbitrap at the nominal resolution setting of 120,000, followed by Orbitrap HCD-MS2 and ETD-MS2 at the nominal resolution setting of 60,000 of the five most abundant multiply charged precursors in the MS1 spectrum; a minimum MS1 signal threshold of 50,000 was used for triggering data-dependent fragmentation events. Stepped collision energy +/-5 % at 27 % was used for HCD MS/MS fragmentation and charge dependent calibrated ETD reaction time was used with CID supplemental activation at 30 % collision energy for ETD MS/MS fragmentation.

For the site-specific glycopeptide identification, the corresponding HCD MS/MS and ETD MS/MS were analyzed by Proteome discoverer 2.2 software (Thermo Fisher Scientific) using Sequest HT as a searching engine. Carbamidomethylation at cysteine was used as fixed modification and oxidation at methionine, asparagine deamidation, and HexNAc or Hex-HexNac at serine/threonine/tyrosine were used as variable modifications. Precursor mass tolerance was set to 10 ppm and fragment ion mass tolerance was set to 0.02 Da. Data were searched against the human-specific UniProt KB/SwissProt-reviewed database downloaded on January, 2013 and construct-dependent viral protein sequence databases. All spectra of interest were manually inspected and validated to prove the correct peptide identification and glyco-site localization.

### In-solution digestion and mass spectrometry analysis with Q Exactive HF-X mass spectrometer

The COV19 double mutant spike trimer from HEK 293F cell was prepared for MS analysis as previously described [27] with minor modifications. In brief, the protein (50 μg) was denatured and aliquots (10 μg) were digested under five different protease conditions including chymotrypsin, a combination of trypsin and chymotrypsin, trypsin, elastase and subtilisin as described. All samples were then pooled and deglycosylated by Endo H followed by PNGase F in O18-water.

The combined sample was analyzed on an Q Exactive HF-X mass spectrometer (Thermo). Each sample was run twice as replicate. Samples were injected directly onto a 25 cm, 100 μm ID column packed with BEH 1.7 μm C18 resin (Waters). Samples were separated at a flow rate of 300 nL/min on a nLC 1200 (Thermo). Solutions A and B were 0.1 % formic acid in 5 % and 80 % acetonitrile, respectively. A gradient of 1–25 % B over 160 min, an increase to 40 % B over 40 min, an increase to 90 % B over another 10 min and held at 90 % B for 30 min was used for a 240 min total run time. Column was re-equilibrated with solution A prior to the injection of sample. Peptides were eluted directly from the tip of the column and nanosprayed directly into the mass spectrometer by application of 2.8 kV voltage at the back of the column. The HFX was operated in a data dependent mode. Full MS1 scans were collected in the Orbitrap at 120k resolution. The ten most abundant ions per scan were selected for HCD MS/MS at 25NCE. Dynamic exclusion was enabled with exclusion duration of 10 s and singly charged ions were excluded.

The MS data were processed as described previously [27] with minor modifications. The data were searched against the proteome database and quantified using peak area in Integrated Proteomics Pipeline-IP2. The parameters were set as: MS1 tolerance ≤50 ppm, MS2 tolerance ≤20 ppm, no enzyme specificity, carboxyamidomethylation (+57.02146 C) as a fixed modification, and oxidation (+15.9994 M), deamidation (+2.988261 N), GlcNAc (+203.079373 N), GalNAc (+203.079373 S/T), Gal-GalNAc (+365.136681 S/T) and pyroglutamate formation from N-terminal glutamine residue (−17.026549 Q) as variable modifications.

### Molecular modelling

We used Conformational Analysis Tools software (www.md-simulations.de/CAT/) interfaced with STRIDE [29] to estimate the solvent accessible surface (SAS) of the amino acids. To visualize the 3D positions of O-glycosites in the context of a fully-glycosylated full-length S protein structure, we downloaded the model 6vsb_1_1_1 from http://www.charmm-gui.org/?doc=archive&lib=covid19 [28]. This model contains N-glycans in agreement with [16]. The O-glycans were attached to the model manually and optimized using YASARA [30].

## Data availability

All mass spectrometry raw data will be deposited to the ProteomeXchange Consortium via the PRIDE partner repository (https://www.ebi.ac.uk/pride/) and will be publically available upon first online publication.

## Funding

This work was supported by the European Commission (GlycoSkin H2020-ERC), Lundbeck Foundation (R219-2016-545), Danish National Research Foundation (DNRF107), Carlsberg Foundation (Semper Ardens), Gudbjørg og Ejnar Honorés Fond, NIH AI144462, and NIH AI113867.

## Author contributions

Conceptualization, I.B., A.J.T., S.Y.V., J.C.P. and H.H.W.; Methodology, I.B., A.J.T., X.W., M.S., C.F., M.F., J.R.Y., and S.Y.V.; Software, M.F., J.R.Y., and S.Y.V.; Validation, I.B. and X.W.; Formal Analysis, I.B., X.W., M.F., and S.Y.V.; Investigation, I.B., A.J.T., X.W., M.S., C.F., M.F., J.K.D., S.Y.V., J.C.P., and H.H.W.; Resources, M.S., C.F., M.F., J.R.Y., A.S., S.Y.V., J.C.P., and H.H.W.; Data Curation, I.B., X.W., and S.Y.V.; Writing – Original Draft Preparation, I.B. and H.H.W.; Writing – Review & Editing, I.B., A.J.T., X.W., M.S., C.F., M.F., J.K.D., J.R.Y., A.S., S.Y.V., J.C.P., and H.H.W.; Visualization, I.B., X.W., M.F., and H.H.W.; Supervision, J.R.Y., A.S., S.Y.V., J.C.P., and H.H.W.; Project Administration, I.B. and H.H.W.; Funding Acquisition, I.B., A.S., J.C.P., and H.H.W.

## Conflicts of Interest

The authors declare no conflict of interest. The funders had no role in the design of the study; in the collection, analyses, or interpretation of data; in the writing of the manuscript, or in the decision to publish the results.

## Notes

### Competing Interest Statement

The authors have declared no competing interest.

