## Supplementary material for "Site-specific O-glycosylation analysis of SARS-CoV-2 spike protein produced in insect and human cells": Figures S1 and S2

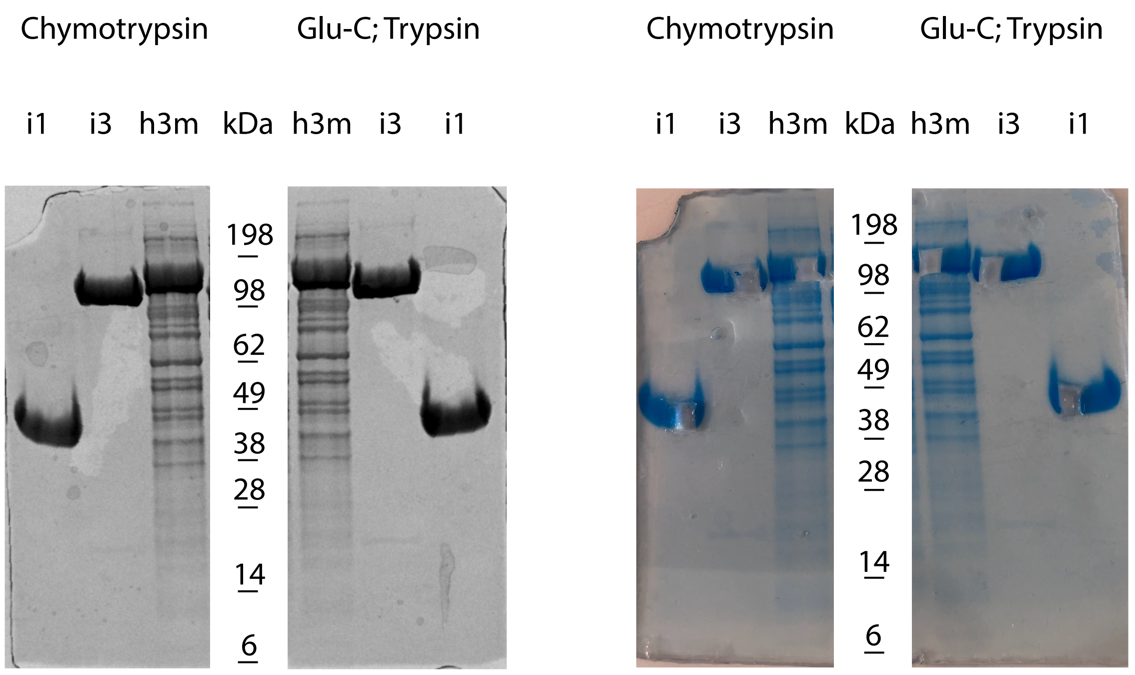


**Figure S1. In gel digestion of S proteins.** 20 µg per lane of of SARS- CoV-2 S monomeric RBD expressed in insect cells (i1), ectodomain expressed in insect cells (i3), and ectodomain expressed in HEK 293F cells (h3m) were separated on a 4-12 % Bis-Tris gradient gel in 1x MES buffer at 150 V for 1 hour on ice followed by 30 min at 200 V in duplicates and stained with InstantBlue® protein stain. One set of proteins was used for chymotrypsin digestion, whereas the other was used for Glu-C followed by trypsin digestion. The image on the right shows excised gel regions for downstream analysis.

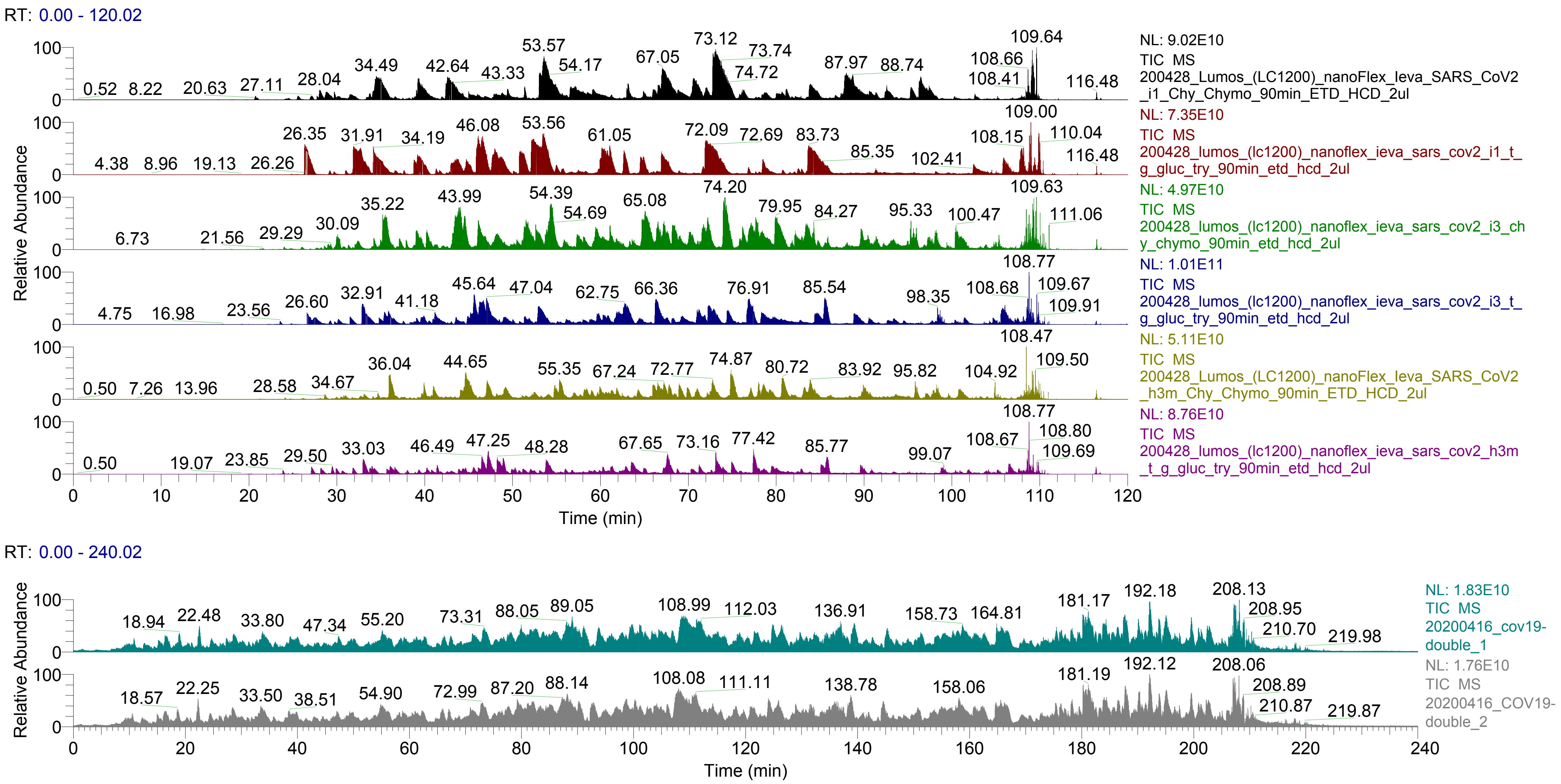


**Figure S2. Total ion chromatograms of analyzed samples.** Raw files were opened using Thermo Xcalibur Browser and total ion chromatograms (TIC) exported. (a) 2 out of 12 µL of each desalted sample was injected to a Thermo Fisher nLC-MS/MS Fusion Lumos Trybrid mass spectrometer. The peptides were eluted over a 120 min gradient. TIC of the six different samples (Fig. S1) over time are shown. (b) TIC of HEK 293F expressed S samples digested in solution analyzed on a Thermo Fisher Q Exactive HF-X mass spectrometer in duplicate. The peptides were eluted over a 240 min gradient.
