## Supplementary material for "Site-specific O-glycosylation analysis of SARS-CoV-2 spike protein produced in insect and human cells": Figure S3

**Figure S3. Examples of ETD MS2 spectra representing the different O-glycosites identified using the Orbitrap Fusion Lumos mass spectrometer.** c and z product fragment ions are annotated in the peptide sequence. The source raw file is indicated in the top left corner (i1 – monomeric RBD expressed in insect cells, i3 – ectodomain expressed in insect cells, h3m – ectodomain expressed in HEK 293F cells).

T63

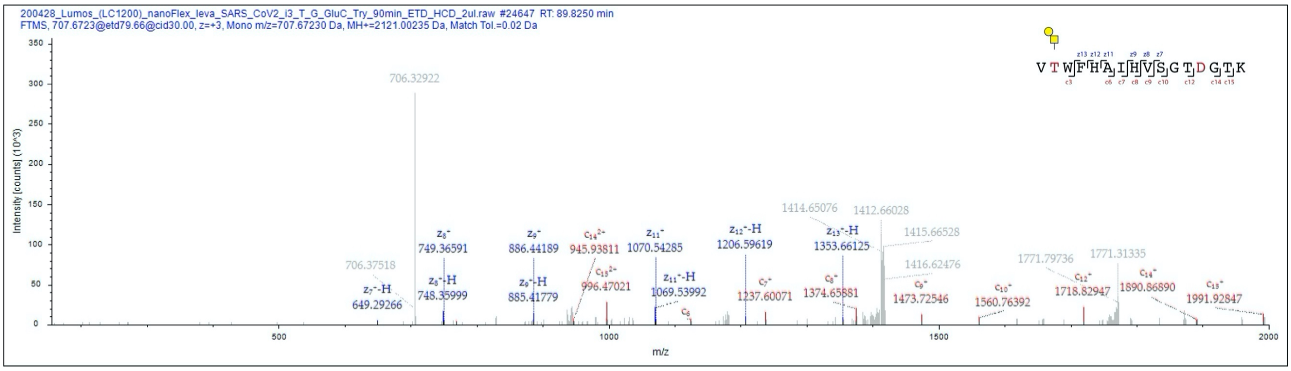


S71, T73

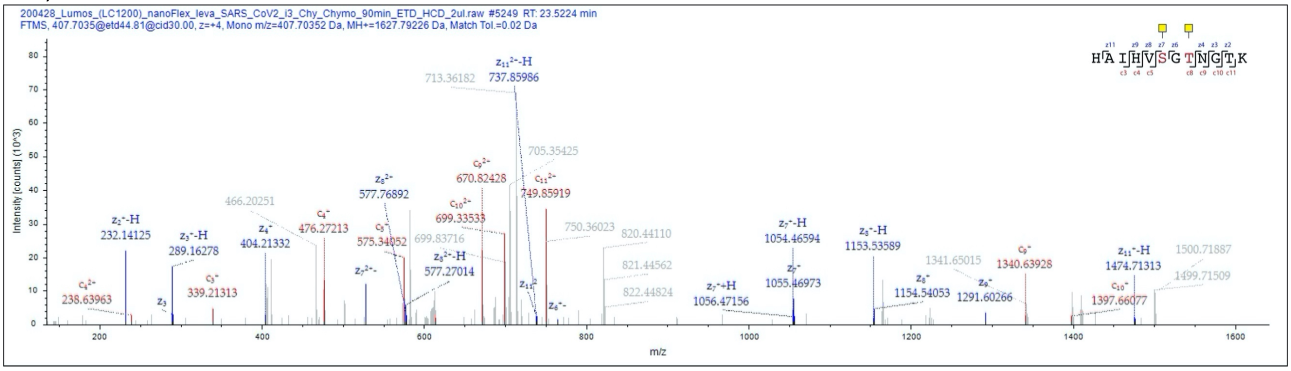


T73, T76

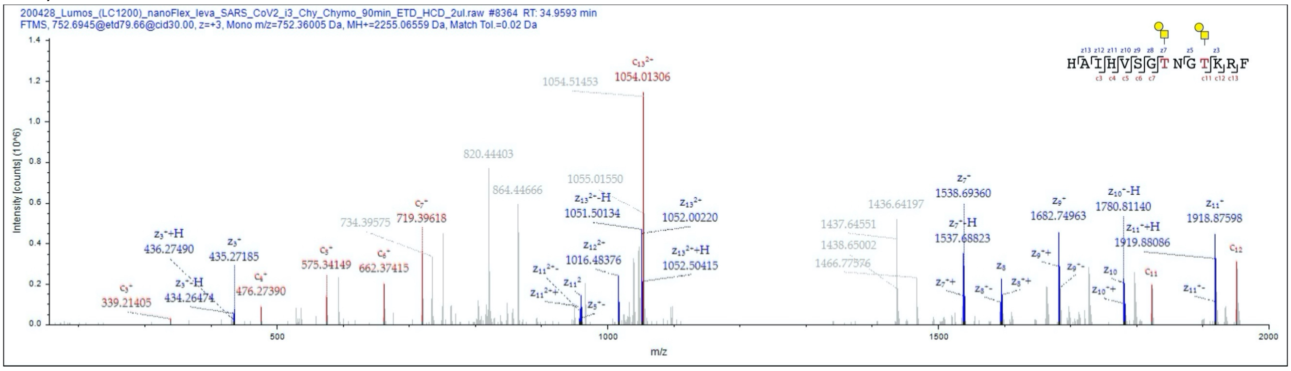


94-95 (1x)

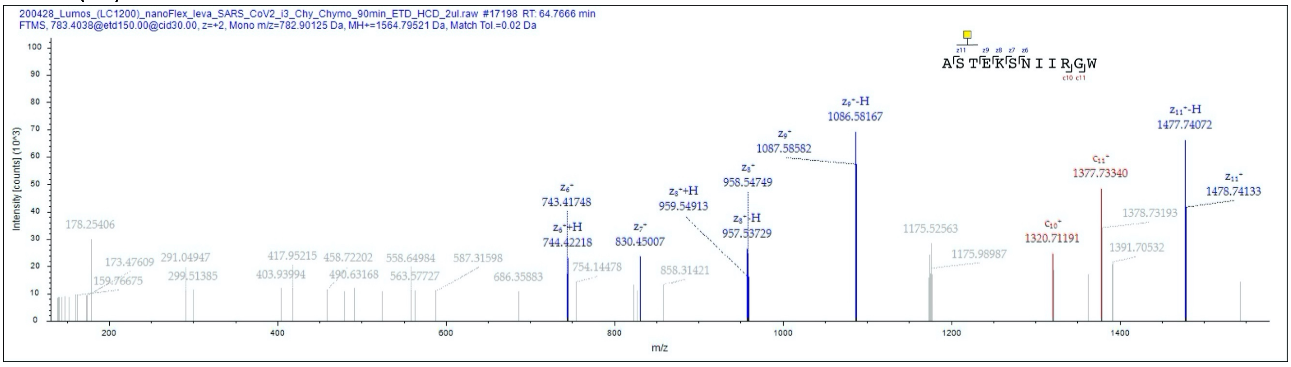

T124

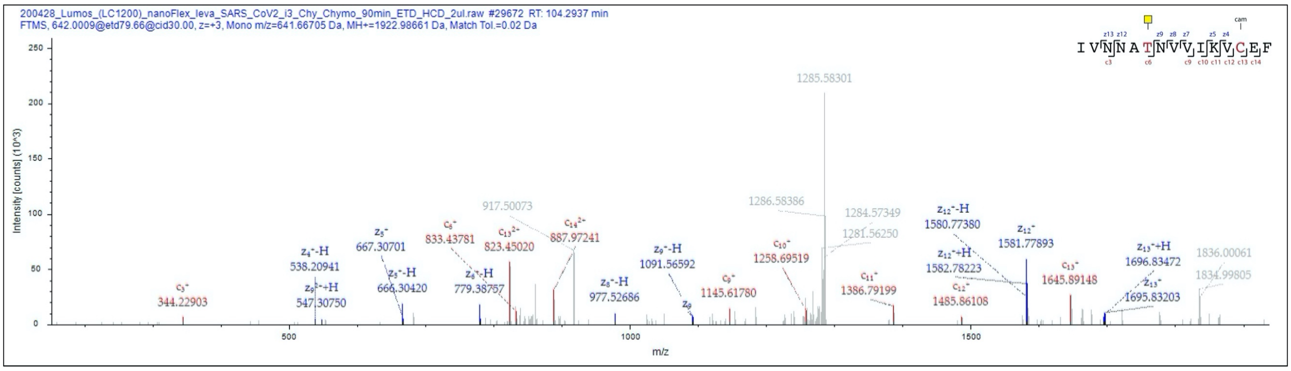


S151

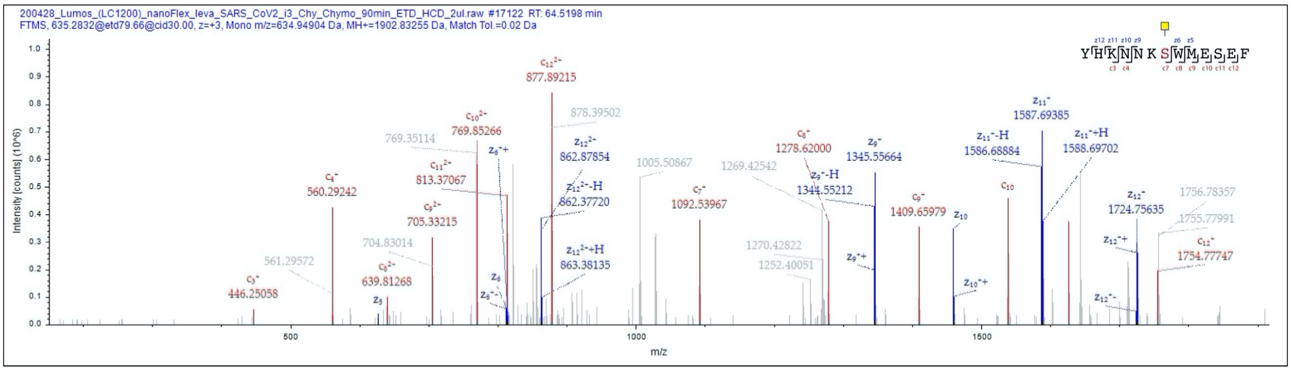


T315

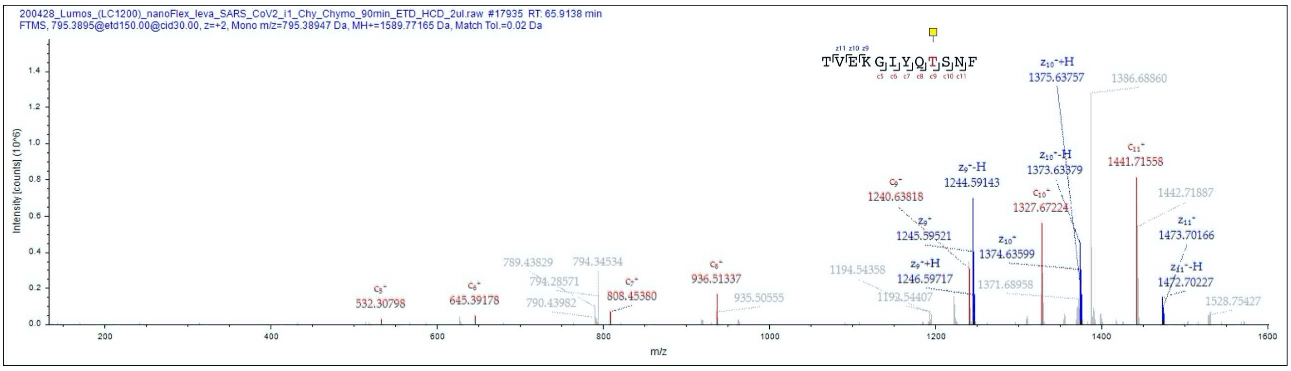


T323

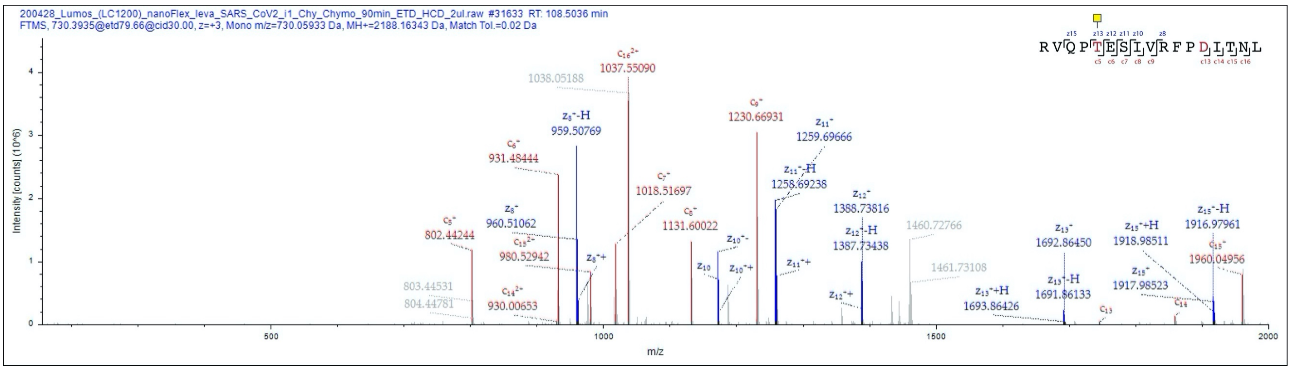

T333

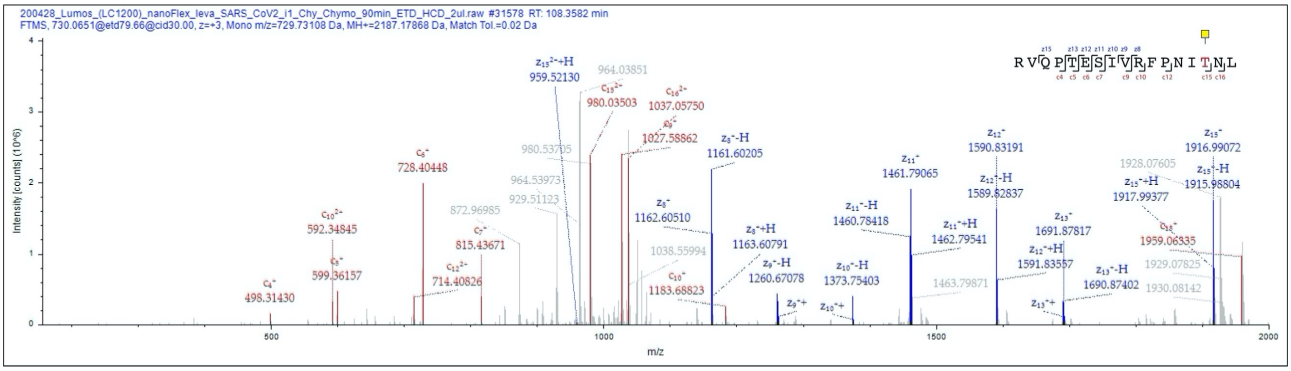


T345

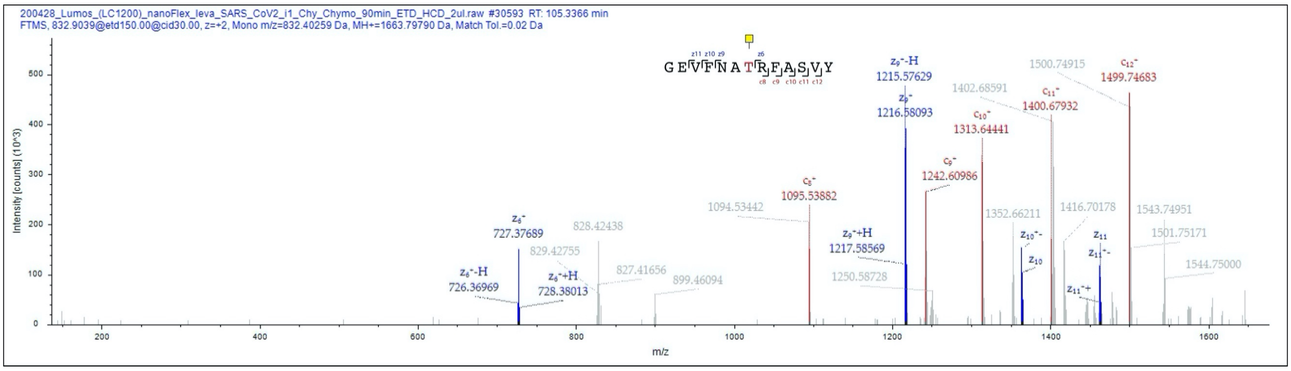


T415

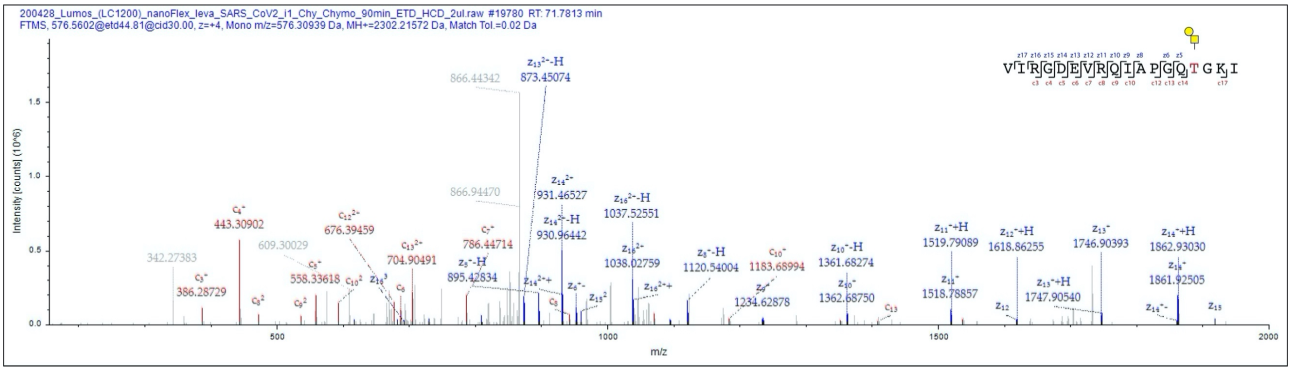


T478

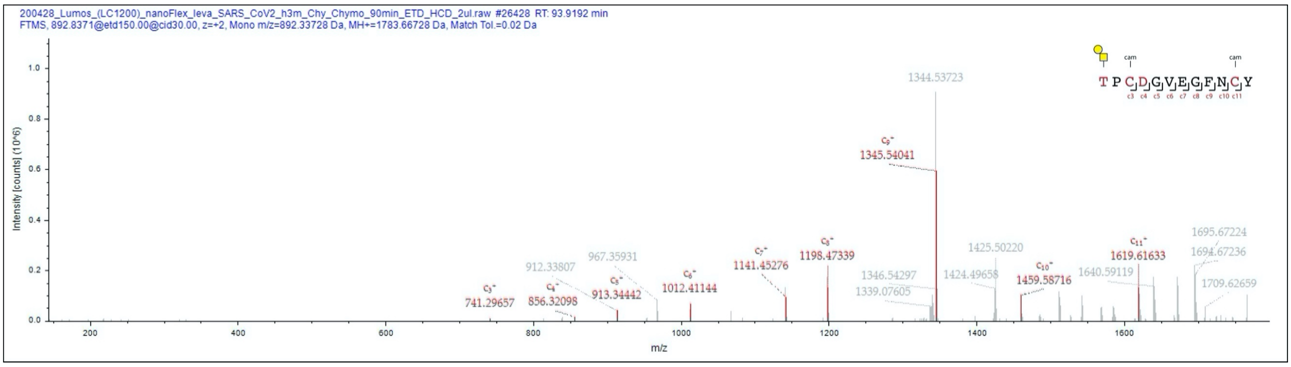

T523

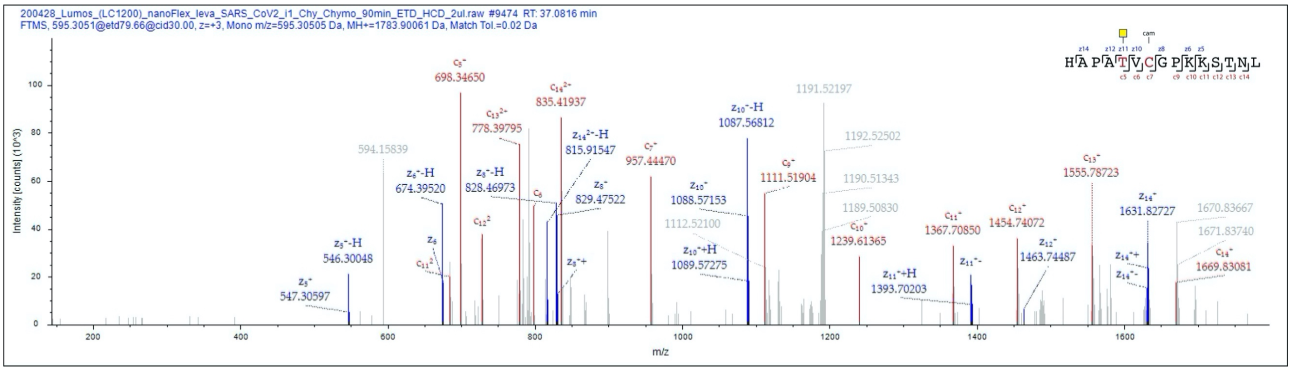


T618

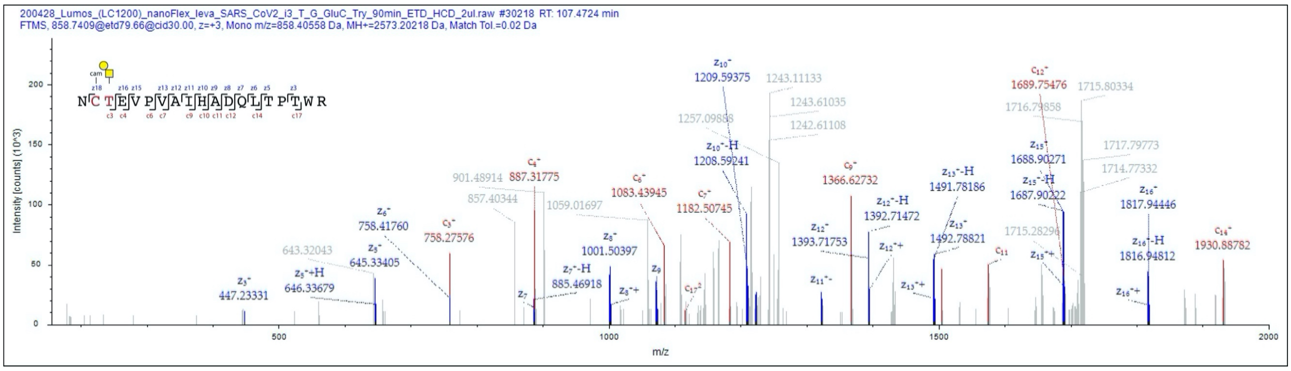


T676, T678

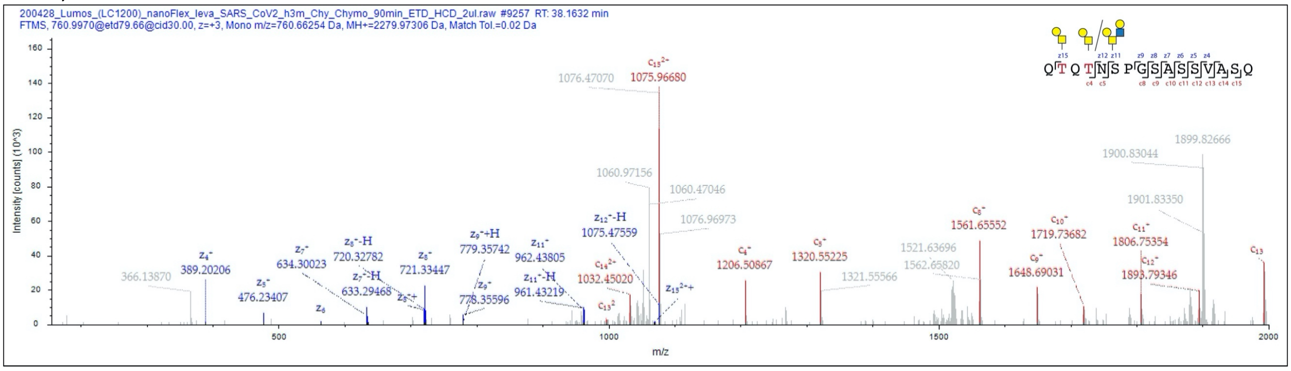


S803

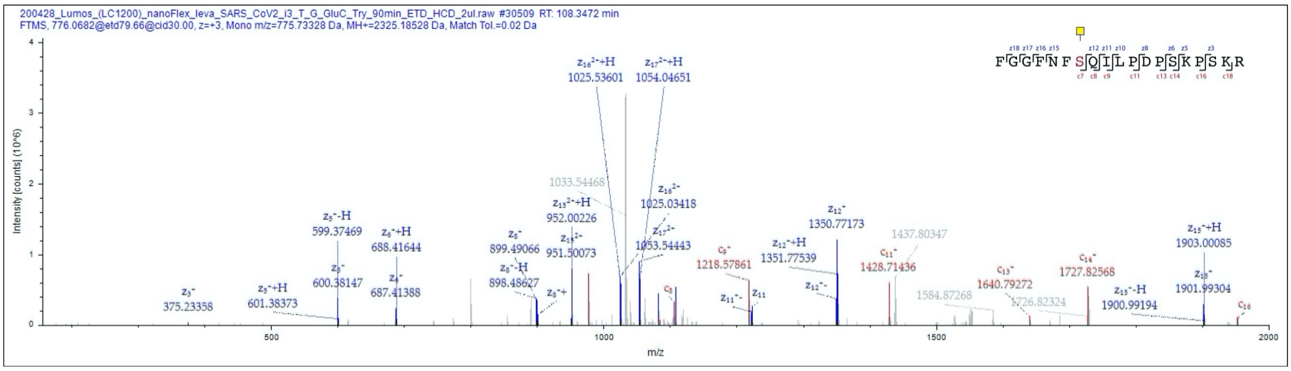

T1076, T1077

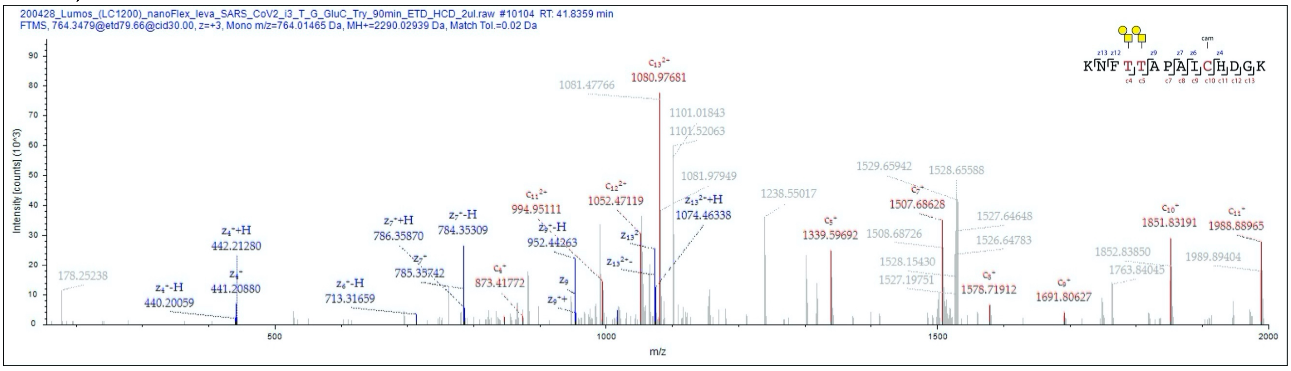


T1097

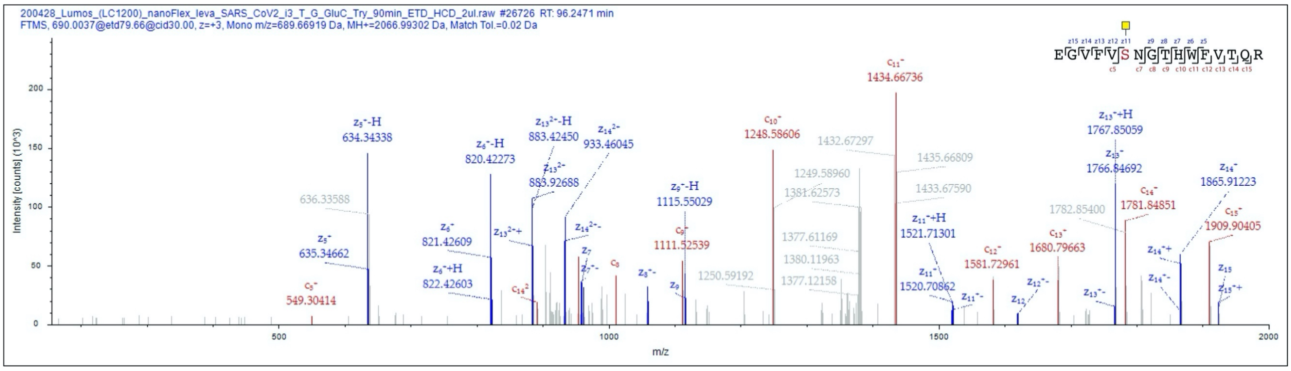


T1097, T1100

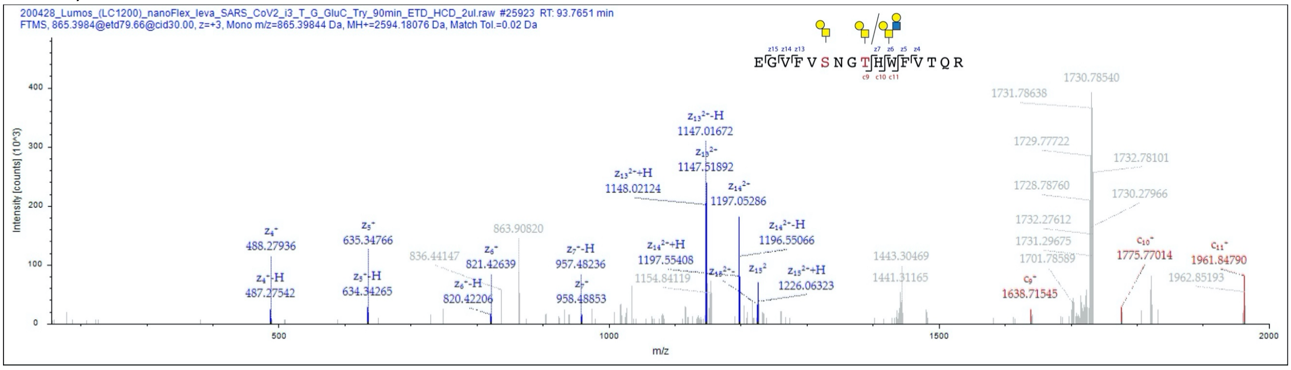
