## Supplementary material for "Site-specific O-glycosylation analysis of SARS-CoV-2 spike protein produced in insect and human cells": Figure S4

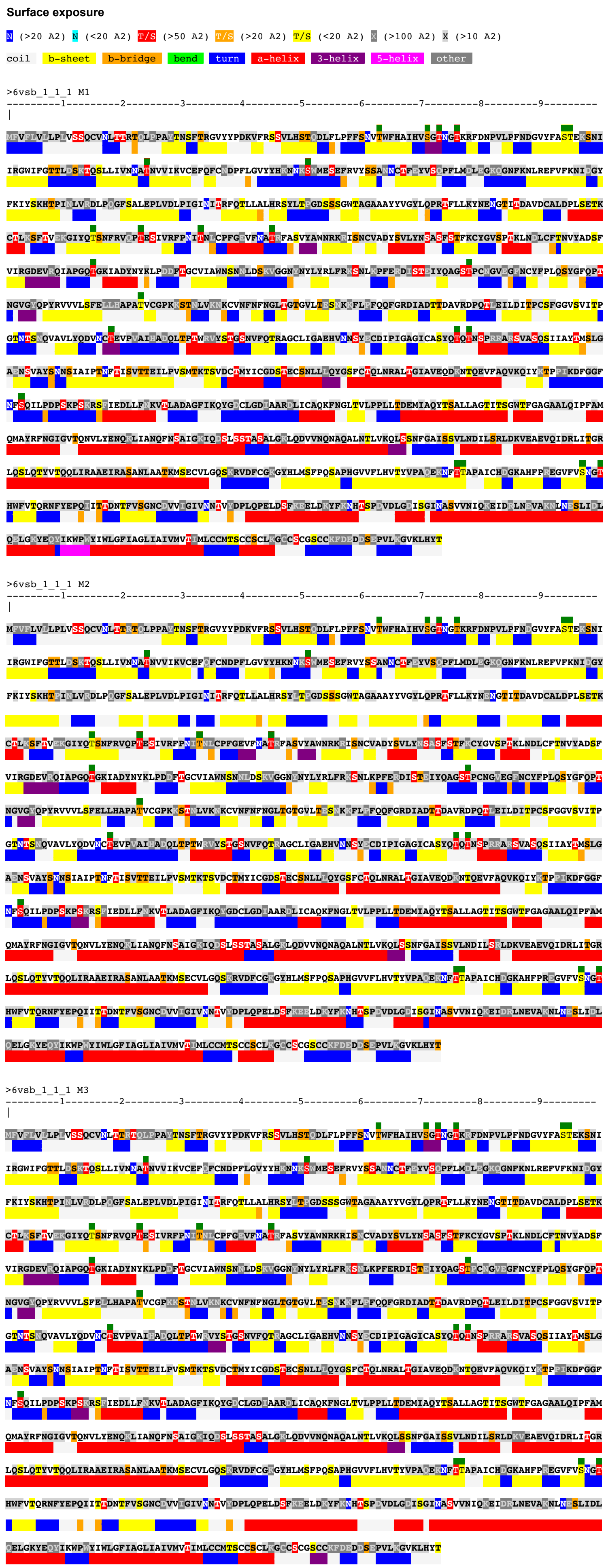


**Surface areas of selected residues**

| Label | A2 |
|---|---|
| A:THR_63 | 36 |
| B:THR_63 | 31 |
| C:THR_63 | 21 |
| A:SER_71 | 30 |
| B:SER_71 | 42 |
| C:SER_71 | 43 |
| A:THR_73 | 130 |
| B:THR_73 | 125 |
| C:THR_73 | 136 |
| A:THR_76 | 51 |
| B:THR_76 | 45 |
| C:THR_76 | 86 |
| A:SER_94 | 2 |
| B:SER_94 | 3 |
| C:SER_94 | 3 |
| A:THR_95 | 5 |
| B:THR_95 | 0 |
| C:THR_95 | 2 |
| A:THR_124 | 106 |
| B:THR_124 | 97 |
| C:THR_124 | 105 |
| A:SER_151 | 56 |
| B:SER_151 | 74 |
| C:SER_151 | 62 |
| A:THR_315 | 13 |
| B:THR_315 | 9 |
| C:THR_315 | 16 |
| A:THR_323 | 73 |
| B:THR_323 | 75 |
| C:THR_323 | 76 |
| A:THR_333 | 120 |
| B:THR_333 | 37 |
| C:THR_333 | 34 |
| A:THR_345 | 148 |
| B:THR_345 | 145 |
| C:THR_345 | 123 |
| A:THR_415 | 101 |
| B:THR_415 | 50 |
| C:THR_415 | 91 |
| A:THR_478 | 97 |
| B:THR_478 | 101 |
| C:THR_478 | 48 |
| A:THR_523 | 17 |
| B:THR_523 | 38 |
| C:THR_523 | 23 |
| A:THR_618 | 89 |
| B:THR_618 | 93 |
| C:THR_618 | 94 |
| A:THR_676 | 72 |
| B:THR_676 | 67 |
| C:THR_676 | 68 |
| A:THR_678 | 60 |
| B:THR_678 | 54 |
| C:THR_678 | 62 |
| A:SER_803 | 79 |
| B:SER_803 | 79 |
| C:SER_803 | 79 |
| A:THR_1076 | 66 |
| B:THR_1076 | 65 |
| C:THR_1076 | 60 |
| A:THR_1077 | 1 |
| B:THR_1077 | 1 |
| C:THR_1077 | 1 |
| A:SER_1097 | 2 |
| B:SER_1097 | 2 |
| C:SER_1097 | 2 |
| A:THR_1100 | 131 |
| B:THR_1100 | 129 |
| C:THR_1100 | 128 |

**Figure S4. Surface exposure analysis.** Conformational Analysis Tools (CAT) were used to estimate the solvent accessible surface (SAS) of the amino acids in the 6vsb_1_1_1 model. SAS (upper lines) of the three individual subunits of protein S are shown in the context of secondary structure (lower lines). Color code for SAS range in square angstroms and secondary structure are indicated. Small dark green squares above the sequence indicate O-glycosylated amino acids identified in the study. SAS values for these select positions are shown in the table on the right.
