## Supplementary material for "Site-specific O-glycosylation analysis of SARS-CoV-2 spike protein produced in insect and human cells": Dataset S7: Dataset S7 legend.rtf

Dataset S7. The zipped file contains spectra of O-glycosylated peptides included in occupancy quantification (Figure 2b and Dataset S6) of data derived from the Q Exactive HF-X experiments. Examples of non-O-glycosylated peptide spectra are also included. The naming of the files follows the following format O-glycosite position-number-Y/N, where the number indicates the order number in Dataset S6, and the Y/N informs, whether the peptide is O-glycosylated (Y - yes), or not (N - no).
