## Supplementary material for "Site-specific O-glycosylation analysis of SARS-CoV-2 spike protein produced in insect and human cells": Dataset S7: S803-1-Y.pdf

### Fragment Ion View

Project name: 20200330COV19
Sample name: 20200416_COV19-double_1
Created date: 2020-04-18 15:22:56.0
Search name:
Activation type: CID

Peptide sequence: K.DFGGFNFS(203.079373)QILPDPSK.P
Scan number: 55714
Calculated M+H: 1971.9335
Measured M+H: 1971.9445
Charge state: 2
File name: 20200416_COV19-double_1
search type: light

**Ions:**
a ☐1⁺ ☐2⁺ ☐3⁺
b ☑1⁺ ☐2⁺ ☐3⁺
c ☐1⁺ ☐2⁺ ☐3⁺
x ☐1⁺ ☐2⁺ ☐3⁺
y ☑1⁺ ☐2⁺ ☐3⁺
z ☐1⁺ ☐2⁺ ☐3⁺
[Deselect All]

**Neutral Loss:**
☐ $H_2O$ (o)
☐ $NH_3$ (*)
☐ $H_3PO_4$ (^)
☐ Precursor (#)
☐ Precursor with reduced charges

Mass Type:
◉Mono ○Avg

Mass Tol: 20
○Dalton ◉PPM

Update

Peak Assignment:
◉Most Intense
○Nearest Match
☐Peak Detect
☐Precursor Removal

Peak Labels:
◉Ion ○m/z
○None

Width: 750

Height: 450

DFGGFNFSQILPDPSK, MH+ 1971.9334, m/z 986.4704
File: 20200416_COV19-double_1, Scan: 55714, Precursor m/z: 986.4759, Charge: 2

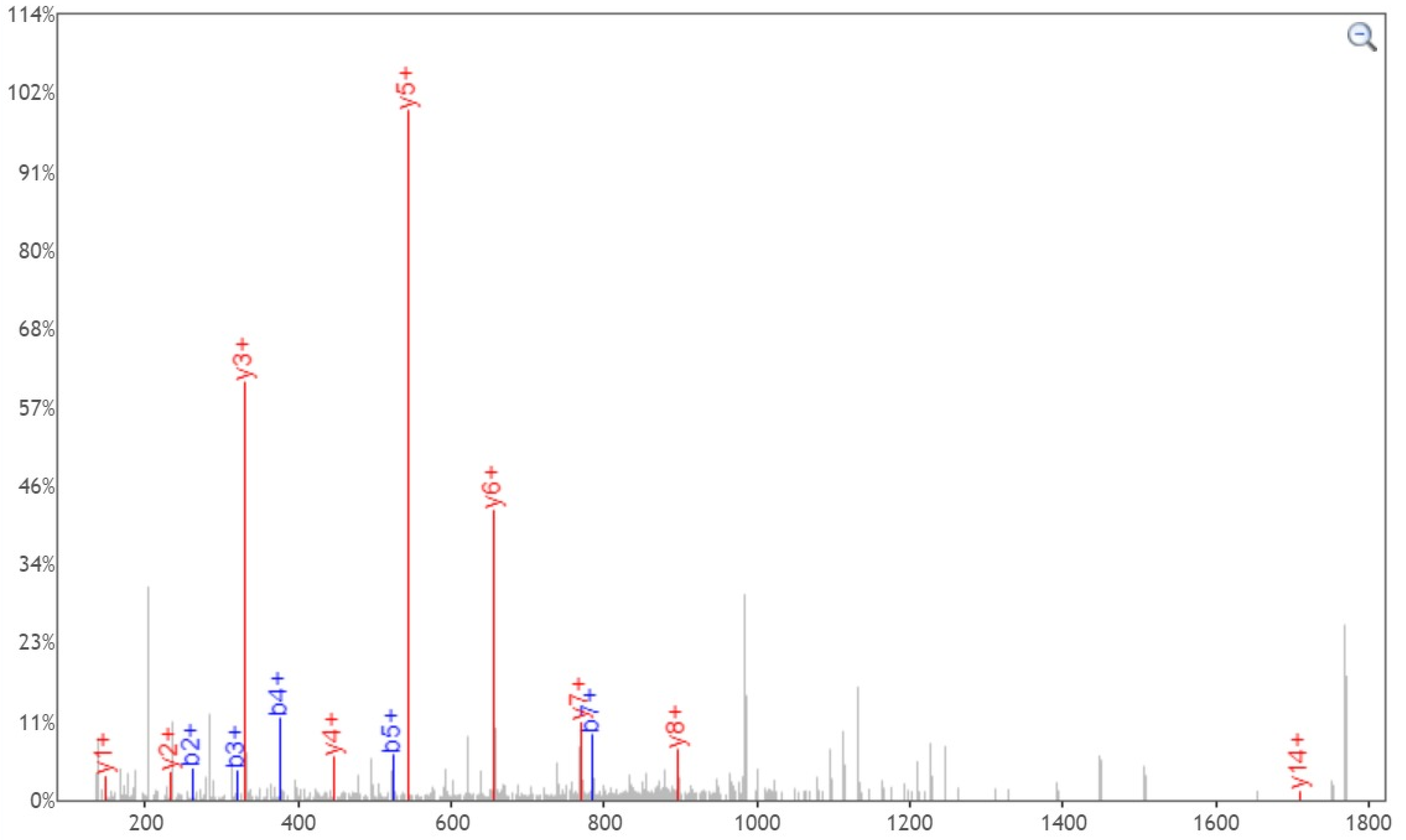


Click and drag in the plot to zoom X: ☑ Y: ☐ Zoom Out Print ☐Enable tooltip

| b+ | # | Seq | # | y+ |
|---|---|---|---|---|
| 116.0342 | 1 | D | 16 | |
| 263.1026 | 2 | F | 15 | 1856.9065 |
| 320.1241 | 3 | G | 14 | 1709.8381 |
| 377.1456 | 4 | G | 13 | 1652.8166 |
| 524.2140 | 5 | F | 12 | 1595.7952 |
| 638.2569 | 6 | N | 11 | 1448.7268 |
| 785.3253 | 7 | F | 10 | 1334.6838 |
| 1075.4367 | 8 | S | 9 | 1187.6154 |
| 1203.4953 | 9 | Q | 8 | 897.5040 |
| 1316.5794 | 10 | I | 7 | 769.4454 |
| 1429.6634 | 11 | L | 6 | 656.3614 |
| 1526.7162 | 12 | P | 5 | 543.2773 |
| 1641.7431 | 13 | D | 4 | 446.2245 |
| 1738.7959 | 14 | P | 3 | 331.1976 |
| 1825.8279 | 15 | S | 2 | 234.1448 |
| | 16 | K | 1 | 147.1128 |

[Click] to move table

Static Modifications:
C: 57.02146

Variable Modifications:
S: 203.079373
