## Supplementary material for "Site-specific O-glycosylation analysis of SARS-CoV-2 spike protein produced in insect and human cells": Dataset S7: S1097-1-Y.pdf

### Fragment Ion View

Project name: 20200330COV19
Sample name: 20200416_COV19-double_1
Created date: 2020-04-18 15:22:56.0
Search name:
Activation type: CID

Peptide sequence: R.EGVFVS(203.079373)NGTHW.F
Scan number: 44189
Calculated M+H: 1435.6488
Measured M+H: 1435.6528
Charge state: 2
File name: 20200416_COV19-double_1
search type: light

Ions:
a 1+ 2+ 3+
b 1+ 2+ 3+
c 1+ 2+ 3+
x 1+ 2+ 3+
y 1+ 2+ 3+
z 1+ 2+ 3+
[Deselect All]

Neutral Loss:
$H_2O$ (o)
$NH_3$ (*)
$H_3PO_4$ (^)
Precursor (#)
Precursor with reduced charges

Mass Type:
Mono Avg

Mass Tol: 20
Dalton PPM

Update

Peak Assignment:
Most Intense
Nearest Match
Peak Detect
Precursor Removal

Peak Labels:
Ion m/z
None

Width: 750

Height: 450

**EGVFVSNGTHW**, MH+ 1435.6488, m/z 718.3281
File: 20200416_COV19-double_1, Scan: 44189, Precursor m/z: 718.3301, Charge: 2

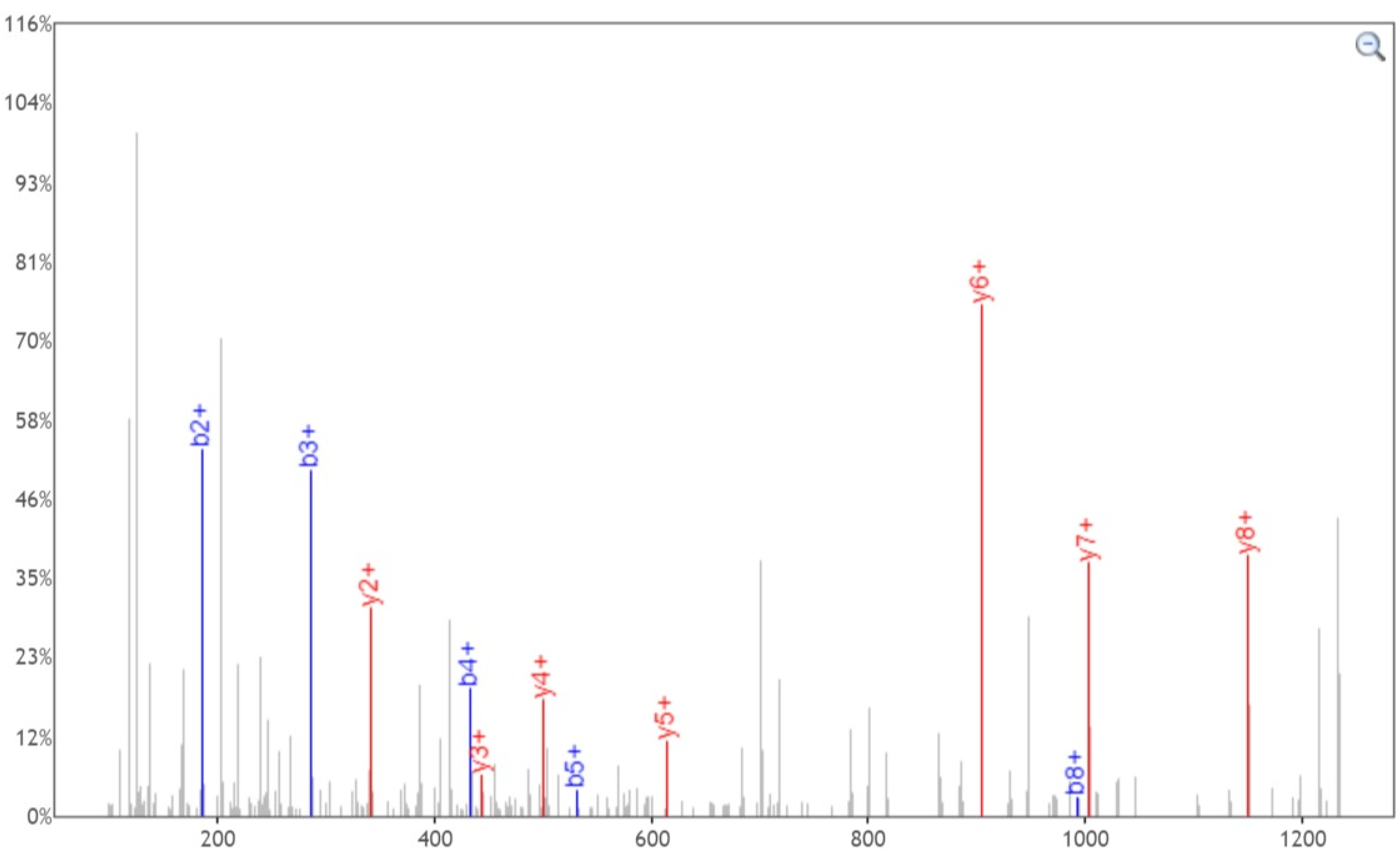


Click and drag in the plot to zoom X: Y: Zoom Out Print Enable tooltip

| b+ | # | Seq | # | y+ |
|---|---|---|---|---|
| 130.0499 | 1 | E | 11 | |
| 187.0713 | 2 | G | 10 | 1306.6062 |
| 286.1397 | 3 | V | 9 | 1249.5848 |
| 433.2082 | 4 | F | 8 | 1150.5164 |
| 532.2766 | 5 | V | 7 | 1003.4480 |
| 822.3880 | 6 | S | 6 | 904.3795 |
| 936.4309 | 7 | N | 5 | 614.2681 |
| 993.4524 | 8 | G | 4 | 500.2252 |
| 1094.5000 | 9 | T | 3 | 443.2037 |
| 1231.5590 | 10 | H | 2 | 342.1561 |
| | 11 | W | 1 | 205.0972 |

[Click] to move table

Static Modifications:
C: 57.02146

Variable Modifications:
S: 203.079373
