## Supplementary material for "Site-specific O-glycosylation analysis of SARS-CoV-2 spike protein produced in insect and human cells": Dataset S7: S1097-2-N.pdf

### Fragment Ion View

Project name: 20200330COV19
Sample name: 20200416_COV19-double_1
Created date: 2020-04-18 15:22:56.0
Search name:
Activation type: CID

Peptide sequence: R.EGVFVSN(2.988261)GTHWF.V
Scan number: 53387
Calculated M+H: 1382.6261
Measured M+H: 1382.631
Charge state: 2
File name: 20200416_COV19-double_1
search type: light

Ions:
a 1+ 2+ 3+
b 1+ 2+ 3+
c 1+ 2+ 3+
x 1+ 2+ 3+
y 1+ 2+ 3+
z 1+ 2+ 3+
[Deselect All]

Neutral Loss:
$H_2O$ (o)
$NH_3$ (*)
$H_3PO_4$ (^)
Precursor (#)
Precursor with reduced charges

Mass Type:
Mono Avg

Mass Tol: 20
Dalton PPM

Update

Peak Assignment:
Most Intense
Nearest Match
Peak Detect
Precursor Removal

Peak Labels:
Ion m/z
None

Width: 750

Height: 450

**EGVFVSNGTHWF**, MH+ 1382.6261, m/z 691.8167
File: 20200416_COV19-double_1, Scan: 53387, Precursor m/z: 691.81915, Charge: 2

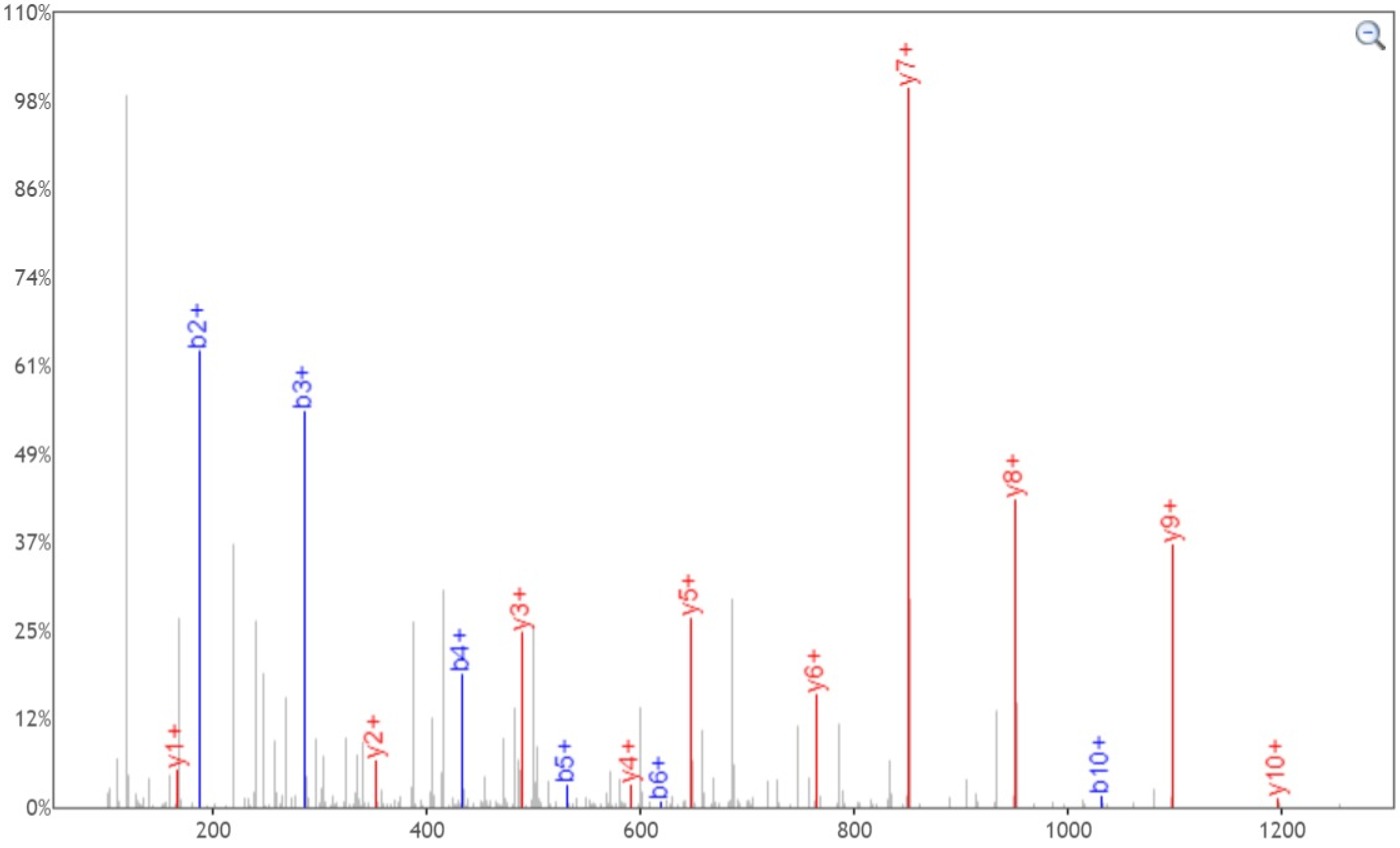


Click and drag in the plot to zoom X: Y: Zoom Out Print Enable tooltip

| b+ | # | Seq | # | y+ |
|---|---|---|---|---|
| 130.0499 | 1 | E | 12 | |
| 187.0713 | 2 | G | 11 | 1253.5835 |
| 286.1397 | 3 | V | 10 | 1196.5621 |
| 433.2082 | 4 | F | 9 | 1097.4937 |
| 532.2766 | 5 | V | 8 | 950.4253 |
| 619.3086 | 6 | S | 7 | 851.3568 |
| 736.3398 | 7 | N | 6 | 764.3248 |
| 793.3613 | 8 | G | 5 | 647.2936 |
| 894.4089 | 9 | T | 4 | 590.2722 |
| 1031.4678 | 10 | H | 3 | 489.2245 |
| 1217.5472 | 11 | W | 2 | 352.1656 |
| | 12 | F | 1 | 166.0863 |

[Click] to move table

Static Modifications:
C: 57.02146

Variable Modifications:
N: 2.988261
