## Supplementary material for "Site-specific O-glycosylation analysis of SARS-CoV-2 spike protein produced in insect and human cells": Dataset S7: T63-2-N.pdf

### Fragment Ion View

Project name: 20200330COV19
Sample name: 20200416_COV19-double_1
Created date: 2020-04-18 15:22:56.0
Search name:
Activation type: CID

Peptide sequence: R.SSVLHSTQDLFLPFFSN(2.988261)VTW.F
Scan number: 56809
Calculated M+H: 2328.1433
Measured M+H: 2328.1624
Charge state: 2
File name: 20200416_COV19-double_1
search type: light

Ions:
a 1+ 2+ 3+
b 1+ 2+ 3+
c 1+ 2+ 3+
x 1+ 2+ 3+
y 1+ 2+ 3+
z 1+ 2+ 3+
[Deselect All]

Neutral Loss:
$H_2O$ (o)
$NH_3$ (*)
$H_3PO_4$ (^)
Precursor (#)
Precursor with reduced charges

Mass Type: Mono Avg

Mass Tol: 20

Dalton PPM

Update

Peak Assignment:
Most Intense
Nearest Match
Peak Detect
Precursor Removal

Peak Labels:
Ion m/z
None

Width: 750

Height: 450

SSVLHSTQDLFLPFFSNVTW, MH+ 2328.1433, m/z 1164.5753
File: 20200416_COV19-double_1, Scan: 56809, Precursor m/z: 1164.5848, Charge: 2

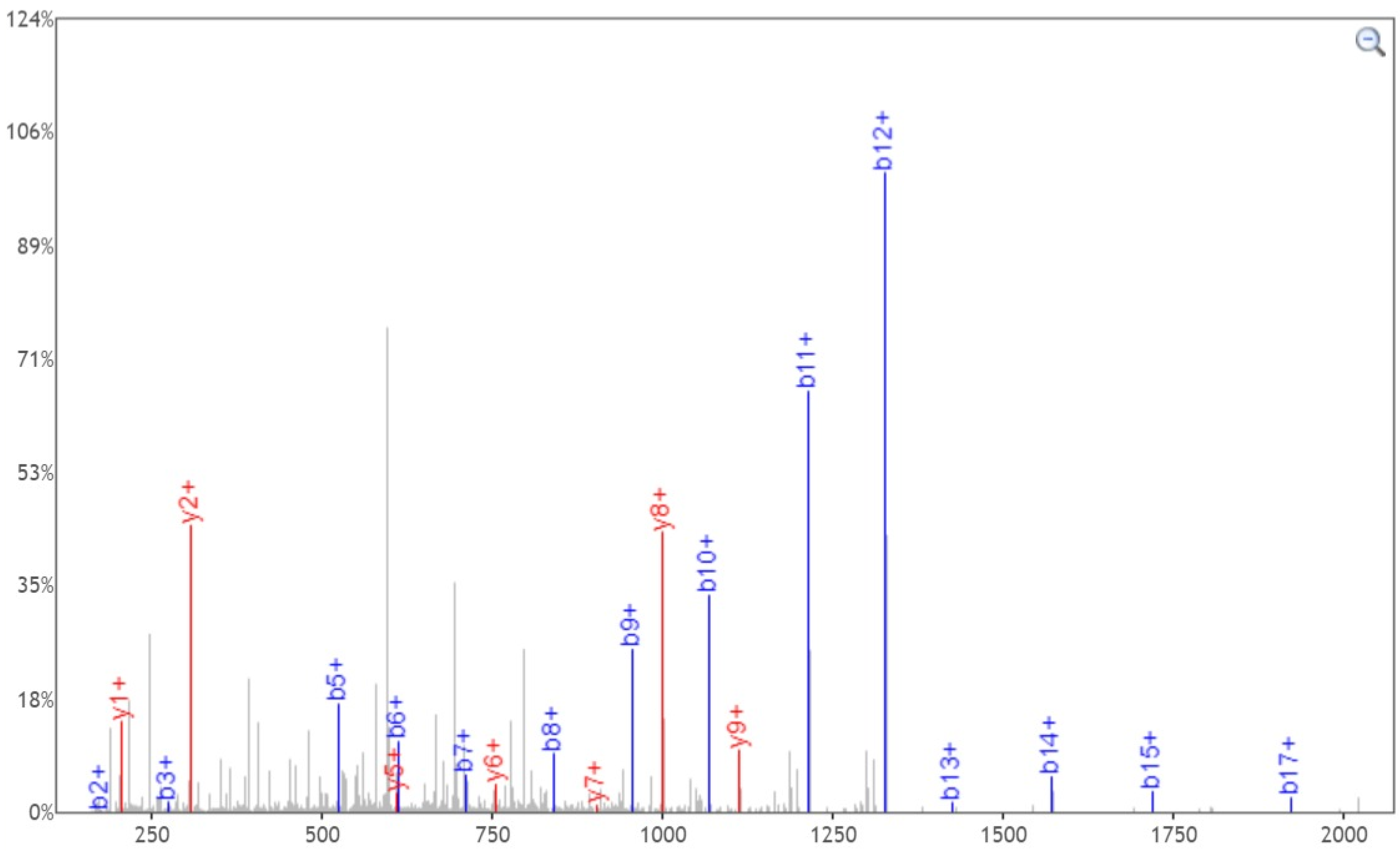


Click and drag in the plot to zoom X: Y: Zoom Out Print Enable tooltip

| b+ | # | Seq | # | y+ |
|---|---|---|---|---|
| 88.0393 | 1 | S | 20 | |
| 175.0713 | 2 | S | 19 | 2241.1112 |
| 274.1397 | 3 | V | 18 | 2154.0792 |
| 387.2238 | 4 | L | 17 | 2055.0108 |
| 524.2827 | 5 | H | 16 | 1941.9267 |
| 611.3148 | 6 | S | 15 | 1804.8678 |
| 712.3624 | 7 | T | 14 | 1717.8358 |
| 840.4210 | 8 | Q | 13 | 1616.7881 |
| 955.4480 | 9 | D | 12 | 1488.7295 |
| 1068.5320 | 10 | L | 11 | 1373.7026 |
| 1215.6004 | 11 | F | 10 | 1260.6185 |
| 1328.6845 | 12 | L | 9 | 1113.5501 |
| 1425.7373 | 13 | P | 8 | 1000.4661 |
| 1572.8057 | 14 | F | 7 | 903.4133 |
| 1719.8741 | 15 | F | 6 | 756.3449 |
| 1806.9061 | 16 | S | 5 | 609.2765 |
| 1923.9373 | 17 | N | 4 | 522.2444 |
| 2023.0057 | 18 | V | 3 | 405.2132 |
| 2124.0534 | 19 | T | 2 | 306.1448 |
| | 20 | W | 1 | 205.0972 |

[Click] to move table

Static Modifications:
C: 57.02146

Variable Modifications:
N: 2.988261
