## Supplementary material for "Site-specific O-glycosylation analysis of SARS-CoV-2 spike protein produced in insect and human cells": Dataset S7: T63-2-Y.pdf

### Fragment Ion View

Project name: 20200330COV19
Sample name: 20200416_COV19-double_1
Created date: 2020-04-18 15:22:56.0
Search name:
Activation type: CID

Peptide sequence: R.SSVLHSTQDLFLPFFSNVT(203.079373)W.F
Scan number: 56688
Calculated M+H: 2528.2344
Measured M+H: 2528.2502
Charge state: 2
File name: 20200416_COV19-double_1
search type: light

Ions:
a 1+ 2+ 3+
b 1+ 2+ 3+
c 1+ 2+ 3+
x 1+ 2+ 3+
y 1+ 2+ 3+
z 1+ 2+ 3+
[Deselect All]

Neutral Loss:
$H_2O$ (o)
$NH_3$ (*)
$H_3PO_4$ (^)
Precursor (#)
Precursor with reduced charges

Mass Type:
Mono Avg

Mass Tol: 20
Dalton PPM

Update

Peak Assignment:
Most Intense
Nearest Match
Peak Detect
Precursor Removal

Peak Labels:
Ion m/z
None

Width: 750

Height: 450

SSVLHSTQDLFLPFFSNVTW, MH+ 2528.2344, m/z 1264.6208
File: 20200416_COV19-double_1, Scan: 56688, Precursor m/z: 1264.6288, Charge: 2

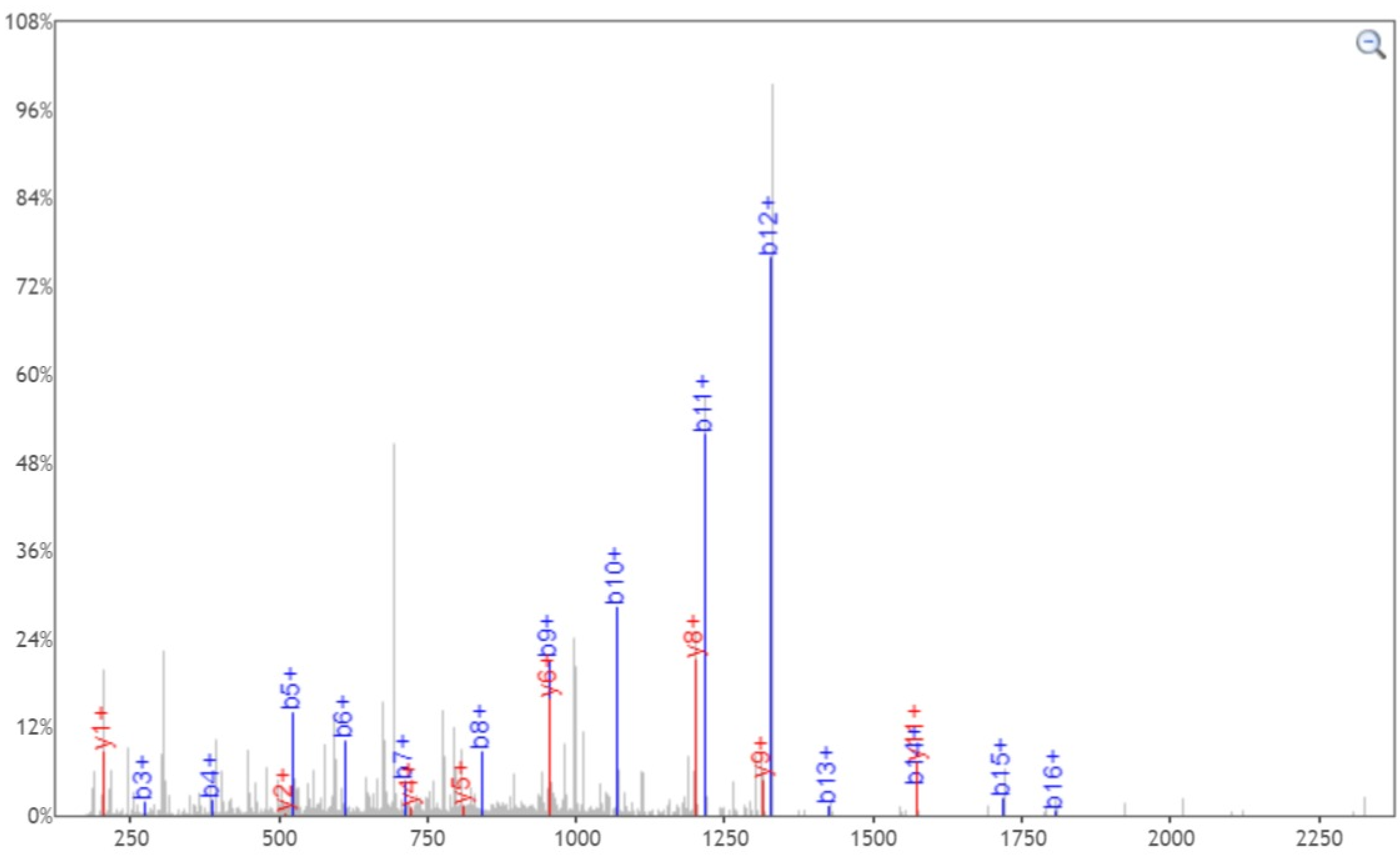


Click and drag in the plot to zoom X: Y: Zoom Out Print Enable tooltip

| b+ | # | Seq | # | y+ |
|---|---|---|---|---|
| 88.0393 | 1 | S | 20 | |
| 175.0713 | 2 | S | 19 | 2441.2024 |
| 274.1397 | 3 | V | 18 | 2354.1703 |
| 387.2238 | 4 | L | 17 | 2255.1019 |
| 524.2827 | 5 | H | 16 | 2142.0178 |
| 611.3148 | 6 | S | 15 | 2004.9589 |
| 712.3624 | 7 | T | 14 | 1917.9269 |
| 840.4210 | 8 | Q | 13 | 1816.8792 |
| 955.4480 | 9 | D | 12 | 1688.8207 |
| 1068.5320 | 10 | L | 11 | 1573.7937 |
| 1215.6004 | 11 | F | 10 | 1460.7096 |
| 1328.6845 | 12 | L | 9 | 1313.6412 |
| 1425.7373 | 13 | P | 8 | 1200.5572 |
| 1572.8057 | 14 | F | 7 | 1103.5044 |
| 1719.8741 | 15 | F | 6 | 956.4360 |
| 1806.9061 | 16 | S | 5 | 809.3676 |
| 1920.9490 | 17 | N | 4 | 722.3355 |
| 2020.0175 | 18 | V | 3 | 608.2926 |
| 2324.1445 | 19 | T | 2 | 509.2242 |
| | 20 | W | 1 | 205.0972 |

[Click] to move table

Static Modifications:
C: 57.02146

Variable Modifications:
T: 203.079373
