## Supplementary material for "Site-specific O-glycosylation analysis of SARS-CoV-2 spike protein produced in insect and human cells": Dataset S7: T76-2-Y.pdf

### Fragment Ion View

Project name: 20200330COV19
Sample name: 20200416_COV19-double_1
Created date: 2020-04-18 15:22:56.0
Search name:
Activation type: CID

Peptide sequence: F.HAIHVSGTNGT(203.079373)K.R
Scan number: 7068
Calculated M+H: 1424.7129
Measured M+H: 1424.7184
Charge state: 2
File name: 20200416_COV19-double_1
search type: light

Ions:
a 1+ 2+ 3+
b 1+ 2+ 3+
c 1+ 2+ 3+
x 1+ 2+ 3+
y 1+ 2+ 3+
z 1+ 2+ 3+
[Deselect All]

Neutral Loss:
$H_2O$ (o)
$NH_3$ (*)
$H_3PO_4$ (^)
Precursor (#)
Precursor with reduced charges

Mass Type:
Mono Avg

Mass Tol: 20
Dalton PPM

Update

Peak Assignment:
Most Intense
Nearest Match
Peak Detect
Precursor Removal

Peak Labels:
Ion m/z
None

Width: 750

Height: 450

**HAIHVSGTNGTK**, MH+ 1424.7128, m/z 712.8601
File: 20200416_COV19-double_1, Scan: 7068, Precursor m/z: 712.86285, Charge: 2

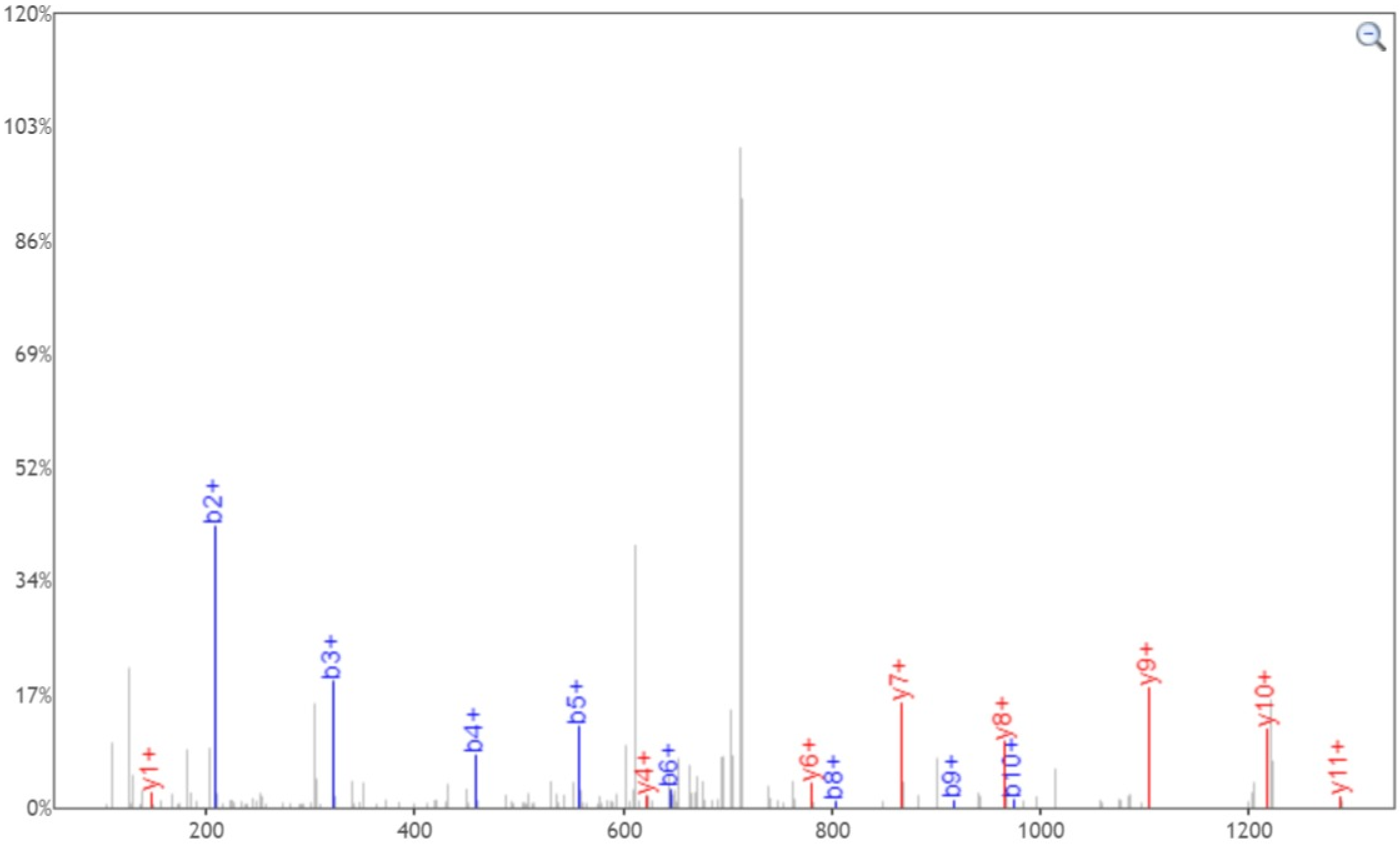


Click and drag in the plot to zoom X: Y: Zoom Out Print Enable tooltip

| b+ | # | Seq | # | y+ |
|---|---|---|---|---|
| 138.0662 | 1 | H | 12 | |
| 209.1033 | 2 | A | 11 | 1287.6539 |
| 322.1874 | 3 | I | 10 | 1216.6168 |
| 459.2463 | 4 | H | 9 | 1103.5327 |
| 558.3147 | 5 | V | 8 | 966.4738 |
| 645.3467 | 6 | S | 7 | 867.4054 |
| 702.3682 | 7 | G | 6 | 780.3734 |
| 803.4159 | 8 | T | 5 | 723.3519 |
| 917.4588 | 9 | N | 4 | 622.3042 |
| 974.4803 | 10 | G | 3 | 508.2613 |
| 1278.6073 | 11 | T | 2 | 451.2399 |
| | 12 | K | 1 | 147.1128 |

[Click] to move table

Static Modifications:
C: 57.02146

Variable Modifications:
T: 203.079373
