## Supplementary material for "Site-specific O-glycosylation analysis of SARS-CoV-2 spike protein produced in insect and human cells": Dataset S7: T167-1-N.pdf

### Fragment Ion View

Project name: 20200330COV19
Sample name: 20200416_COV19-double_1
Created date: 2020-04-18 15:22:56.0
Search name:
Activation type: CID

Peptide sequence: R.VYSSANN(2.988261)CTFEYVSQPFLMDLEGK.Q
Scan number: 56061
Calculated M+H: 2802.2522
Measured M+H: 2802.2622
Charge state: 3
File name: 20200416_COV19-double_1
search type: light

**Ions:**
a 1+ 2+ 3+
b 1+ 2+ 3+
c 1+ 2+ 3+
x 1+ 2+ 3+
y 1+ 2+ 3+
z 1+ 2+ 3+
[Deselect All]

**Neutral Loss:**
$H_2O$ (o)
$NH_3$ (*)
$H_3PO_4$ (^)
Precursor (#)
Precursor with reduced charges

Mass Type: Mono Avg

Mass Tol: 20

Dalton PPM

Update

Peak Assignment:
Most Intense
Nearest Match
Peak Detect
Precursor Removal

Peak Labels:
Ion m/z
None

Width: 750

Height: 450

VYSSANNCTFEYVSQPFLMDLEGK, MH+ 2802.2523, m/z 934.7556
File: 20200416_COV19-double_1, Scan: 56061, Precursor m/z: 934.7589, Charge: 3

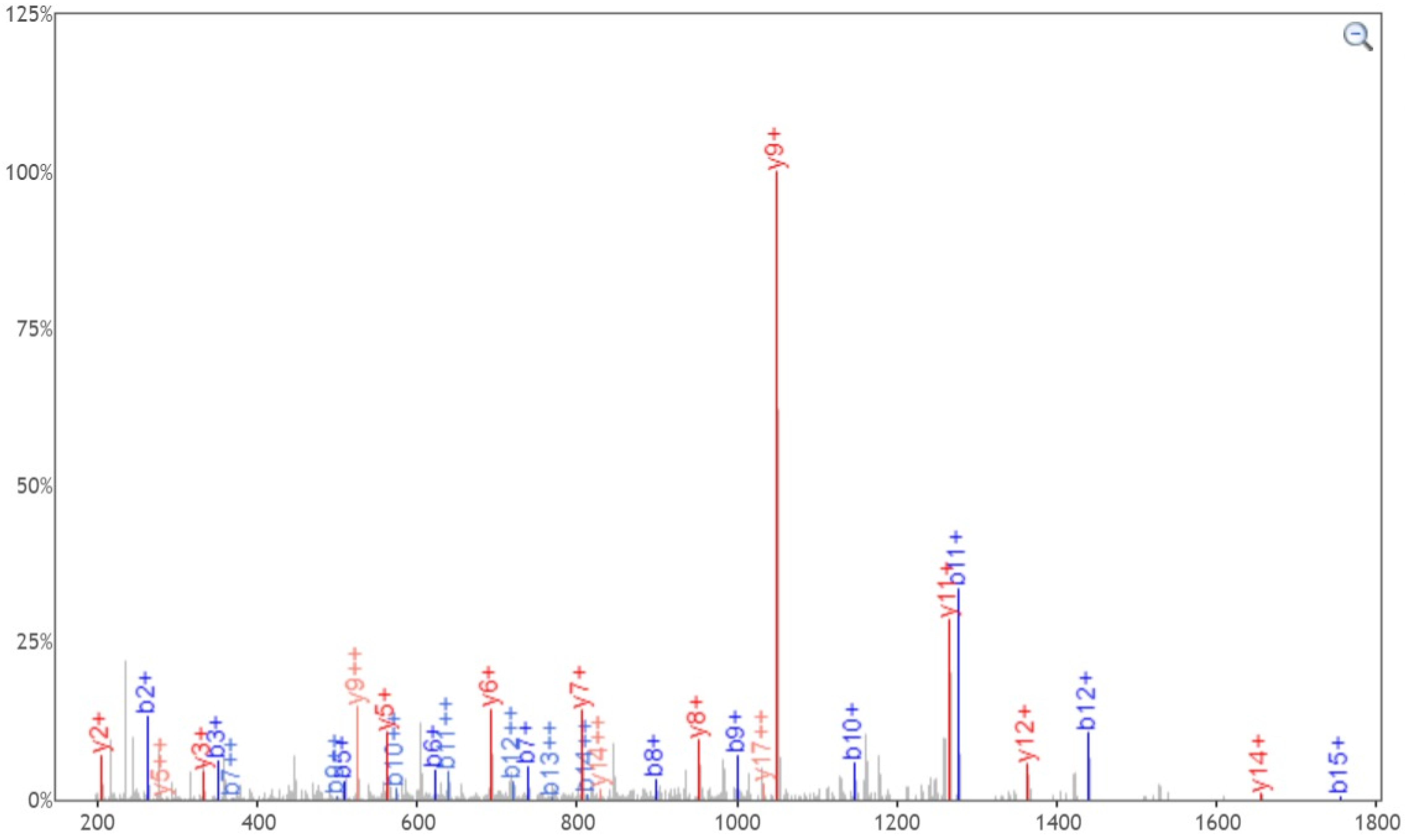


Click and drag in the plot to zoom X: Y: Zoom Out Print Enable tooltip

| b+ | b2+ | # | Seq | # | y+ | y2+ |
|---|---|---|---|---|---|---|
| 100.0757 | 50.5415 | 1 | V | 24 | | |
| 263.1390 | 132.0731 | 2 | Y | 23 | 2703.1839 | 1352.0956 |
| 350.1710 | 175.5892 | 3 | S | 22 | 2540.1206 | 1270.5639 |
| 437.2031 | 219.1052 | 4 | S | 21 | 2453.0885 | 1227.0479 |
| 508.2402 | 254.6237 | 5 | A | 20 | 2366.0565 | 1183.5319 |
| 622.2831 | 311.6452 | 6 | N | 19 | 2295.0194 | 1148.0133 |
| 739.3143 | 370.1608 | 7 | N | 18 | 2180.9765 | 1090.9919 |
| 899.3449 | 450.1761 | 8 | C | 17 | 2063.9453 | 1032.4763 |
| 1000.3926 | 500.7000 | 9 | T | 16 | 1903.9146 | 952.4610 |
| 1147.4610 | 574.2342 | 10 | F | 15 | 1802.8669 | 901.9371 |
| 1276.5036 | 638.7555 | 11 | E | 14 | 1655.7985 | 828.4029 |
| 1439.5670 | 720.2871 | 12 | Y | 13 | 1526.7559 | 763.8816 |
| 1538.6354 | 769.8213 | 13 | V | 12 | 1363.6926 | 682.3499 |
| 1625.6674 | 813.3373 | 14 | S | 11 | 1264.6242 | 632.8157 |
| 1753.7260 | 877.3666 | 15 | Q | 10 | 1177.5922 | 589.2997 |
| 1850.7787 | 925.8930 | 16 | P | 9 | 1049.5336 | 525.2704 |
| 1997.8472 | 999.4272 | 17 | F | 8 | 952.4808 | 476.7441 |
| 2110.9312 | 1055.9693 | 18 | L | 7 | 805.4124 | 403.2098 |
| 2241.9717 | 1121.4895 | 19 | M | 6 | 692.3284 | 346.6678 |
| 2356.9987 | 1179.0030 | 20 | D | 5 | 561.2879 | 281.1476 |
| 2470.0827 | 1235.5450 | 21 | L | 4 | 446.2609 | 223.6341 |
| 2599.1253 | 1300.0663 | 22 | E | 3 | 333.1769 | 167.0921 |
| 2656.1468 | 1328.5770 | 23 | G | 2 | 204.1343 | 102.5708 |
| | | 24 | K | 1 | 147.1128 | 74.0600 |

[Click] to move table

**Static Modifications:**
**C: 57.02146**

**Variable Modifications:**
**N: 2.988261**
