## Supplementary material for "Site-specific O-glycosylation analysis of SARS-CoV-2 spike protein produced in insect and human cells": Dataset S7: T167-1-Y.pdf

### Fragment Ion View

Project name: 20200330COV19
Sample name: 20200416_COV19-double_1
Created date: 2020-04-18 15:22:56.0
Search name:
Activation type: CID

Peptide sequence: R.VYSSANNCT(203.079373)FEYVSQPFLMDLEGK.Q
Scan number: 55874
Calculated M+H: 3002.3435
Measured M+H: 3002.3613
Charge state: 3
File name: 20200416_COV19-double_1
search type: light

Ions:
a 1+ 2+ 3+
b [x] 1+ [x] 2+ 3+
c 1+ 2+ 3+
x 1+ 2+ 3+
y [x] 1+ [x] 2+ 3+
z 1+ 2+ 3+
[Deselect All]

Neutral Loss:
$H_2O$ (o)
$NH_3$ (*)
$H_3PO_4$ (^)
Precursor (#)
Precursor with reduced charges

Mass Type: (•) Mono ( ) Avg

Mass Tol: 20
( ) Dalton (•) PPM

Update

Peak Assignment:
(•) Most Intense
( ) Nearest Match
Peak Detect
Precursor Removal

Peak Labels:
(•) Ion ( ) m/z
( ) None

Width: 750

Height: 450

VYSSANNCTFEYVSQPFLMDLEGK, MH+ 3002.3434, m/z 1001.4527
File: 20200416_COV19-double_1, Scan: 55874, Precursor m/z: 1001.4586, Charge: 3

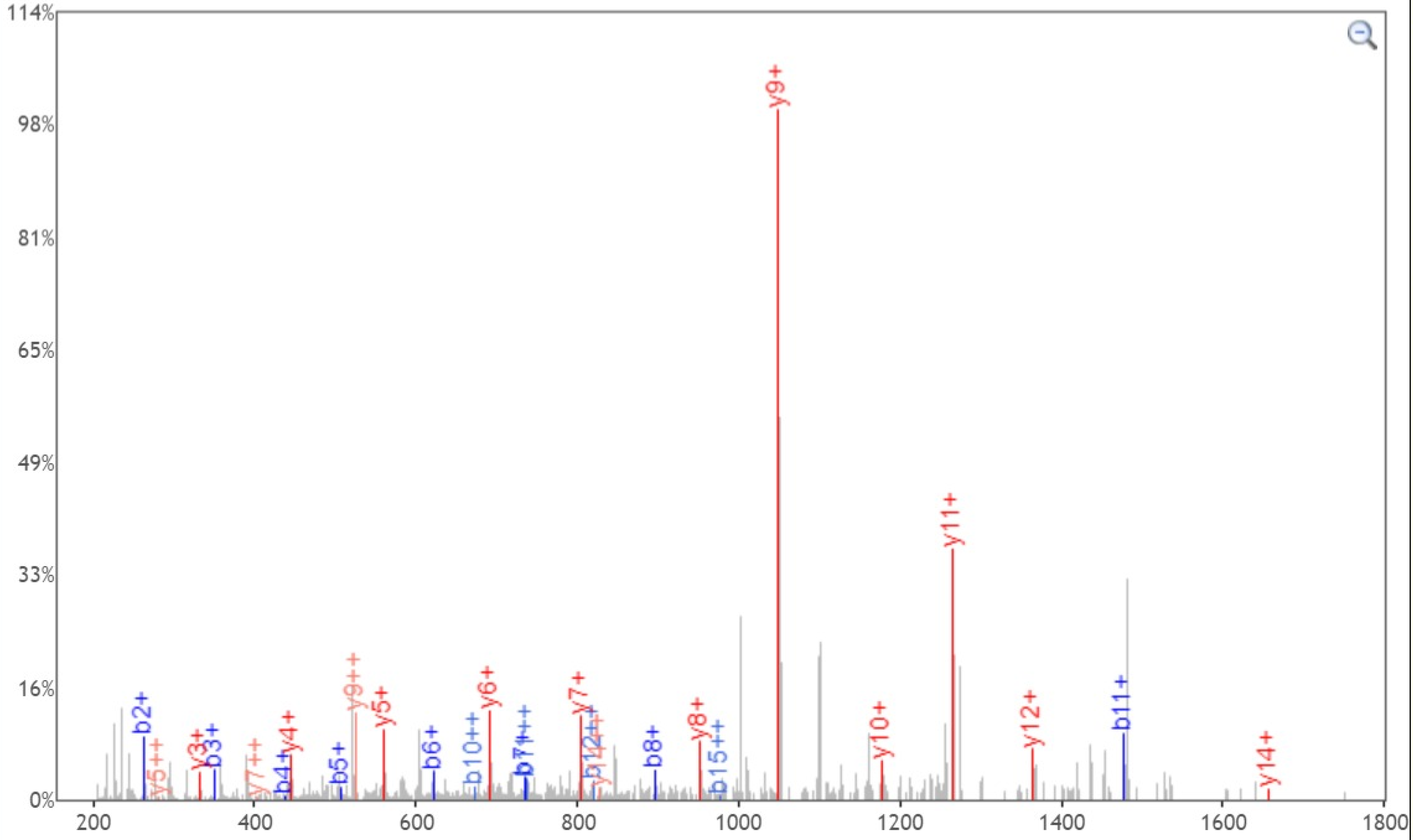


Click and drag in the plot to zoom X: [x] Y: [ ] Zoom Out Print [ ] Enable tooltip

| b+ | b2+ | # | Seq | # | y+ | y2+ |
|---|---|---|---|---|---|---|
| 100.0757 | 50.5415 | 1 | V | 24 | | |
| 263.1390 | 132.0731 | 2 | Y | 23 | 2903.2750 | 1452.1411 |
| 350.1710 | 175.5892 | 3 | S | 22 | 2740.2117 | 1370.6095 |
| 437.2031 | 219.1052 | 4 | S | 21 | 2653.1796 | 1327.0935 |
| 508.2402 | 254.6237 | 5 | A | 20 | 2566.1476 | 1283.5774 |
| 622.2831 | 311.6452 | 6 | N | 19 | 2495.1105 | 1248.0589 |
| 736.3260 | 368.6667 | 7 | N | 18 | 2381.0676 | 1191.0374 |
| 896.3567 | 448.6820 | 8 | C | 17 | 2267.0246 | 1134.0160 |
| 1200.4837 | 600.7455 | 9 | T | 16 | 2106.9940 | 1054.0006 |
| 1347.5522 | 674.2797 | 10 | F | 15 | 1802.8669 | 901.9371 |
| 1476.5947 | 738.8010 | 11 | E | 14 | 1655.7985 | 828.4029 |
| 1639.6581 | 820.3327 | 12 | Y | 13 | 1526.7559 | 763.8816 |
| 1738.7265 | 869.8669 | 13 | V | 12 | 1363.6926 | 682.3499 |
| 1825.7585 | 913.3829 | 14 | S | 11 | 1264.6242 | 632.8157 |
| 1953.8171 | 977.4122 | 15 | Q | 10 | 1177.5922 | 589.2997 |
| 2050.8699 | 1025.9386 | 16 | P | 9 | 1049.5336 | 525.2704 |
| 2197.9383 | 1099.4728 | 17 | F | 8 | 952.4808 | 476.7441 |
| 2311.0223 | 1156.0148 | 18 | L | 7 | 805.4124 | 403.2098 |
| 2442.0628 | 1221.5350 | 19 | M | 6 | 692.3284 | 346.6678 |
| 2557.0898 | 1279.0485 | 20 | D | 5 | 561.2879 | 281.1476 |
| 2670.1738 | 1335.5906 | 21 | L | 4 | 446.2609 | 223.6341 |
| 2799.2164 | 1400.1118 | 22 | E | 3 | 333.1769 | 167.0921 |
| 2856.2379 | 1428.6226 | 23 | G | 2 | 204.1343 | 102.5708 |
| | | 24 | K | 1 | 147.1128 | 74.0600 |

[Click] to move table

Static Modifications:
C: 57.02146

Variable Modifications:
T: 203.079373
