## Supplementary material for "Site-specific O-glycosylation analysis of SARS-CoV-2 spike protein produced in insect and human cells": Dataset S7: T236-1-Y.pdf

### Fragment Ion View

Project name: 20200330COV19
Sample name: 20200416_COV19-double_1
Created date: 2020-04-18 15:22:56.0
Search name:
Activation type: CID

Peptide sequence: F.SALEPLVDLPIGINIT(203.079373).R
Scan number: 56301
Calculated M+H: 1868.0262
Measured M+H: 1868.0326
Charge state: 2
File name: 20200416_COV19-double_1
search type: light

Ions:
a 1+ 2+ 3+
b 1+ 2+ 3+
c 1+ 2+ 3+
x 1+ 2+ 3+
y 1+ 2+ 3+
z 1+ 2+ 3+
[Deselect All]

Neutral Loss:
$H_2O$ (o)
$NH_3$ (*)
$H_3PO_4$ (^)
Precursor (#)
Precursor with reduced charges

Mass Type: Mono Avg

Mass Tol: 20

Dalton PPM

Update

Peak Assignment:
Most Intense
Nearest Match
Peak Detect
Precursor Removal

Peak Labels:
Ion m/z
None

Width: 750

Height: 450

SALEPLVDLPIGINIT, MH+ 1868.0263, m/z 934.5168
File: 20200416_COV19-double_1, Scan: 56301, Precursor m/z: 934.51996, Charge: 2

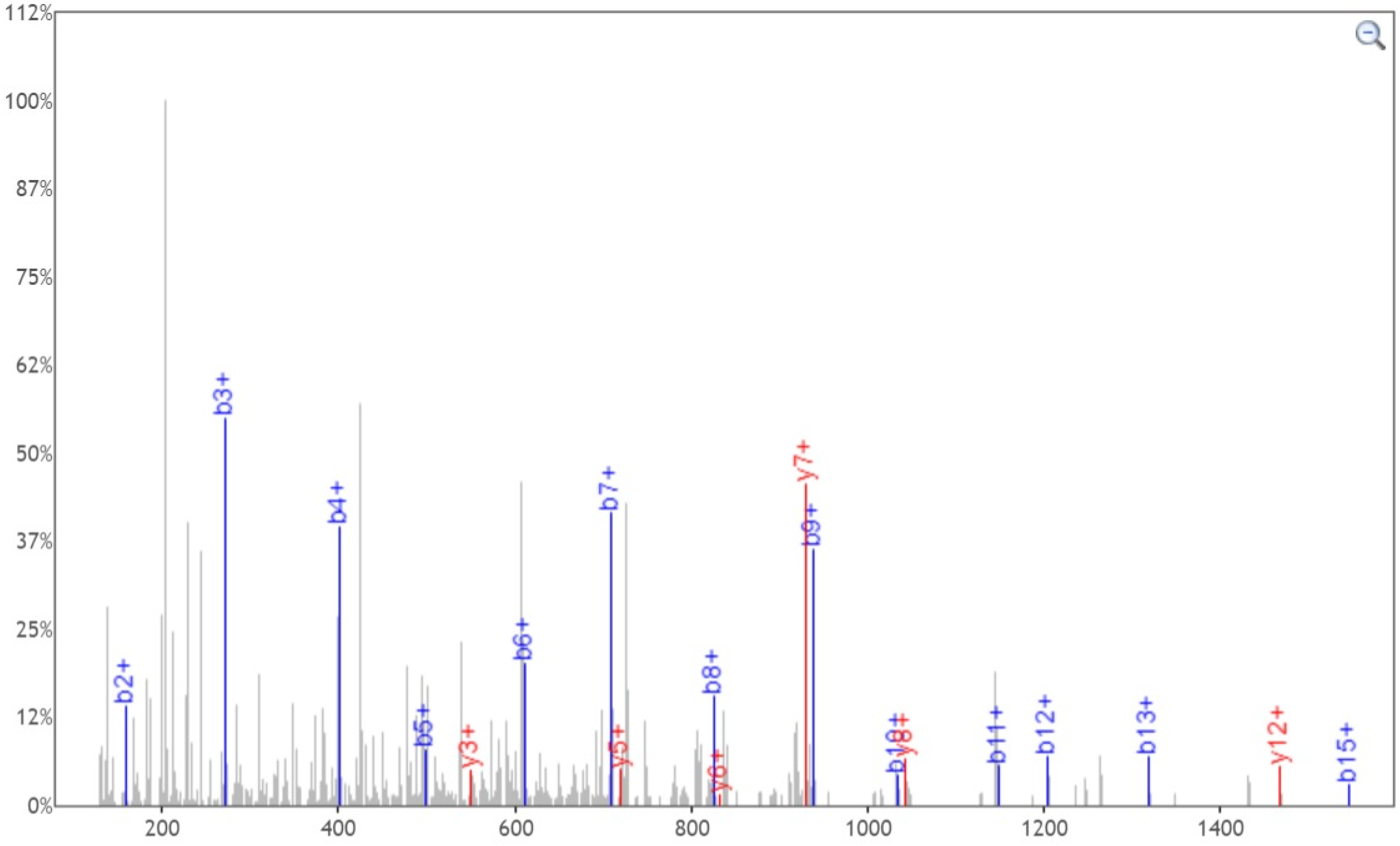


Click and drag in the plot to zoom X: Y: Zoom Out Print Enable tooltip

| b+ | # | Seq | # | y+ |
|---|---|---|---|---|
| 88.0393 | 1 | S | 16 | |
| 159.0764 | 2 | A | 15 | 1780.9943 |
| 272.1605 | 3 | L | 14 | 1709.9571 |
| 401.2031 | 4 | E | 13 | 1596.8731 |
| 498.2558 | 5 | P | 12 | 1467.8305 |
| 611.3399 | 6 | L | 11 | 1370.7777 |
| 710.4083 | 7 | V | 10 | 1257.6937 |
| 825.4353 | 8 | D | 9 | 1158.6252 |
| 938.5193 | 9 | L | 8 | 1043.5983 |
| 1035.5721 | 10 | P | 7 | 930.5142 |
| 1148.6562 | 11 | I | 6 | 833.4615 |
| 1205.6776 | 12 | G | 5 | 720.3774 |
| 1318.7617 | 13 | I | 4 | 663.3559 |
| 1432.8046 | 14 | N | 3 | 550.2719 |
| 1545.8887 | 15 | I | 2 | 436.2290 |
| | 16 | T | 1 | 323.1449 |

[Click] to move table

Static Modifications:
C: 57.02146

Variable Modifications:
T: 203.079373
