## Supplementary material for "Site-specific O-glycosylation analysis of SARS-CoV-2 spike protein produced in insect and human cells": Dataset S7: T236-2-Y.pdf

### Fragment Ion View

Project name: 20200330COV19
Sample name: 20200416_COV19-double_1
Created date: 2020-04-18 15:22:56.0
Search name:
Activation type: CID

Peptide sequence: L.VDLPIGINIT(203.079373).R
Scan number: 53501
Calculated M+H: 1257.6936
Measured M+H: 1257.6973
Charge state: 2
File name: 20200416_COV19-double_1
search type: light

**Ions:**
a ☐1+ ☐2+ ☐3+
b ☑1+ ☐2+ ☐3+
c ☐1+ ☐2+ ☐3+
x ☐1+ ☐2+ ☐3+
y ☑1+ ☐2+ ☐3+
z ☐1+ ☐2+ ☐3+
[Deselect All]

**Neutral Loss:**
☐ $H_2O$ (o)
☐ $NH_3$ (*)
☐ $H_3PO_4$ (^)
☐ Precursor (#)
☐ Precursor with reduced charges

Mass Type: ◉Mono ○Avg

Mass Tol: 20

○Dalton ◉PPM

Update

Peak Assignment:
◉Most Intense
○Nearest Match
☐Peak Detect
☐Precursor Removal

Peak Labels:
◉Ion ○m/z
○None

Width: 750

Height: 450

VDLPIGINIT, MH+ 1257.6937, m/z 629.3505
File: 20200416_COV19-double_1, Scan: 53501, Precursor m/z: 629.3523, Charge: 2

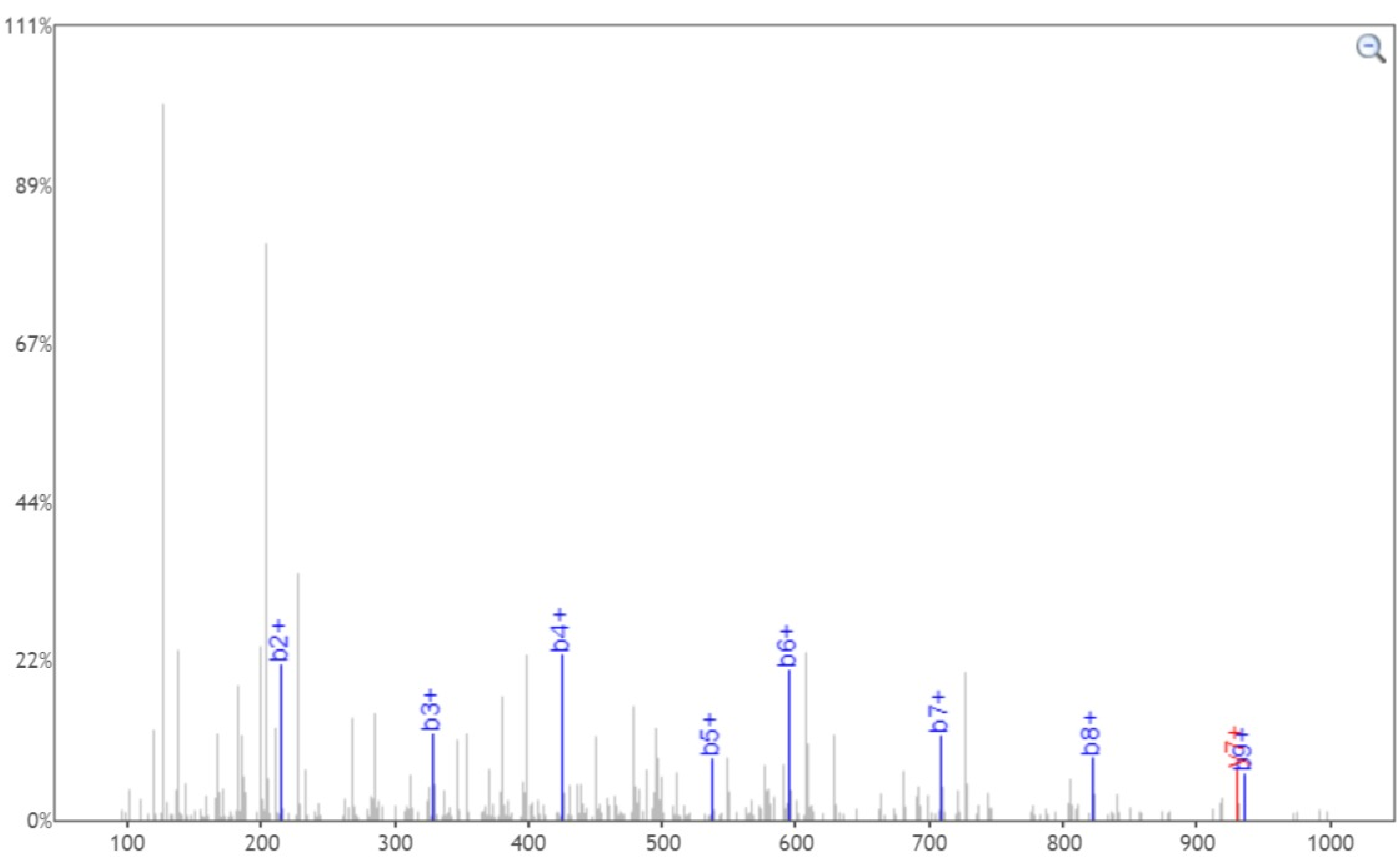


Click and drag in the plot to zoom X:☑ Y:☐ Zoom Out Print ☐Enable tooltip

| b+ | # | Seq | # | y+ |
|---|---|---|---|---|
| 100.0757 | 1 | V | 10 | |
| 215.1026 | 2 | D | 9 | 1158.6252 |
| 328.1867 | 3 | L | 8 | 1043.5983 |
| 425.2395 | 4 | P | 7 | 930.5142 |
| 538.3235 | 5 | I | 6 | 833.4615 |
| 595.3450 | 6 | G | 5 | 720.3774 |
| 708.4291 | 7 | I | 4 | 663.3559 |
| 822.4720 | 8 | N | 3 | 550.2719 |
| 935.5560 | 9 | I | 2 | 436.2290 |
| | 10 | T | 1 | 323.1449 |

[Click] to move table

Static Modifications:
C: 57.02146

Variable Modifications:
T: 203.079373
