## Supplementary material for "Site-specific O-glycosylation analysis of SARS-CoV-2 spike protein produced in insect and human cells": Dataset S7: S803-1-N.pdf

### Fragment Ion View

Project name: 20200330COV19
Sample name: 20200416_COV19-double_2
Created date: 2020-04-18 15:22:56.0
Search name:
Activation type: CID

Peptide sequence: K.DFGGFN(2.988261)FSQILPDPSK.P
Scan number: 55962
Calculated M+H: 1771.8424
Measured M+H: 1771.85
Charge state: 2
File name: 20200416_COV19-double_2
search type: light

**Ions:**
a ☐1+ ☐2+ ☐3+
b ☑1+ ☐2+ ☐3+
c ☐1+ ☐2+ ☐3+
x ☐1+ ☐2+ ☐3+
y ☑1+ ☐2+ ☐3+
z ☐1+ ☐2+ ☐3+
[Deselect All]

**Neutral Loss:**
☐ $H_2O$ (o)
☐ $NH_3$ (*)
☐ $H_3PO_4$ (^)
☐ Precursor (#)
☐ Precursor with reduced charges

Mass Type:
◉Mono ○Avg

Mass Tol: 20
○Dalton ◉PPM

Update

Peak Assignment:
◉Most Intense
○Nearest Match
☐Peak Detect
☐Precursor Removal

Peak Labels:
◉Ion ○m/z
○None

Width: 750
Height: 450

DFGGFNFSQILPDPSK, MH+ 1771.8423, m/z 886.4248
File: 20200416_COV19-double_2, Scan: 55962, Precursor m/z: 886.42865, Charge: 2

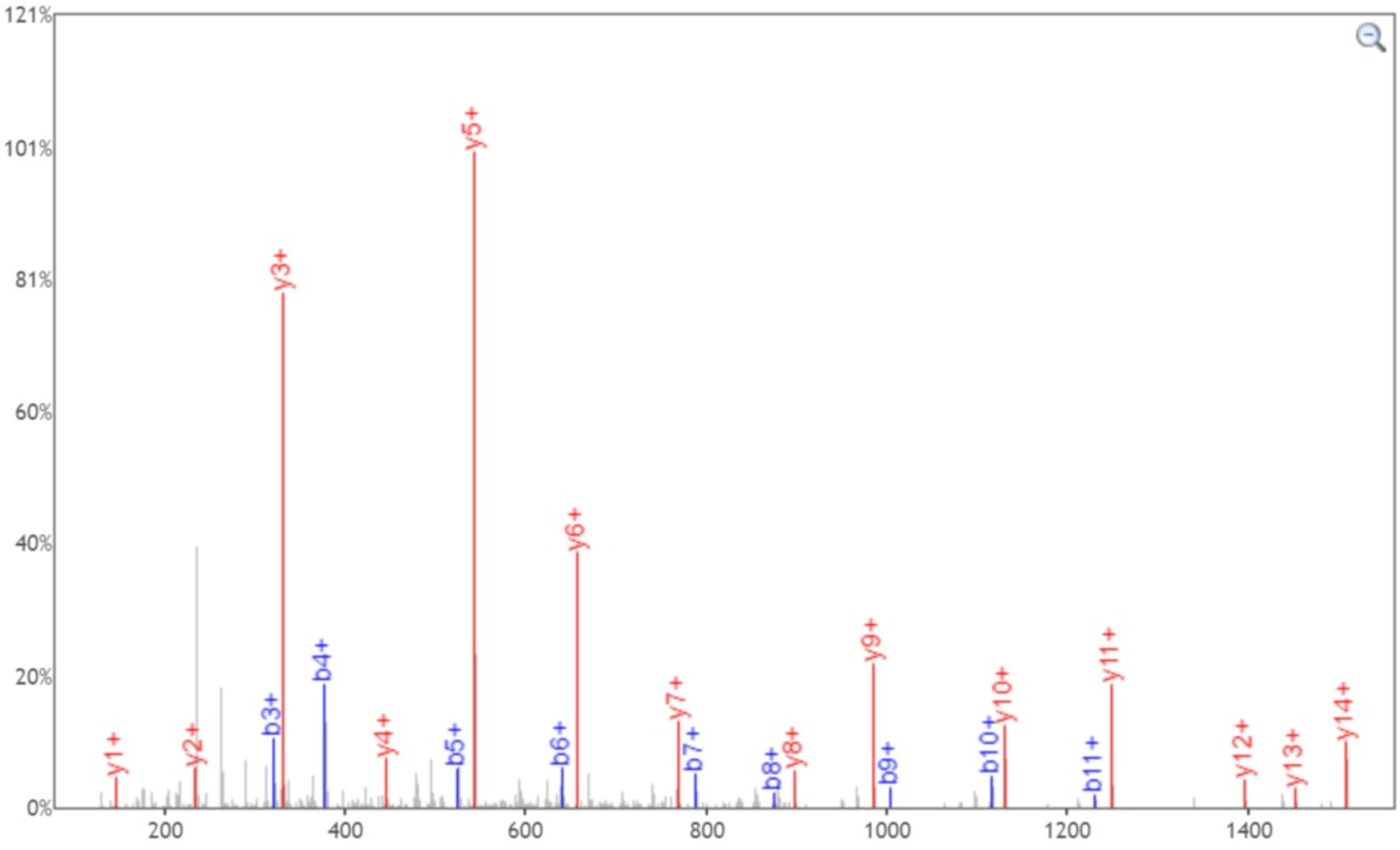


Click and drag in the plot to zoom X:☑ Y:☐ Zoom Out Print ☐Enable tooltip

| b+ | # | Seq | # | y+ |
|---|---|---|---|---|
| 116.0342 | 1 | D | 16 | |
| 263.1026 | 2 | F | 15 | 1656.8154 |
| 320.1241 | 3 | G | 14 | 1509.7470 |
| 377.1456 | 4 | G | 13 | 1452.7255 |
| 524.2140 | 5 | F | 12 | 1395.7041 |
| 641.2452 | 6 | N | 11 | 1248.6356 |
| 788.3136 | 7 | F | 10 | 1131.6045 |
| 875.3456 | 8 | S | 9 | 984.5360 |
| 1003.4042 | 9 | Q | 8 | 897.5040 |
| 1116.4882 | 10 | I | 7 | 769.4454 |
| 1229.5723 | 11 | L | 6 | 656.3614 |
| 1326.6251 | 12 | P | 5 | 543.2773 |
| 1441.6520 | 13 | D | 4 | 446.2245 |
| 1538.7048 | 14 | P | 3 | 331.1976 |
| 1625.7368 | 15 | S | 2 | 234.1448 |
| | 16 | K | 1 | 147.1128 |

[Click] to move table

Static Modifications:
C: 57.02146

Variable Modifications:
N: 2.988261
