## Supplementary material for "Site-specific O-glycosylation analysis of SARS-CoV-2 spike protein produced in insect and human cells": Dataset S7: S803-1-Y.pdf

### Fragment Ion View

Project name: 20200330COV19
Sample name: 20200416_COV19-double_2
Created date: 2020-04-18 15:22:56.0
Search name:
Activation type: CID

Peptide sequence: K.DFGGFNFS(203.079373)QILPDPSK.P
Scan number: 55675
Calculated M+H: 1971.9335
Measured M+H: 1971.9424
Charge state: 2
File name: 20200416_COV19-double_2
search type: light

Ions:
a 1+ 2+ 3+
b ☑1+ 2+ 3+
c 1+ 2+ 3+
x 1+ 2+ 3+
y ☑1+ 2+ 3+
z 1+ 2+ 3+
[Deselect All]

Neutral Loss:
- $H_2O$ (o)
- $NH_3$ (*)
- $H_3PO_4$ (^)
- Precursor (#)
- Precursor with reduced charges

Mass Type: ◉Mono ○Avg

Mass Tol: 20

○Dalton ◉PPM

Update

Peak Assignment:
- ◉Most Intense
- ○Nearest Match
- Peak Detect
- Precursor Removal

Peak Labels: ◉Ion ○m/z ○None

Width: 750

Height: 450

DFGGFNFSQILPDPSK, MH+ 1971.9334, m/z 986.4704
File: 20200416_COV19-double_2, Scan: 55675, Precursor m/z: 986.47485, Charge: 2

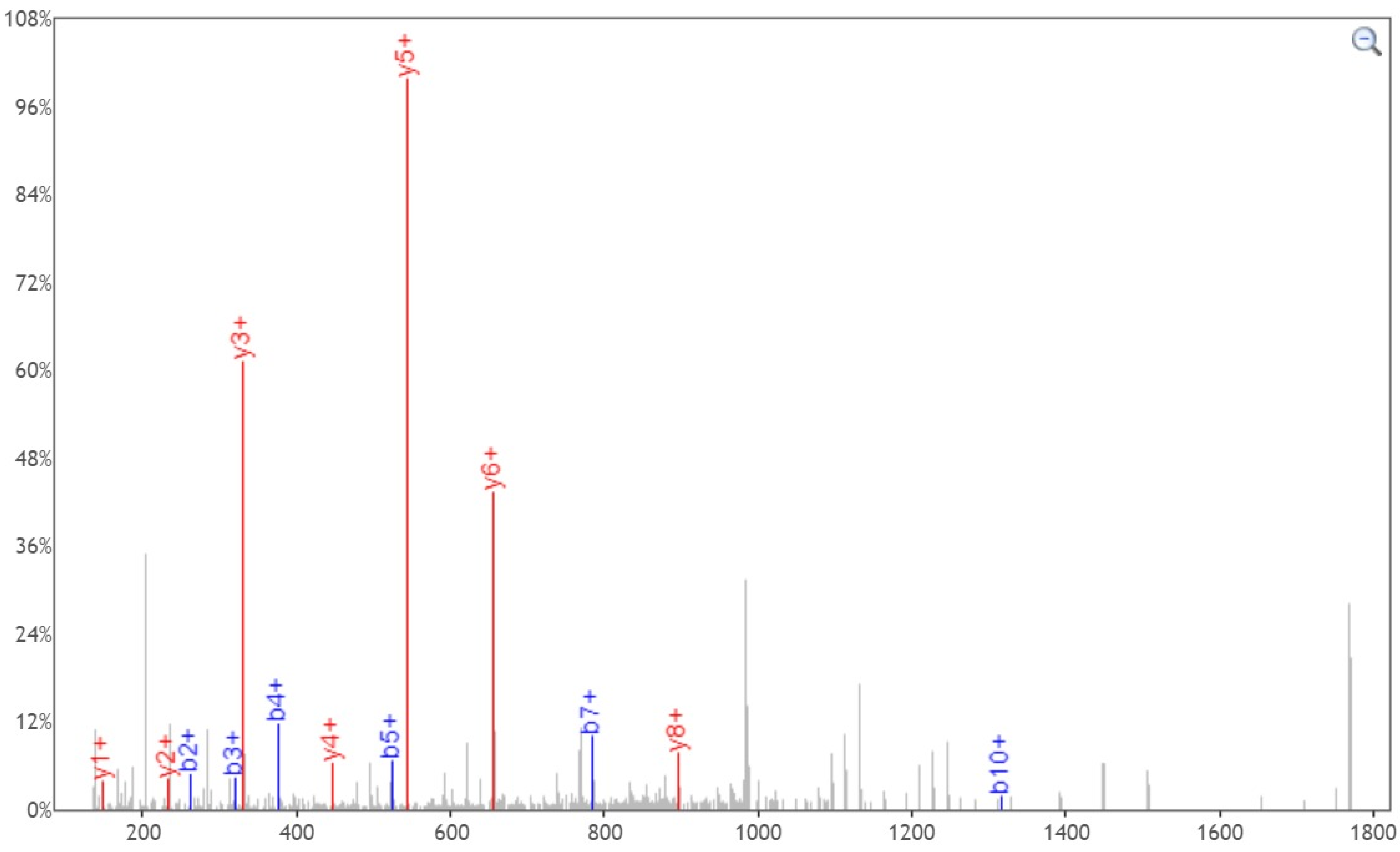


Click and drag in the plot to zoom X:☑ Y: Zoom Out Print Enable tooltip

| b+ | # | Seq | # | y+ |
|---|---|---|---|---|
| 116.0342 | 1 | D | 16 | |
| 263.1026 | 2 | F | 15 | 1856.9065 |
| 320.1241 | 3 | G | 14 | 1709.8381 |
| 377.1456 | 4 | G | 13 | 1652.8166 |
| 524.2140 | 5 | F | 12 | 1595.7952 |
| 638.2569 | 6 | N | 11 | 1448.7268 |
| 785.3253 | 7 | F | 10 | 1334.6838 |
| 1075.4367 | 8 | S | 9 | 1187.6154 |
| 1203.4953 | 9 | Q | 8 | 897.5040 |
| 1316.5794 | 10 | I | 7 | 769.4454 |
| 1429.6634 | 11 | L | 6 | 656.3614 |
| 1526.7162 | 12 | P | 5 | 543.2773 |
| 1641.7431 | 13 | D | 4 | 446.2245 |
| 1738.7959 | 14 | P | 3 | 331.1976 |
| 1825.8279 | 15 | S | 2 | 234.1448 |
| | 16 | K | 1 | 147.1128 |

[Click] to move table

Static Modifications:
C: 57.02146

Variable Modifications:
S: 203.079373
