## Supplementary material for "Site-specific O-glycosylation analysis of SARS-CoV-2 spike protein produced in insect and human cells": Dataset S7: S1097-2-Y.pdf

### Fragment Ion View

Project name: 20200330COV19
Sample name: 20200416_COV19-double_2
Created date: 2020-04-18 15:22:56.0
Search name:
Activation type: CID

Peptide sequence: R.EGVFVS(203.079373)NGTHWF.V
Scan number: 52170
Calculated M+H: 1582.7173
Measured M+H: 1582.7219
Charge state: 2
File name: 20200416_COV19-double_2
search type: light

Ions:
a □1+ □2+ □3+
b ☑1+ □2+ □3+
c □1+ □2+ □3+
x □1+ □2+ □3+
y ☑1+ □2+ □3+
z □1+ □2+ □3+
[Deselect All]

Neutral Loss:
□ $H_2O$ (o)
□ $NH_3$ (*)
□ $H_3PO_4$ (^)
□ Precursor (#)
□ Precursor with reduced charges

Mass Type: ◉Mono ○Avg

Mass Tol: 20

○Dalton ◉PPM

Update

Peak Assignment:
◉Most Intense
○Nearest Match
□Peak Detect
□Precursor Removal

Peak Labels:
◉Ion ○m/z
○None

Width: 750

Height: 450

EGVFVSNGTHWF, MH+ 1582.7172, m/z 791.8623
File: 20200416_COV19-double_2, Scan: 52170, Precursor m/z: 791.8646, Charge: 2

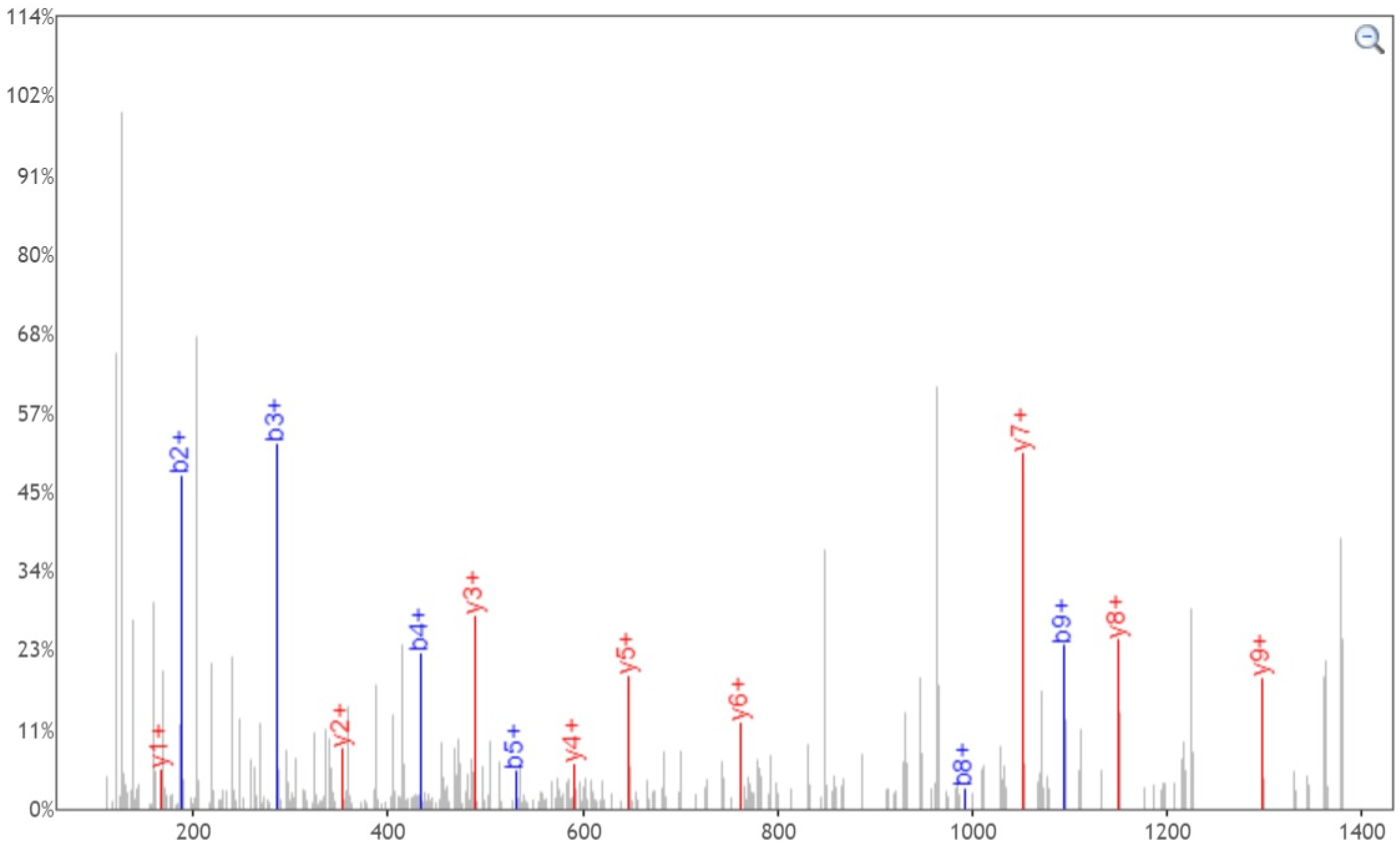


Click and drag in the plot to zoom X:☑ Y:□ Zoom Out Print □Enable tooltip

| b+ | # | Seq | # | y+ |
|---|---|---|---|---|
| 130.0499 | 1 | E | 12 | |
| 187.0713 | 2 | G | 11 | 1453.6747 |
| 286.1397 | 3 | V | 10 | 1396.6532 |
| 433.2082 | 4 | F | 9 | 1297.5848 |
| 532.2766 | 5 | V | 8 | 1150.5164 |
| 822.3880 | 6 | S | 7 | 1051.4480 |
| 936.4309 | 7 | N | 6 | 761.3365 |
| 993.4524 | 8 | G | 5 | 647.2936 |
| 1094.5000 | 9 | T | 4 | 590.2722 |
| 1231.5590 | 10 | H | 3 | 489.2245 |
| 1417.6383 | 11 | W | 2 | 352.1656 |
| | 12 | F | 1 | 166.0863 |

[Click] to move table

Static Modifications:
C: 57.02146

Variable Modifications:
S: 203.079373
