## Supplementary material for "Site-specific O-glycosylation analysis of SARS-CoV-2 spike protein produced in insect and human cells": Dataset S7: T63-3-Y.pdf

### Fragment Ion View

Project name: 20200330COV19
Sample name: 20200416_COV19-double_2
Created date: 2020-04-18 15:22:56.0
Search name:
Activation type: CID

Peptide sequence: R.SSVLHSTQDLFLPFFSNVT(203.079373)WF.H
Scan number: 56810
Calculated M+H: 2675.3027
Measured M+H: 2675.3083
Charge state: 2
File name: 20200416_COV19-double_2
search type: light

**Ions:**
a 1+ 2+ 3+
b 1+ 2+ 3+
c 1+ 2+ 3+
x 1+ 2+ 3+
y 1+ 2+ 3+
z 1+ 2+ 3+
[Deselect All]

**Neutral Loss:**
$H_2O$ (o)
$NH_3$ (*)
$H_3PO_4$ (^)
Precursor (#)
Precursor with reduced charges

Mass Type:
Mono Avg

Mass Tol: 20
Dalton PPM
Update

Peak Assignment:
Most Intense
Nearest Match
Peak Detect
Precursor Removal

Peak Labels:
Ion m/z
None

Width: 750

Height: 450

SSVLHSTQDLFLPFFSNVTWF, MH+ 2675.3028, m/z 1338.1550
File: 20200416_COV19-double_2, Scan: 56810, Precursor m/z: 1338.1578, Charge: 2

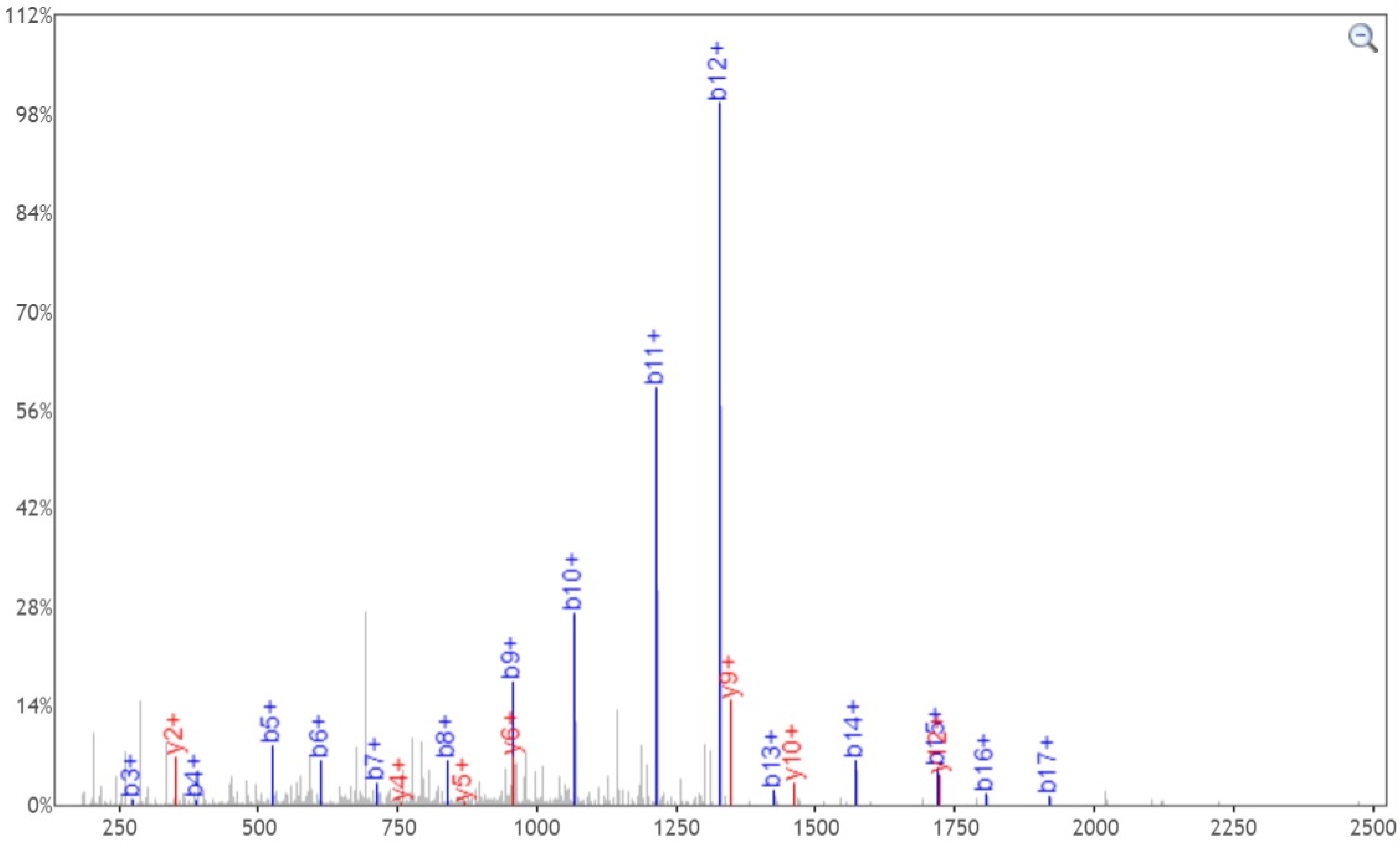


Click and drag in the plot to zoom X: Y: Zoom Out Print Enable tooltip

| b+ | # | Seq | # | y+ |
|---|---|---|---|---|
| 88.0393 | 1 | S | 21 | |
| 175.0713 | 2 | S | 20 | 2588.2708 |
| 274.1397 | 3 | V | 19 | 2501.2387 |
| 387.2238 | 4 | L | 18 | 2402.1703 |
| 524.2827 | 5 | H | 17 | 2289.0863 |
| 611.3148 | 6 | S | 16 | 2152.0274 |
| 712.3624 | 7 | T | 15 | 2064.9953 |
| 840.4210 | 8 | Q | 14 | 1963.9476 |
| 955.4480 | 9 | D | 13 | 1835.8891 |
| 1068.5320 | 10 | L | 12 | 1720.8621 |
| 1215.6004 | 11 | F | 11 | 1607.7781 |
| 1328.6845 | 12 | L | 10 | 1460.7096 |
| 1425.7373 | 13 | P | 9 | 1347.6256 |
| 1572.8057 | 14 | F | 8 | 1250.5728 |
| 1719.8741 | 15 | F | 7 | 1103.5044 |
| 1806.9061 | 16 | S | 6 | 956.4360 |
| 1920.9490 | 17 | N | 5 | 869.4040 |
| 2020.0175 | 18 | V | 4 | 755.3610 |
| 2324.1445 | 19 | T | 3 | 656.2926 |
| 2510.2238 | 20 | W | 2 | 352.1656 |
| | 21 | F | 1 | 166.0863 |

[Click] to move table

Static Modifications:
C: 57.02146

Variable Modifications:
T: 203.079373
