## Supplementary material for "Site-specific O-glycosylation analysis of SARS-CoV-2 spike protein produced in insect and human cells": Dataset S7: T76-1-N.pdf

### Fragment Ion View

Project name: 20200330COV19
Sample name: 20200416_COV19-double_2
Created date: 2020-04-18 15:22:56.0
Search name:
Activation type: CID

Peptide sequence: F.FSN(2.988261)VTWFHAIHVSGTN(2.988261)GTK.R
Scan number: 49970
Calculated M+H: 2109.017
Measured M+H: 2109.0225
Charge state: 3
File name: 20200416_COV19-double_2
search type: light

Ions:
a 1+ 2+ 3+
b 1+ 2+ 3+
c 1+ 2+ 3+
x 1+ 2+ 3+
y 1+ 2+ 3+
z 1+ 2+ 3+
[Deselect All]

Neutral Loss:
$H_2O$ (o)
$NH_3$ (*)
$H_3PO_4$ (^)
Precursor (#)
Precursor with reduced charges

Mass Type:
Mono Avg

Mass Tol: 20
Dalton PPM
Update

Peak Assignment:
Most Intense
Nearest Match
Peak Detect
Precursor Removal

Peak Labels:
Ion m/z
None

Width: 750
Height: 450

FSNVTWFHAIHVSGTNGTK, MH+ 2109.0172, m/z 703.6772
File: 20200416_COV19-double_2, Scan: 49970, Precursor m/z: 703.679, Charge: 3

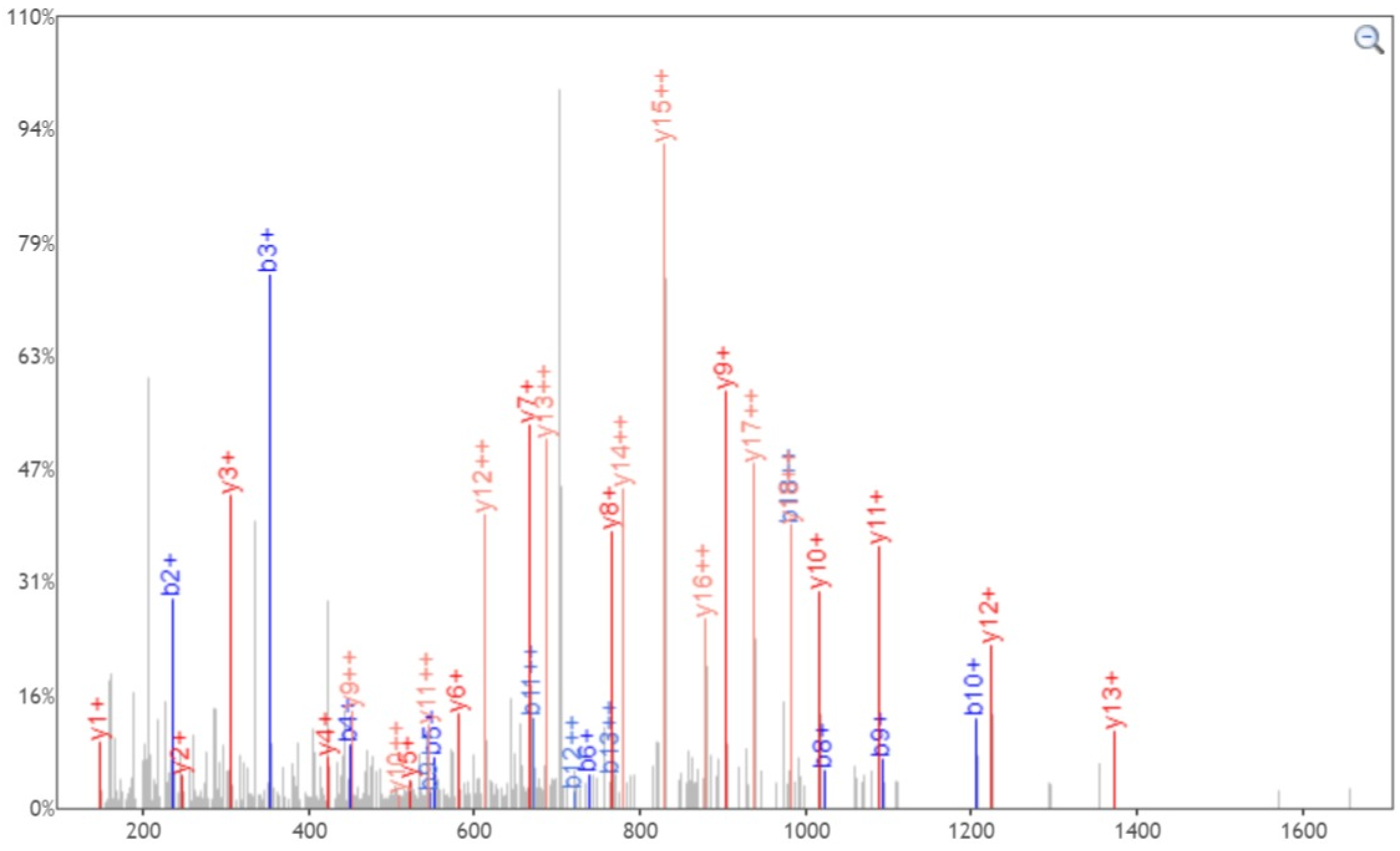


Click and drag in the plot to zoom X: Y: Zoom Out Print Enable tooltip

| b+ | b2+ | # | Seq | # | y+ | y2+ |
|---|---|---|---|---|---|---|
| 148.0757 | 74.5415 | 1 | F | 19 | | |
| 235.1077 | 118.0575 | 2 | S | 18 | 1961.9488 | 981.4780 |
| 352.1389 | 176.5731 | 3 | N | 17 | 1874.9167 | 937.9620 |
| 451.2073 | 226.1073 | 4 | V | 16 | 1757.8855 | 879.4464 |
| 552.2550 | 276.6311 | 5 | T | 15 | 1658.8171 | 829.9122 |
| 738.3343 | 369.6708 | 6 | W | 14 | 1557.7694 | 779.3884 |
| 885.4027 | 443.2050 | 7 | F | 13 | 1371.6901 | 686.3487 |
| 1022.4616 | 511.7345 | 8 | H | 12 | 1224.6217 | 612.8145 |
| 1093.4988 | 547.2530 | 9 | A | 11 | 1087.5628 | 544.2850 |
| 1206.5828 | 603.7950 | 10 | I | 10 | 1016.5257 | 508.7665 |
| 1343.6417 | 672.3245 | 11 | H | 9 | 903.4416 | 452.2245 |
| 1442.7101 | 721.8587 | 12 | V | 8 | 766.3827 | 383.6950 |
| 1529.7422 | 765.3747 | 13 | S | 7 | 667.3143 | 334.1608 |
| 1586.7636 | 793.8855 | 14 | G | 6 | 580.2823 | 290.6448 |
| 1687.8113 | 844.4093 | 15 | T | 5 | 523.2608 | 262.1340 |
| 1804.8425 | 902.9249 | 16 | N | 4 | 422.2131 | 211.6102 |
| 1861.8640 | 931.4356 | 17 | G | 3 | 305.1819 | 153.0946 |
| 1962.9116 | 981.9595 | 18 | T | 2 | 248.1605 | 124.5839 |
| | | 19 | K | 1 | 147.1128 | 74.0600 |

[Click] to move table

**Static Modifications:**
**C: 57.02146**

**Variable Modifications:**
**N: 2.988261**
