## Supplementary material for "Site-specific O-glycosylation analysis of SARS-CoV-2 spike protein produced in insect and human cells": Dataset S7: T76-1-Y.pdf

### Fragment Ion View

Project name: 20200330COV19
Sample name: 20200416_COV19-double_2
Created date: 2020-04-18 15:22:56.0
Search name:
Activation type: CID

Peptide sequence: F.FSNVT(203.079373)WFHAIHVSGTN(2.988261)GTK.R
Scan number: 47985
Calculated M+H: 2309.1084
Measured M+H: 2309.118
Charge state: 4
File name: 20200416_COV19-double_2
search type: light

Ions:
a 1+ 2+ 3+
b 1+ 2+ 3+
c 1+ 2+ 3+
x 1+ 2+ 3+
y 1+ 2+ 3+
z 1+ 2+ 3+
[Deselect All]

Neutral Loss:
$H_2O$ (o)
$NH_3$ (*)
$H_3PO_4$ (^)
Precursor (#)
Precursor with reduced charges

Mass Type:
Mono Avg

Mass Tol:
20
Dalton PPM
Update

Peak Assignment:
Most Intense
Nearest Match
Peak Detect
Precursor Removal

Peak Labels:
Ion m/z
None

Width: 750

Height: 450

FSNVTWFHAIHVSGTNGTK, MH+ 2309.1083, m/z 578.0325
File: 20200416_COV19-double_2, Scan: 47985, Precursor m/z: 578.0349, Charge: 4

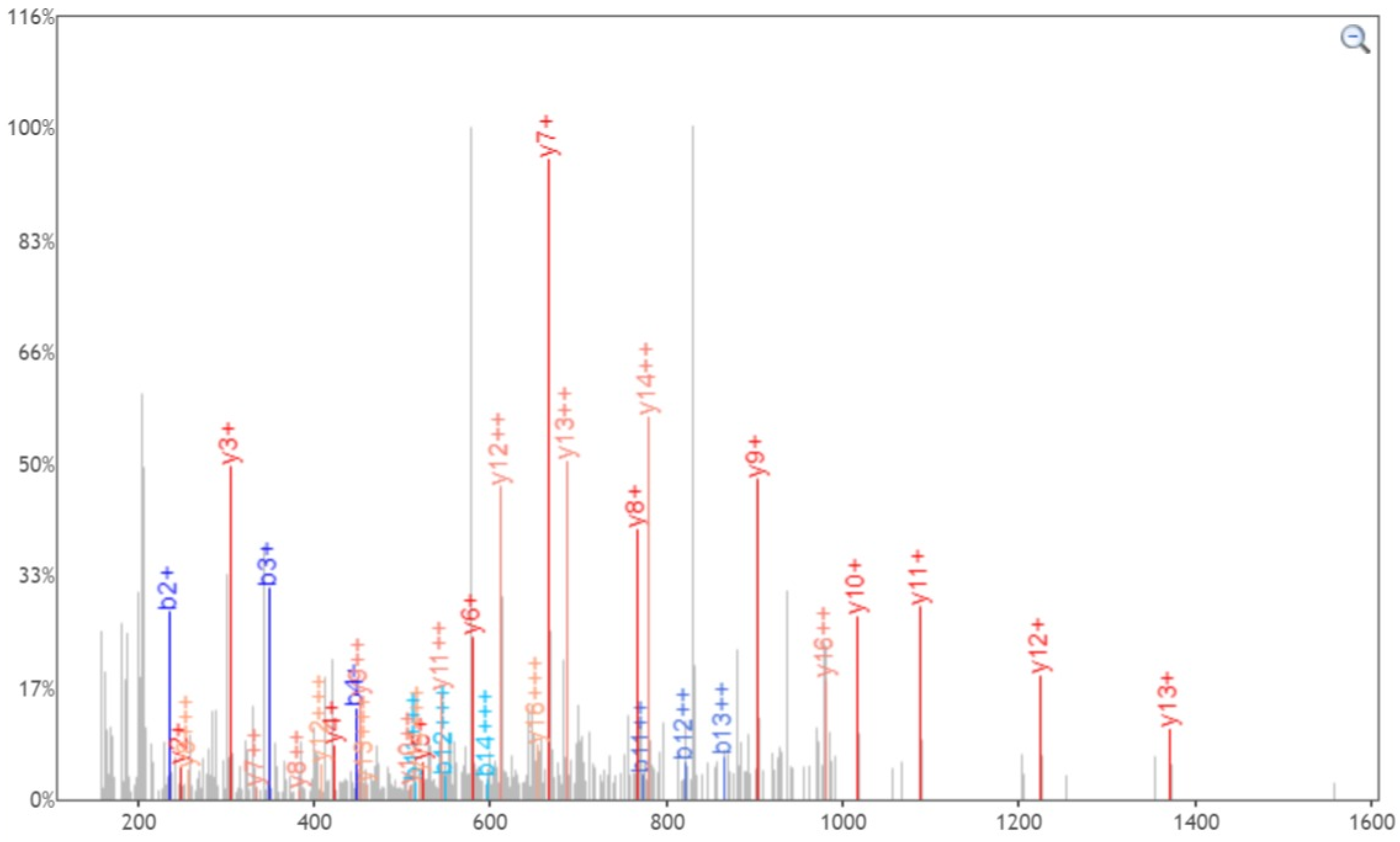


Click and drag in the plot to zoom X: Y: Zoom Out Print Enable tooltip

| b+ | b2+ | b3+ | # | Seq | # | y+ | y2+ | y3+ |
|---|---|---|---|---|---|---|---|---|
| 148.0757 | 74.5415 | 50.0301 | 1 | F | 19 | | | |
| 235.1077 | 118.0575 | 79.0408 | 2 | S | 18 | 2162.0399 | 1081.5236 | 721.3515 |
| 349.1506 | 175.0790 | 117.0551 | 3 | N | 17 | 2075.0078 | 1038.0076 | 692.3408 |
| 448.2191 | 224.6132 | 150.0779 | 4 | V | 16 | 1960.9649 | 980.9861 | 654.3265 |
| 752.3461 | 376.6767 | 251.4536 | 5 | T | 15 | 1861.8965 | 931.4519 | 621.3037 |
| 938.4254 | 469.7164 | 313.4800 | 6 | W | 14 | 1557.7694 | 779.3884 | 519.9280 |
| 1085.4938 | 543.2506 | 362.5028 | 7 | F | 13 | 1371.6901 | 686.3487 | 457.9016 |
| 1222.5528 | 611.7800 | 408.1891 | 8 | H | 12 | 1224.6217 | 612.8145 | 408.8788 |
| 1293.5899 | 647.2986 | 431.8681 | 9 | A | 11 | 1087.5628 | 544.2850 | 363.1925 |
| 1406.6739 | 703.8406 | 469.5628 | 10 | I | 10 | 1016.5257 | 508.7665 | 339.5134 |
| 1543.7328 | 772.3701 | 515.2491 | 11 | H | 9 | 903.4416 | 452.2245 | 301.8187 |
| 1642.8013 | 821.9043 | 548.2719 | 12 | V | 8 | 766.3827 | 383.6950 | 256.1324 |
| 1729.8333 | 865.4203 | 577.2826 | 13 | S | 7 | 667.3143 | 334.1608 | 223.1096 |
| 1786.8547 | 893.9310 | 596.2898 | 14 | G | 6 | 580.2823 | 290.6448 | 194.0989 |
| 1887.9024 | 944.4549 | 629.9723 | 15 | T | 5 | 523.2608 | 262.1340 | 175.0918 |
| 2004.9336 | 1002.9704 | 668.9827 | 16 | N | 4 | 422.2131 | 211.6102 | 141.4092 |
| 2061.9551 | 1031.4812 | 687.9899 | 17 | G | 3 | 305.1819 | 153.0946 | 102.3988 |
| 2163.0028 | 1082.0050 | 721.6724 | 18 | T | 2 | 248.1605 | 124.5839 | 83.3917 |
| | | | 19 | K | 1 | 147.1128 | 74.0600 | 49.7091 |

[Click] to move table

Static Modifications:
C: 57.02146

Variable Modifications:
T: 203.079373
N: 2.988261
