## Supplementary material for "Site-specific O-glycosylation analysis of SARS-CoV-2 spike protein produced in insect and human cells": Dataset S7: T76-2-N.pdf

### Fragment Ion View

Project name: 20200330COV19
Sample name: 20200416_COV19-double_2
Created date: 2020-04-18 15:22:56.0
Search name:
Activation type: CID

Peptide sequence: F.HAIHVSGTN(2.988261)GTK.R
Scan number: 7968
Calculated M+H: 1224.6217
Measured M+H: 1224.6249
Charge state: 3
File name: 20200416_COV19-double_2
search type: light

Ions:
a 1+ 2+ 3+
b 1+ 2+ 3+
c 1+ 2+ 3+
x 1+ 2+ 3+
y 1+ 2+ 3+
z 1+ 2+ 3+
[Deselect All]

Neutral Loss:
$H_2O$ (o)
$NH_3$ (*)
$H_3PO_4$ (^)
Precursor (#)
Precursor with reduced charges

Mass Type:
Mono Avg

Mass Tol: 20
Dalton PPM

Update

Peak Assignment:
Most Intense
Nearest Match
Peak Detect
Precursor Removal

Peak Labels:
Ion m/z
None

Width: 750

Height: 450

**HAIHVSGTNGTK**, MH+ 1224.6217, m/z 408.8788
File: 20200416_COV19-double_2, Scan: 7968, Precursor m/z: 408.8798, Charge: 3

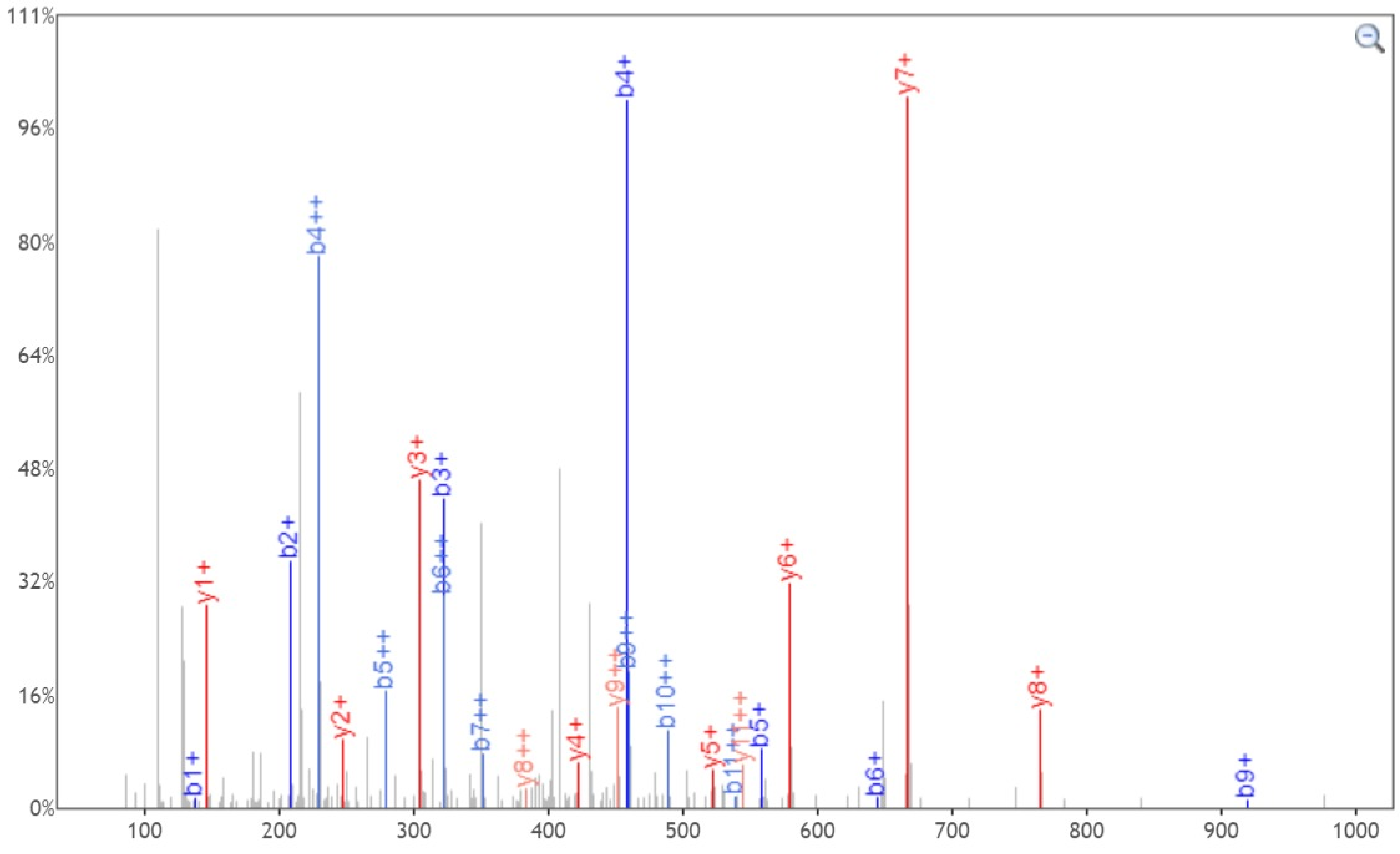


Click and drag in the plot to zoom X: Y: Zoom Out Print Enable tooltip

| b+ | b2+ | # | Seq | # | y+ | y2+ |
|---|---|---|---|---|---|---|
| 138.0662 | 69.5367 | 1 | H | 12 | | |
| 209.1033 | 105.0553 | 2 | A | 11 | 1087.5628 | 544.2850 |
| 322.1874 | 161.5973 | 3 | I | 10 | 1016.5257 | 508.7665 |
| 459.2463 | 230.1268 | 4 | H | 9 | 903.4416 | 452.2245 |
| 558.3147 | 279.6610 | 5 | V | 8 | 766.3827 | 383.6950 |
| 645.3467 | 323.1770 | 6 | S | 7 | 667.3143 | 334.1608 |
| 702.3682 | 351.6877 | 7 | G | 6 | 580.2823 | 290.6448 |
| 803.4159 | 402.2116 | 8 | T | 5 | 523.2608 | 262.1340 |
| 920.4471 | 460.7272 | 9 | N | 4 | 422.2131 | 211.6102 |
| 977.4685 | 489.2379 | 10 | G | 3 | 305.1819 | 153.0946 |
| 1078.5162 | 539.7617 | 11 | T | 2 | 248.1605 | 124.5839 |
| | | 12 | K | 1 | 147.1128 | 74.0600 |

[Click] to move table

Static Modifications:
C: 57.02146

Variable Modifications:
N: 2.988261
