## Supplementary material for "Site-specific O-glycosylation analysis of SARS-CoV-2 spike protein produced in insect and human cells": Dataset S7: T114-1-N.pdf

### Fragment Ion View

Project name: 20200330COV19
Sample name: 20200416_COV19-double_2
Created date: 2020-04-18 15:22:56.0
Search name:
Activation type: CID

Peptide sequence: K.TQSLLIVN(203.079373)NATNVVIK.V
Scan number: 49009
Calculated M+H: 1930.0856
Measured M+H: 1930.0911
Charge state: 2
File name: 20200416_COV19-double_2
search type: light

**Ions:**
c 1+ 2+ 3+
x 1+ 2+ 3+
y [x] 1+ 2+ 3+
z 1+ 2+ 3+
[Deselect All]

**Neutral Loss:**
- $H_2O$ (o)
- $NH_3$ (*)
- $H_3PO_4$ (^)
- Precursor (#)
- Precursor with reduced charges

Mass Type: (•) Mono ( ) Avg

Mass Tol: 20
( ) Dalton (•) PPM

Update

Peak Assignment:
(•) Most Intense
( ) Nearest Match
Peak Detect
Precursor Removal

Peak Labels:
(•) Ion ( ) m/z
( ) None

TQSLLIVNNATNVVIK, MH+ 1930.0855, m/z 965.5464

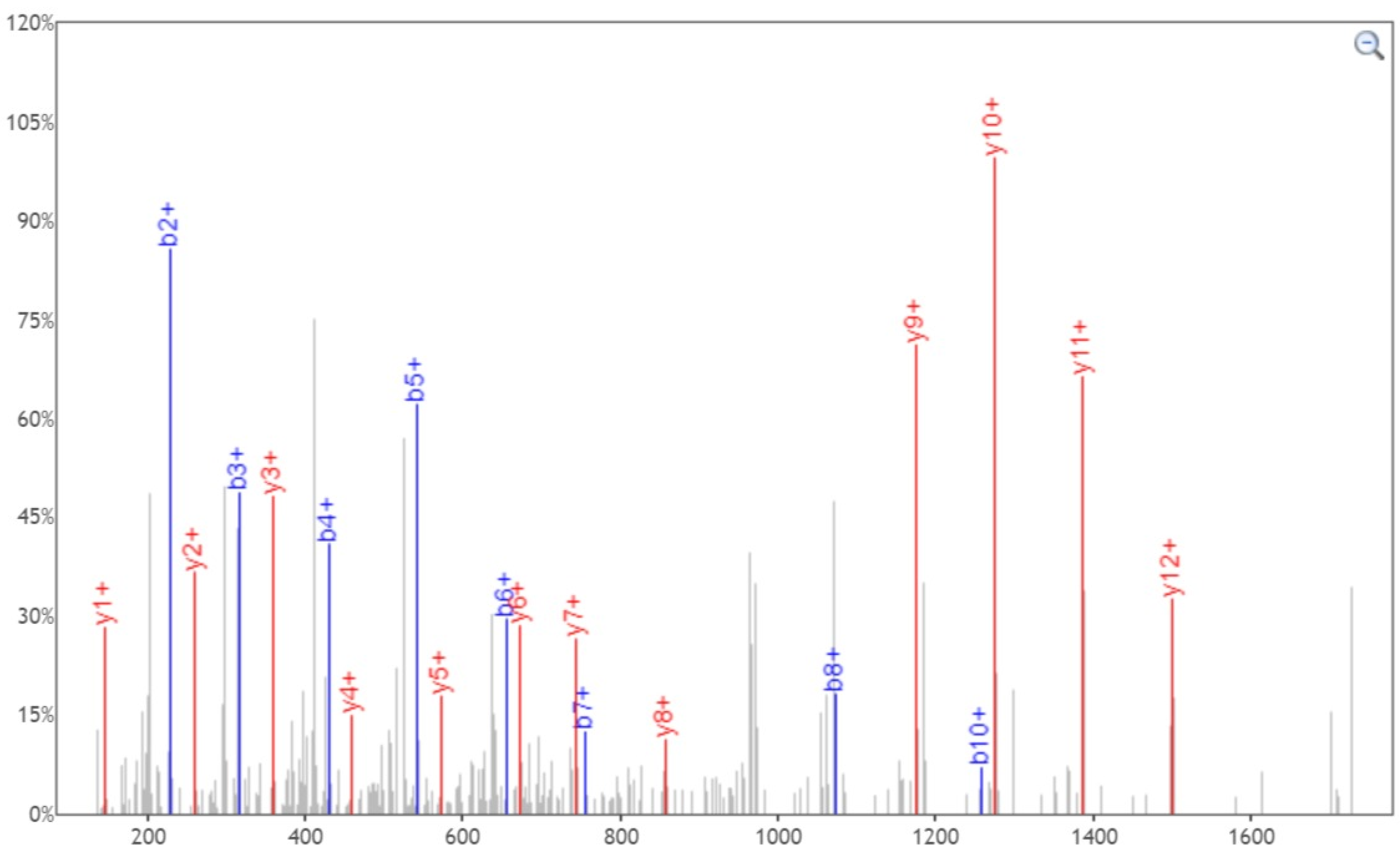


| b+ | # | Seq | # | y+ |
|---|---|---|---|---|
| 102.0550 | 1 | T | 16 | |
| 317.1456 | 3 | S | 14 | 1700.9793 |
| 430.2296 | 4 | L | 13 | 1613.9472 |
| 543.3137 | 5 | L | 12 | 1500.8632 |
| 656.3978 | 6 | I | 11 | 1387.7791 |
| 755.4662 | 7 | V | 10 | 1274.6951 |
| 1072.5885 | 8 | N | 9 | 1175.6266 |
| 1186.6314 | 9 | N | 8 | 858.5043 |
| 1257.6685 | 10 | A | 7 | 744.4614 |
| 1358.7162 | 11 | T | 6 | 673.4243 |
| 1472.7591 | 12 | N | 5 | 572.3766 |
| 1571.8275 | 13 | V | 4 | 458.3337 |
| 1670.8959 | 14 | V | 3 | 359.2653 |
| 1783.9800 | 15 | I | 2 | 260.1969 |
| | 16 | K | 1 | 147.1128 |

[Click] to move table

Static Modifications:
C: 57.02146

Variable Modifications:
N: 203.079373
