## Supplementary material for "Site-specific O-glycosylation analysis of SARS-CoV-2 spike protein produced in insect and human cells": Dataset S7: T114-1-Y.pdf

### Fragment Ion View

Project name: 20200330COV19
Sample name: 20200416_COV19-double_2
Created date: 2020-04-18 15:22:56.0
Search name:
Activation type: CID

Peptide sequence: K.T(203.079373)QSLLIVNNATNVVIK.V
Scan number: 48968
Calculated M+H: 1930.0856
Measured M+H: 1930.0898
Charge state: 3
File name: 20200416_COV19-double_2
search type: light

Ions:
a 1+ 2+ 3+
b 1+ 2+ 3+
c 1+ 2+ 3+
x 1+ 2+ 3+
y 1+ 2+ 3+
z 1+ 2+ 3+
[Deselect All]

Neutral Loss:
$H_2O$ (o)
$NH_3$ (*)
$H_3PO_4$ (^)
Precursor (#)
Precursor with reduced charges

Mass Type:
Mono Avg

Mass Tol: 20
Dalton PPM

Update

Peak Assignment:
Most Intense
Nearest Match
Peak Detect
Precursor Removal

Peak Labels:
Ion m/z
None

Width: 750

Height: 450

TQSLLIVNNATNVVIK, MH+ 1930.0855, m/z 644.0334
File: 20200416_COV19-double_2, Scan: 48968, Precursor m/z: 644.0348, Charge: 3

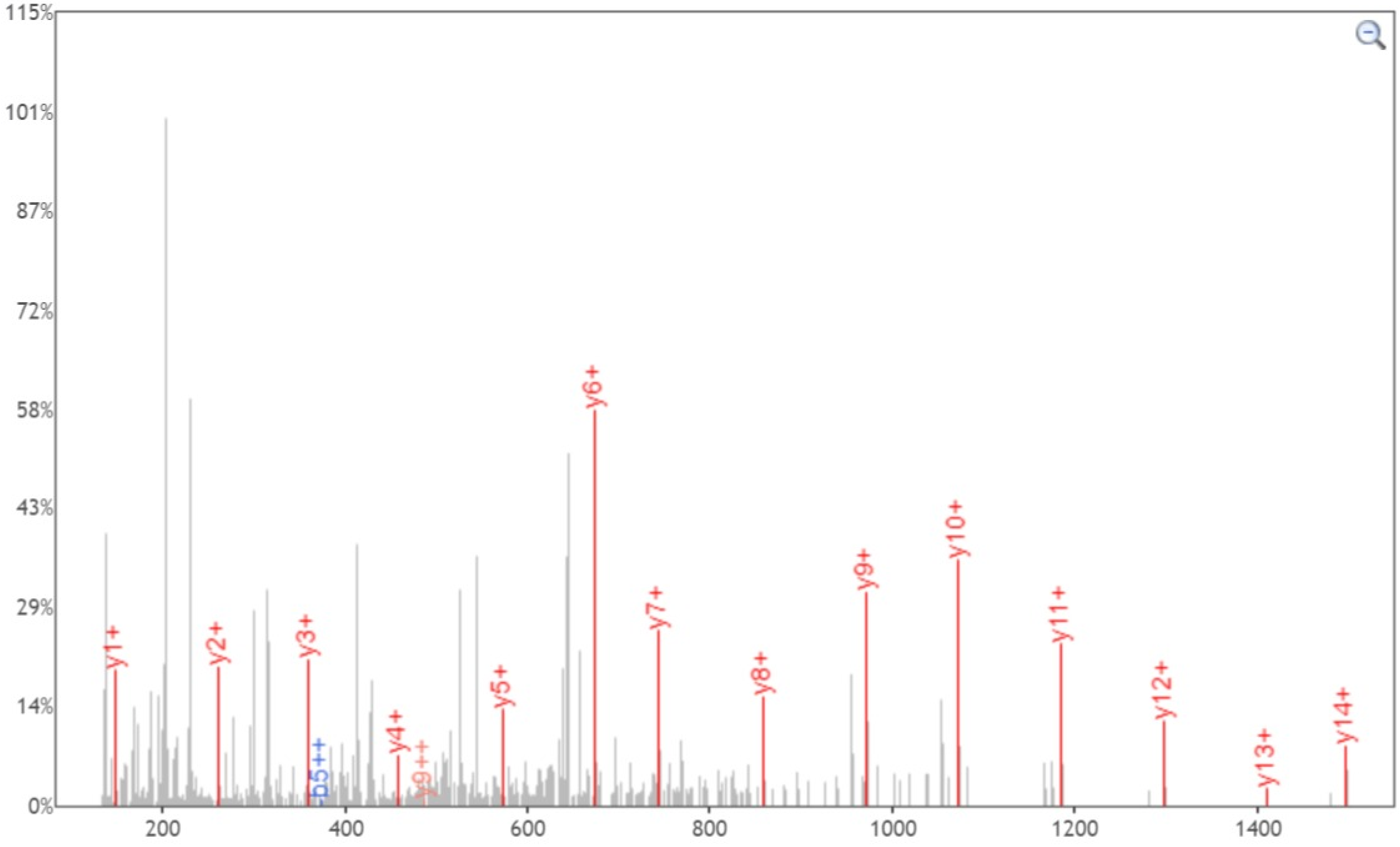


Click and drag in the plot to zoom X: Y: Zoom Out Print Enable tooltip

| b+ | b2+ | # | Seq | # | y+ | y2+ |
|---|---|---|---|---|---|---|
| 305.1343 | 153.0708 | 1 | T | 16 | | |
| 433.1929 | 217.1001 | 2 | Q | 15 | 1625.9585 | 813.4829 |
| 520.2249 | 260.6161 | 3 | S | 14 | 1497.8999 | 749.4536 |
| 633.3090 | 317.1581 | 4 | L | 13 | 1410.8679 | 705.9376 |
| 746.3931 | 373.7002 | 5 | L | 12 | 1297.7838 | 649.3955 |
| 859.4771 | 430.2422 | 6 | I | 11 | 1184.6997 | 592.8535 |
| 958.5455 | 479.7764 | 7 | V | 10 | 1071.6157 | 536.3115 |
| 1072.5885 | 536.7979 | 8 | N | 9 | 972.5473 | 486.7773 |
| 1186.6314 | 593.8193 | 9 | N | 8 | 858.5043 | 429.7558 |
| 1257.6685 | 629.3379 | 10 | A | 7 | 744.4614 | 372.7343 |
| 1358.7162 | 679.8617 | 11 | T | 6 | 673.4243 | 337.2158 |
| 1472.7591 | 736.8832 | 12 | N | 5 | 572.3766 | 286.6919 |
| 1571.8275 | 786.4174 | 13 | V | 4 | 458.3337 | 229.6705 |
| 1670.8959 | 835.9516 | 14 | V | 3 | 359.2653 | 180.1363 |
| 1783.9800 | 892.4936 | 15 | I | 2 | 260.1969 | 130.6021 |
| | | 16 | K | 1 | 147.1128 | 74.0600 |

[Click] to move table

Static Modifications:
**C: 57.02146**

Variable Modifications:
**T: 203.079373**
